# A high-quality reference genome for the fission yeast *Schizosaccharomyces osmophilus*

**DOI:** 10.1101/2022.12.06.519325

**Authors:** Guo-Song Jia, Wen-Cai Zhang, Yue Liang, Xi-Han Liu, Nicholas Rhind, Alison Pidoux, Michael Brysch-Herzberg, Li-Lin Du

**Affiliations:** National Institute of Biological Sciences, Beijing 102206, China; Department of Biochemistry and Molecular Biotechnology, University of Massachusetts Medical School, Worcester, MA 01605, USA; Wellcome Centre for Cell Biology, Institute of Cell Biology, School of Biological Sciences, The University of Edinburgh, Edinburgh EH9 3BF, Scotland, UK; Laboratory for Wine Microbiology, Department International Business, Heilbronn University, 74081 Heilbronn, Germany; Tsinghua Institute of Multidisciplinary Biomedical Research, Tsinghua University, Beijing 102206, China

**Keywords:** fission yeast, *Schizosaccharomyces osmophilus*, centromere, telomere, retrotransposon, double-hairpin element

## Abstract

Fission yeasts are an ancient group of fungal species that diverged from each other from tens to hundreds of million years ago. Among them is the preeminent model organism *Schizosaccharomyces pombe*, which has significantly contributed to our understandings of molecular mechanisms underlying fundamental cellular processes. The availability of the genomes of *S. pombe* and three other fission yeast species *S. japonicus*, *S. octosporus*, and *S. cryophilus* has enabled cross-species comparisons that provide insights into the evolution of genes, pathways, and genomes. Here, we performed genome sequencing on the type strain of the recently identified fission yeast species *S. osmophilus* and obtained a complete mitochondrial genome and a nuclear genome assembly with gaps only at rRNA gene arrays. A total of 5098 protein-coding nuclear genes were annotated and orthologs for more than 95% of them were identified. Genome-based phylogenetic analysis showed that *S. osmophilus* is most closely related to *S. octosporus* and these two species diverged around 16 million years ago. To demonstrate the utility of this *S. osmophilus* reference genome, we conducted cross-species comparative analyses of centromeres, telomeres, transposons, the mating-type region, Cbp1 family proteins, and mitochondrial genomes. These analyses revealed conservation of repeat arrangements and sequence motifs in centromere cores, identified telomeric sequences composed of two types of repeats, delineated relationships among Tf1/sushi group retrotransposons, characterized the evolutionary origins and trajectories of Cbp1 family domesticated transposases, and discovered signs of interspecific transfer of two types of mitochondrial selfish elements.

## Introduction

Fission yeasts (genus *Schizosaccharomyces*) are an ancient group of fungal species that originated more than 200 million years ago (Rhind *et al*. 2011; Shen *et al*. 2020). They belong to the *Taphrinomycotina* subphylum of *Ascomycota* (Liu *et al*. 2009). There are five currently recognized species of fission yeasts: *S. pombe*, *S. japonicus*, *S. octosporus*, *S. cryophilus*, and *S. osmophilus* (Helston *et al*. 2010; Vaughan-Martini and Martini 2011; Brysch-Herzberg *et al*. 2019). *S. pombe* was adopted for biological research more than 70 years ago (Leupold 1950; Fantes and Hoffman 2016), and is one of the most prominent model organisms (Hoffman *et al*. 2015; Hayles and Nurse 2018; Vyas *et al*. 2021). In the last decade, *S. japonicus* has emerged as a powerful new experimental model for biologists, especially those interested in evolutionary cell biology (Klar 2013; Aoki *et al*. 2017; Rutherford *et al*. 2022). Genetic tools have also been developed for *S. octosporus* (Seike and Niki 2017), adding another fission yeast species to the list of experimentally amenable organisms.

The value of fission yeast species as research subjects relies to a great extent on the availability of well-annotated genomes. *S. pombe* is the sixth eukaryote to have its genome fully sequenced (Wood *et al*. 2002), and has one of the most completely annotated genomes (Harris *et al*. 2022). The genomes of *S. japonicus*, *S. octosporus*, and *S. cryophilus* were published in 2011 (Rhind *et al*. 2011). More recently, long-read sequencing has been used to generate more contiguous genome assemblies of *S. octosporus* and *S. cryophilus* (Tong *et al*. 2019). These genomes have not only served as critical resources for molecular studies using these species, but also enabled cross-species comparative genomic analyses, which have shed light on how centromere structure, transposon diversity, the roles of RNAi, and carbon metabolism have evolved in the past tens to hundreds of million years (Rhind *et al*. 2011; Tong *et al*. 2019).

*S. osmophilus* is the newest fission yeast species to be identified. It was initially discovered in bee bread of solitary bee species belonging to the *Megachilinae* subfamily (Brysch-Herzberg *et al*. 2019). Bee bread, which is a mixture of pollen and nectar, is the food source for growing bee larvae (Figure 1A). An extensive fission yeast isolation study revealed that bee bread of solitary bees is the only type of habitat where *S. osmophilus* can be frequently isolated (Brysch-Herzberg *et al*. 2022). Like other fission yeast species, *S. osmophilus* cells propagate vegetatively by medial fission (Figure 1B). During sexual reproduction, *S. osmophilus* usually form asci containing eight spores (Figure 1C). Analyses of the sequences of rRNA genes and the *rrk1* gene showed that *S. osmophilus* is more closely related to *S. cryophilus* and *S. octosporus* than to *S. pombe* or *S. japonicus* (Brysch-Herzberg *et al*. 2019). Unique among fission yeast species, *S. osmophilus* is an obligate osmophilic organism that requires a high-osmolarity environment for growth, likely due to its long-term evolution in high-osmolarity habitats such as bee bread, nectar, and honey (Brysch-Herzberg *et al*. 2019).

**Figure 1.**
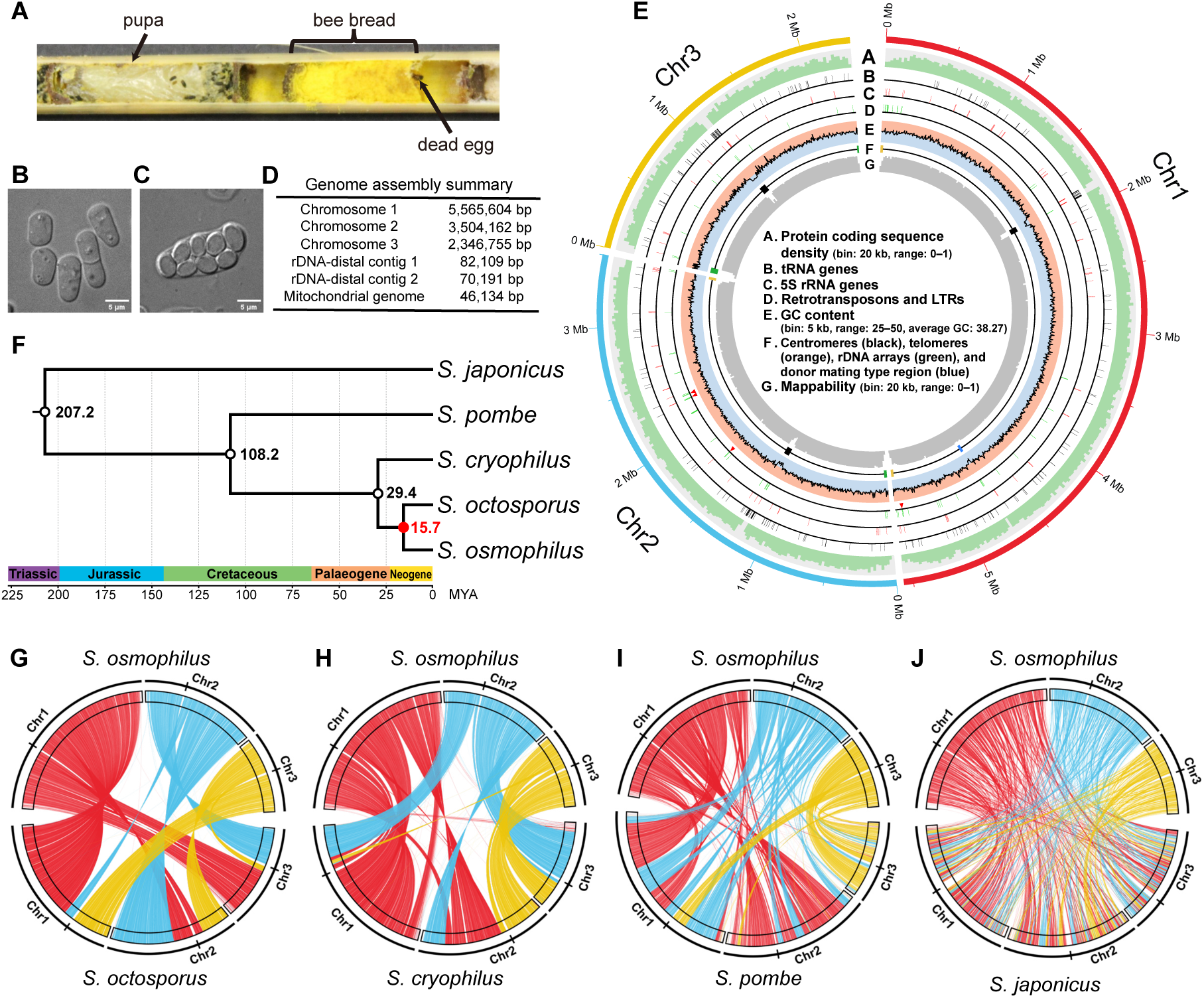
The genome assembly of the *S. osmophilus* type strain CBS 15793^T^. (A) A cross-section photo of a nesting tube for solitary bees. Two nest chambers can be seen in the photo. A pupa occupies the left chamber, where little bee bread is remaining. In the right chamber, a dead egg is visible and the chamber is full of bee bread. Bee bread of solitary bees is the type of substrate from which *S. osmophilus* has most frequently been isolated. (B) An image of vegetative cells of the *S. osmophilus* type strain CBS 15793^T^ grown in liquid YES medium containing 30% glucose at 25°C. (C) An image of an ascus of the *S. osmophilus* type strain CBS 15793^T^ undergoing sporulation in liquid PMG medium containing 30% glucose at 25°C. (D) The sizes of the contigs in the genome assembly of the *S. osmophilus* type strain CBS 15793^T^. (E) Circos diagram illustrating the features of the three chromosomal contigs. From outer to inner: protein coding sequence density (A, bin size = 20 kb, range = 0–1); tRNA genes (B); 5S rRNA genes (C); full-length and intact retrotransposons (red arrowheads) and long terminal repeats (LTRs) (green lines) (D); GC content (E, bin size = 5 kb, range = 25%–50%, the boundary between the blue and the red backgrounds represents the genome-wide average GC content, 38.27%); centromeres (black), telomeres (orange, not drawn to scale), rDNA arrays (green), and the donor mating type region (blue) (E); mappability calculated by running GenMap using 100 bp k-mers with up to two mismatches (Pockrandt *et al*. 2020) (G). (F) Time-calibrated phylogeny of fission yeast species. This timetree is based on a maximum likelihood tree inferred using a concatenation super-matrix of 1060 “complete and single-copy” BUSCO genes present in five fission yeast species and the outgroup species *S. complicate* (Figure S5A). The three time calibration nodes are shown as empty black circles. The divergence time between *S. octosporus* and *S. osmophilus* was calculated using RelTime and the corresponding node is shown as a filled red circle. (G–J) Plots showing synteny conservation between the nuclear genome of *S. osmophilus* (top of each plot) and the nuclear genomes of the other four fission yeast species (bottom of each plot). The orientations of the chromosomes of *S. osmophilus* are clockwise and the orientations of the chromosomes of other species are counterclockwise. The midpoints of centromeres are indicated by vertical black lines on the outer ring. Colored lines connect single-copy orthologous genes of the two species shown in each plot. The color of a line is based on which chromosome the *S. osmophilus* gene is on (chromosome 1, red; chromosome 2, blue; chromosome 3, yellow). The *S. octosporus* and *S. cryophilus* genomes used in this analysis are those reported in (Tong *et al*. 2019). The *S. pombe* and *S. japonicus* genomes used in this analysis are the reference genomes.

In this study, we generated a high quality genome assembly for the type strain of *S. osmophilus* and comprehensively annotated the genes in this genome assembly. We performed a series of comparative analyses using this reference genome of *S. osmophilus* and obtained new insights into the evolution of centromeres, telomeres, retrotransposons, Cbp1 family proteins, and mitochondrial genomes in the fission yeast lineage.

## Materials and Methods

### Strain and media

The strain used for genome sequencing in this study is the *S. osmophilus* holotype strain CBS 15793^T^ (isotype: CLIB 3267^T^ = NCAIM Y.02225^T^; MycoBank #: 829586; original name SZ134-FG-A) (Brysch-Herzberg *et al*. 2019). *S. osmophilus* cells were cultured in a modified Yeast Extract Supplemented (YES) medium (30% glucose, 0.5% yeast extract, 200 mg/L of leucine, adenine, uracil, and histidine) for vegetative growth and in a modified Pombe Minimal Glutamate (PMG) medium (30% glucose, 3 g/L potassium hydrogen phthalate, 2.2 g/L sodium hydrogen phosphate anhydrous, 5 g/L L-glutamate, standard concentrations of salts, vitamins, and minerals) for sporulation (Forsburg and Rhind 2006).

### Genome sequencing, assembly, and quality assessment

*S. osmophilus* cells were grown in the modified YES medium at 25°C and cells in the logarithmic phase of vegetative growth were collected and frozen using liquid nitrogen. For long-read sequencing library construction, frozen cells were sent to Frasergen Bioinformatics Co., Ltd. (Wuhan, China) for genomic DNA extraction, sequencing library construction, and long-read sequencing on the Sequel II platform in the circular consensus sequencing (CCS) mode. A total of 430.07 Mb of PacBio HiFi read data was generated using CCS (v4.0, https://github.com/PacificBiosciences/ccs). The HiFi reads have been deposited at NCBI SRA under the accession number SRR21149833.

For short-read sequencing library construction, genomic DNA was extracted using the MasterPure Yeast DNA Purification Kit (Epicentre). An Illumina sequencing library was prepared using home-made Tn5 transposase as previously described (Tao *et al*. 2019). Paired-end sequencing was performed on the Illumina NovaSeq 6000 System (2×150 bp read pairs) by Novogene Co., Ltd. (Beijing, China), and a total of 4.86 Gb of raw read data was generated.

Low-quality reads were filtered or trimmed using fastp (v0.20.0, https://github.com/OpenGene/fastp) and 4.67 Gb of high-quality paired-end read data was retained (Chen *et al*. 2018). The short-read genomic DNA sequencing data has been deposited at NCBI SRA under the accession number SRR21149832.

PacBio HiFi reads were used to assemble the mitochondrial genome (mitogenome) using a procedure based on the mitoVGP pipeline (Formenti *et al*. 2021). Briefly, we used pbmm2 (v1.4.0, https://github.com/PacificBiosciences/pbmm2) to map the HiFi reads to a reference, which is a collection of fungal mitogenomes (https://github.com/Kinggerm/GetOrganelleDB/blob/master/0.0.1/SeedDatabase/fungus_mt.fasta) supplemented with the *S. cryophilus* mitogenome sequence (Jin *et al*. 2020). Reads mapped to the reference were given to the assembler canu (v2.2, https://github.com/marbl/canu) to perform de novo assembly (Nurk *et al*. 2020). The mitogenome contig in the assembly was identified by BLASTN search. Overlapping sequences at the ends of the mitogenome contig were identified by self-to-self BLASTN analysis and trimmed accordingly. One round of polishing with Illumina short reads was performed using Pilon v1.23 (Walker *et al*. 2014). The annotation of the mitogenome is described in a later section. The sequence and annotation of the *S. osmophilus* mitogenome have been deposited at GenBank under the accession number OP310968.

For the assembly of the nuclear genome using PacBio HiFi reads, we ran the assembler hifiasm (v0.16.1, https://github.com/chhylp123/hifiasm) under the “inbred/homozygous” mode (- l0) and only the primary assembly (named p_ctg by hifiasm) was retained (Cheng *et al*. 2021). To optimize the number of input HiFi reads, down-sampling was performed using rasusa (v0.3.0, https://github.com/mbhall88/rasusa) with the estimated genome size set to 12 Mb (Hall 2022). After several preliminary assembly trials, about 30× coverage of HiFi reads were finally used to generate the initial genome assembly. The initial genome assembly was polished once using HiFi reads and racon (v1.4.20, https://github.com/isovic/racon) (Vaser *et al*. 2017) and then polished once using Illumina paired-end reads and pilon (v1.24, https://github.com/broadinstitute/pilon) (Walker *et al*. 2014). The polished genome assembly contains 124 contigs and has a total length of 15,485,841 bp. Manual inspection of all 124 contigs assembled by hifiasm identified three megabase-sized contigs as chromosomal contigs and they were named as chromosomes 1 to 3 from largest to smallest. Among the remaining kilobase-sized contigs, 114 contigs could be aligned to the mitogenome by BLASTN and were discarded. The other seven contigs contained repeats of 18S–5.8S–25S rRNA genes (rDNA). Two of them also contained telomeric repeats. These two contigs were retained as rDNA-distal contigs and the other five were discarded. The final assembly includes three chromosomal contigs (chromosome 1: 5,565,604 bp; chromosome 2: 3,504,162 bp; chromosome 3: 2,346,755 bp), two rDNA-distal contigs (rDNA-distal contig 1: 82,109 bp; rDNA-distal contig 2: 70,191 bp), and one mitogenome contig (46,134 bp). The nuclear genome assembly together with gene annotations (see below) has been submitted to GenBank under the submission number SUB1225547.

To assess the quality of the genome assembly, raw HiFi reads and fastp-filtered Illumina paired-end reads were mapped to the genome assembly using minimap2 (v2.21, https://github.com/lh3/minimap2) (Li 2018) and BWA (v 0.7.17-r1188, https://github.com/lh3/bwa) (Li and Durbin 2009), respectively. The overall mapping ratio was calculated using GATK CollectAlignmentSummaryMetrics (v4.2.0.0, https://gatk.broadinstitute.org/hc/en-us) (McKenna *et al*. 2010). Average read depths in 10-kb sliding windows along the genome were calculated. Variant calling based on the mapping result of Illumina paired-end reads was performed using three variant callers: bcftools (v1.1.1, http://www.htslib.org/doc/1.0/bcftools.html) (Li 2011), GATK (v4.2.0.0), and deepvariant (v1.1.0, https://github.com/google/deepvariant) (Poplin *et al*. 2018). Homozygous variants that were called by at least two variant callers and passed a read depth cutoff (DP>10) were retained. Variant calling based on the mapping result of HiFi reads was performed using deepvariant (v1.1.0, --model_type PACBIO) and homozygous variants that passed both a read depth cutoff (DP>10) and a genotype quality cutoff (GQ>30) were retained. Benchmarking Universal Single-Copy Orthologs (BUSCO, v3.0.2, https://busco.ezlab.org/) was used to assess the assembly completeness based on the presence/absence of 1315 predefined single-copy orthologs of *Ascomycota* (ascomycota_odb9 gene set) (Simão *et al*. 2015).

To compare the BUSCO completeness of our *S. osmophilus* assembly to those of previously published genome assemblies of other fission yeast species, six genome assemblies were downloaded and analyzed. The reference genome of *S. pombe* (ASM294v2.29) was downloaded from PomBase (https://www.pombase.org/) (Harris *et al*. 2022). The reference genomes of *S. japonicus*, *S. octosporus*, and *S. cryophilus* were downloaded from Ensembl Fungi (https://fungi.ensembl.org/) (Rhind *et al*. 2011). More recently published PacBio sequencing-based genome assemblies of *S. octosporus* and *S. cryophilus* were downloaded from http://bifx-core.bio.ed.ac.uk/~ptong/genome_assembly/ (Tong *et al*. 2019). These six genome assemblies were analyzed using BUSCO as described above.

### Obtaining high-confidence transcript sequences

*S. osmophilus* cells were grown in the modified YES medium at 25°C and cells in the logarithmic phase of vegetative growth were collected for total RNA extraction. Cells were washed once in DEPC water at 4°C and subsequently resuspended with TES buffer (10 mM Tris pH 7.5, 10 mM EDTA pH 8.0, 0.5% SDS). One volume of acidic phenol-chloroform (1:1) was added and incubated at 65°C for 1 hour. Then the mixture was centrifuged at 4°C, and the aqueous phase was collected and treated with phenol-chloroform (1:1) and chloroform:isoamyl alcohol (24:1) sequentially. 1/10 volume of 3 M NaAc (pH 5.2) and 2 volumes of isopropanol were added to the aqueous phase, mixed thoroughly by inverting, and stored at -20°C overnight before centrifugation at 4°C. After centrifuging, the supernatant was removed, and the RNA pellet was washed with 70% ethanol twice. The RNA pellet was dissolved in DEPC water after air-drying and stored at -80°C.

The total RNA was sent to Annoroad Gene Technology (Beijing, China) for RNA-seq library construction and paired-end Illumina sequencing. A total of 2.34 Gb of raw read data was obtained. Reads were filtered or trimmed using fastp (v0.20.0) with default parameters. Cleaned reads were mapped to the *S. osmophilus* genome using STAR (v2.6.0a, https://github.com/alexdobin/STAR) with the following parameters: --alignIntronMin 29 -- alignIntronMax 819 --outFilterMultimapNmax 1 --outFilterMismatchNmax 0 --alignEndsType EndToEnd (Dobin *et al*. 2013). The Illumina paired-end RNA sequencing data has been deposited at NCBI SRA under the accession number SRR21149830.

For long-read cDNA sequencing using the Oxford Nanopore Technologies (ONT) platform, the total RNA obtained above was sent to Biomarker Technologies (Qingdao, China) for sequencing library preparation and ONT cDNA sequencing. A total of 2.15 Gb of raw read data was obtained. Reads were processed using pychopper (https://github.com/epi2me-labs/pychopper) and 1.65 Gb of full-length read data was obtained. The Full-Length Alternative Isoform analysis of RNA (FLAIR, v1.5, https://github.com/BrooksLabUCSC/flair) pipeline was used for downstream analysis (Tang *et al*. 2020). FLAIR is a pipeline designed to perform reads mapping, reads correcting, and isoform clustering for noisy long reads generated by ONT cDNA sequencing. It can also be run optionally with short-read RNA sequencing data to help increase the accuracy of splicing site identification. Briefly, the long reads were mapped to the *S. osmophilus* genome using “flair.py align” submodule with default parameters. The splicing junction information generated by short-read RNA sequencing was extracted using a FLAIR script “junctions_from_sam.py” from the reads mapping SAM file and then submitted to “flair.py correct” submodule. Finally, high-confidence transcript sequences were obtained by running a FLAIR submodule “flair.py collapse”. The ONT long-read cDNA sequencing data has been deposited at NCBI SRA under the accession number SRR21149831.

### Annotation of protein-coding genes

Protein-coding genes in the *S. osmophilus* nuclear genome were annotated using the MAKER pipeline (v3.01.04, https://www.yandell-lab.org/software/maker.html) (Campbell *et al*. 2014). The MAKER pipeline utilizes the protein evidence, the EST evidence, and ab initio gene predictions for gene annotation. For the protein evidence, protein sequences encoded in the reference genomes of *S. pombe*, *S. octosporus*, *S. cryophilus*, and *S. japonicus* were downloaded from Ensembl Fungi (https://fungi.ensembl.org/) and combined into a single FASTA file. For the EST evidence, high-confidence transcript sequences obtained as described above were used.

Two ab initio gene predictors were used: SNAP (v2006-07-28, https://github.com/KorfLab/SNAP) (Korf 2004) and AUGUSTUS (v3.2.3, https://github.com/Gaius-Augustus/Augustus) (Stanke *et al*. 2008), both of which needed to be trained. Because the training of SNAP and AUGUSTUS requires pre-existing gene models as training data, the first round of MAKER annotation was carried out and a genome annotation was generated using only the protein evidence and the EST evidence (the protein2genome, est2genome, and correct_est_fusion options in the maker_opts.ctl control file were set to 1). The resulting gene models were utilized for the training of SNAP and AUGUSTUS. After SNAP and AUGUSTUS were trained, their predictions were used in the second round of MAKER annotation. In this round, the protein2genome and est2genome options were set to 0. The resulting gene models were used to train SNAP and AUGUSTUS again. The third round of MAKER annotation was conducted with the same settings as the second round and the resulting gene models were used to re-train SNAP and AUGUSTUS. Finally, the last round of MAKER annotation was conducted with the protein2genome, est2genome, and correct_est_fusion options set to 1. All resulting gene models were retained (the “keep_preds” option was set to 1) and recorded in a GFF3 format file for downstream analysis.

To further improve the quality of gene models, EVidenceModeler (EVM, v1.1.1, https://evidencemodeler.github.io/) was employed to generate weighted consensus gene predictions (Haas *et al*. 2008). We adopted the weight values used in a published protocol for annotating the genomes of *S. cerevisiae* isolates (https://github.com/yjx1217/LRSDAY) (Yue and Liti 2018). Finally, all resulting EVM gene models were loaded into the Integrative Genomics Viewer (IGV, v2.9.2, https://software.broadinstitute.org/software/igv/) (Robinson *et al*. 2011) together with protein evidence alignments extracted from the MAKER output and ONT cDNA-seq read alignments for visual inspection. Erroneously fused genes were manually corrected and unannotated genes suggested by orthogroup analysis (see blow) were manually annotated. Systematic names containing a prefix of SOMG (for *<u>S</u>chizosaccharomyces <u>o</u>s<u>m</u>ophilus* <u>g</u>enes) followed by a five-digit number were assigned to annotated genes. The annotation and naming of *wtf* genes are described in a later section. The annotations were recorded in a GFF3 format file.

### Orthogroup analysis

Orthologous gene groups (orthogroups, OGs) of nuclear-encoded protein-coding genes across *S. pombe*, *S. octosporus*, *S. cryophilus*, *S. osmophilus*, and *S. japonicus* were identified using proteinortho (v6.1.0, https://gitlab.com/paulklemm_PHD/proteinortho) (Lechner *et al*. 2011). The protein sequences of *S. pombe* and *S. japonicus* were retrieved from PomBase and JaponicusDB (*S. pombe*: https://www.pombase.org/data/genome_sequence_and_features/feature_sequences/peptide.fa.gz; *S. japonicus*: https://www.japonicusdb.org/data/genome_sequence_and_features/feature_sequences/peptide.fa.gz). The protein sequences of *S. octosporus* and *S. cryophilus* were retrieved from Ensembl Fungi (*S. octosporus*: http://ftp.ensemblgenomes.org/pub/fungi/release-54/fasta/schizosaccharomyces_octosporus/pep/Schizosaccharomyces_octosporus.GCA_000150505.2.pep.all.fa.gz; *S. cryophilus*: http://ftp.ensemblgenomes.org/pub/fungi/release-54/fasta/schizosaccharomyces_cryophilus/pep/Schizosaccharomyces_cryophilus.GCA_000004155.2.pep.all.fa.gz). To enhance the accuracy of orthogroup identification, we enabled the POFF extension of proteinortho, which takes into account the relative gene order (synteny) (Lechner *et al*. 2014). The synteny information for POFF was generated from GFF3 files using custom scripts. The GFF3 files of *S. pombe* and *S. japonicus* were retrieved from PomBase and JaponicusDB (*S. pombe*: https://www.pombase.org/data/genome_sequence_and_features/gff3/Schizosaccharomyces_pombe_all_chromosomes.gff3.gz; *S. japonicus*: https://www.japonicusdb.org/data/genome_sequence_and_features/gff3/Schizosaccharomyces_japonicus_all_chromosomes.gff3.gz). The GFF3 files of *S. octosporus* and *S. cryophilus* were downloaded from Ensembl Fungi (*S. octosporus*: ftp://ftp.ensemblgenomes.org/pub/fungi/release-51/gff3/schizosaccharomyces_octosporus/Schizosaccharomyces_octosporus.GCA_000150505.2.51.gff3.gz; *S. cryophilus*: ftp://ftp.ensemblgenomes.org/pub/fungi/release-51/gff3/schizosaccharomyces_cryophilus/Schizosaccharomyces_cryophilus.GCA_000004155.2.51.gff3.gz). The GFF3 file of *S. osmophilus* was generated in this study as described above.

### Functional annotation of protein-coding genes

Functional annotation of protein-coding genes in the *S. osmophilus* nuclear genome was conducted with the aid of the orthogroup information. For an *S. osmophilus* gene, if it has an *S. pombe* ortholog and the ortholog is not annotated as a “hypothetical protein”, the gene name and the product description of the *S. pombe* ortholog were assigned to it. If it does not have an *S. pombe* ortholog, we examined whether it has an ortholog in another fission yeast species (in the order of *S. japonicus*, *S. octosporus*, and *S. cryophilus*), and if an ortholog is found and the ortholog is not annotated as a “hypothetical protein”, the product description of the ortholog is assigned to it. All remaining genes not functionally annotated were submitted to eggNOG-mapper (online version, http://eggnog-mapper.embl.de/) (Cantalapiedra *et al*. 2021) for ortholog searching and functional annotation. Genes without reliable hits in eggNOG-mapper were annotated as “hypothetical protein”. Based on the discrepancy reports generated during the submission of the genome to NCBI, we manually adjusted the annotations of some genes to conform with the requirement of NCBI.

### Annotation of non-coding RNA genes

For tRNA genes in the nuclear genome, tRNAScan-SE (v2.0.8, http://lowelab.ucsc.edu/tRNAscan-SE/) was used for prediction (Chan *et al*. 2021). For rRNA genes in the nuclear genome, rnammer (v1.2, https://services.healthtech.dtu.dk/service.php?RNAmmer-1.2) (Lagesen *et al*. 2007) and barrnap (v0.9, https://github.com/tseemann/barrnap) were used for prediction. snRNA and snoRNA genes in the nuclear genome were predicted by Infernal (v1.1.3, http://eddylab.org/infernal/) (Nawrocki and Eddy 2013) and BLASTN searches using snRNA and snoRNA genes of *S. pombe* as queries. Infernal outputs and BLASTN hits were merged using custom scripts. The following four functional non-coding RNA genes were also annotated using BLASTN and/or synteny analysis: *srp7* (7SL signal recognition particle component), *mrp1* (RNase MRP component), *rrk1* (RNase P component), and *ter1* (telomerase component).

### Species tree inference and estimation of the divergence time

To infer the phylogenetic relationship of fission yeast species, we used single-copy BUSCO genes of five fission yeast species and an outgroup species *Saitoella complicata*, which also belongs to the *Taphrinomycotina* subphylum of the *Ascomycota* phylum (Liu *et al*. 2009). The genome assembly of *S. complicata* was downloaded from NCBI (https://ftp.ncbi.nlm.nih.gov/genomes/all/GCF/001/661/265/GCF_001661265.1_Saico1/GCF_001661265.1_Saico1_genomic.fna.gz) and submitted to BUSCO (v3.0.2, https://busco.ezlab.org/) to assess the presence or absence of 1315 predefined single-copy orthologs of *Ascomycota* (ascomycota_odb9 gene set) (Simão *et al*. 2015). A total of 1060 “complete and single-copy” BUSCO genes present in all five fission yeast species and *S. complicata* were used for phylogenetic analysis. For each of these BUSCO genes, protein sequences generated by BUSCO analysis were aligned using MAFFT (v7.475) with the options “--thread 4 --auto --maxiterate 1000” (Katoh and Standley 2013). The resulting multiple sequence alignments were trimmed using trimAL (v1.4.rev15, http://trimal.cgenomics.org/) with the option “-gappyout” (Capella-Gutiérrez *et al*. 2009).

For each trimmed alignment, the best-fitting amino acid substitution model was inferred using the IQ-TREE build-in ModelFinder (v 2.0.3, https://github.com/Cibiv/IQ-TREE) with options “-m TESTONLY -nt 1” (Kalyaanamoorthy *et al*. 2017). All trimmed multiple sequence alignments were concatenated into a single alignment using catsequences (https://github.com/ChrisCreevey/catsequences/tree/73c11ef) (Creevey and Weeks 2021). A concatenation-based ML tree was inferred using IQ-TREE (v2.0.3, https://github.com/Cibiv/IQ-TREE) (Minh *et al*. 2020) with parameters -m LG + G4 -alrt 1000 -bb 1000. The “LG + G4” model was used because it is the best-fitting model for a majority of BUSCO genes (61.4%, 651 of 1060 BUSCO genes, Table S11).

The RelTime method in MEGA11 was employed to estimate the divergence time between *S. osmophilus* and *S. octosporus* using the branch lengths of the concatenation-based ML tree as input (Tamura *et al*. 2021). *S. complicata* was used as outgroup in the analysis. We used three calibration nodes based on a recently published time-calibrated phylogeny of *Ascomycota*: the *S. japonicus*–*S. pombe* split (207.2 million years ago), the *S. pombe*–*S. octosporus* split (108.2 million years ago), and the *S. octosporus*–*S. cryophilus* split (29.4 million years ago) (Shen *et al*. 2020). To assess the robustness of the analysis, we also carried out the analysis using only two calibration nodes (the *S. japonicus*–*S. pombe* split and the *S. pombe*–*S. octosporus* split) and only one calibration node (the *S. japonicus*–*S. pombe* split). Similar divergence times between *S. octosporus* and *S. osmophilus* were obtained (using three calibration nodes: 15.7 million years; using two calibration nodes: 15.5 million years; using one calibration node: 14.3 million years).

The species tree was visualized using Figtree (v1.4.4, http://tree.bio.ed.ac.uk/software/figtree/). For the timetree in Figure 1F, the outgroup species *S. complicata* was removed manually. A geologic time scale was manually added according to a document retrieved from the TimeTree resource (http://www.timetree.org/public/data/pdf/Gradstein2009Chap03.pdf) (Kumar *et al*. 2017).

### Synteny analysis of nuclear genomes

To visualize the syntenic relationship of the nuclear genomes of two species, single-copy orthologous gene pairs were extracted from the proteinortho output. Two genes of the same orthologous gene pair were linked using a colored line in Circos (v0.69, http://circos.ca/) (Krzywinski *et al*. 2009). The *S. octosporus* and *S. cryophilus* genomes used in this analysis are the PacBio sequencing-based genome assemblies downloaded from http://bifx-core.bio.ed.ac.uk/~ptong/genome_assembly/ (Tong *et al*. 2019). The *S. pombe* and *S. japonicus* genomes used in this analysis are reference genomes downloaded from PomBase and Ensembl Fungi respectively.

### Centromere-related analyses

The centromeric regions in the *S. osmophilus* genome were identified based on the synteny of centromere-flanking protein-coding genes (Table S13) (Ács-Szabó *et al*. 2018).

Centromeric repeats were identified by all-to-all comparison of centromeric sequences using BLASTN and YASS (Noé and Kucherov 2005) and manually annotated. The central core (*cnt*) region in each centromere was identified as the central region flanked by a pair of near-perfect inverted repeat sequences because this is a conserved characteristics of the *cnt* regions in *S. pombe*, *S. octosporus*, and *S. cryophilus* (Pidoux and Allshire 2004; Tong *et al*. 2019). The annotation of retrotransposons and LTRs was described in a section below. Plots depicting centromeric sequence features were generated using gggenomes (https://github.com/thackl/gggenomes) (Hackl *et al*. 2021) and manually adjusted in Adobe Illustrator.

To identify conserved sequence motifs enriched in the *cnt* regions, de novo motif discovery was performed using the MEME (Multiple Em for Motif Elicitation) tool in the MEME Suite (version 5.4.1) with the *cnt* sequences from *S. osmophilus*, *S. octosporus*, and *S. cryophilus* as input (Bailey *et al*. 2009). The “Any Number of Repetitions (anr)” option was selected. We ran MEME multiple times, with the “maximum width (maxw)” parameter set at a different value each time. Based on the recommendation given by the authors of the MEME Suite (Bailey *et al*. 2009), we chose to set the maxw parameter at all integers from 10 to 20, and at 25, 30, 35, 40, 45, and 50. After manual inspection of all MEME output, a 11-bp motif was selected as the conserved *cnt* motif because it can be found in all nine *cnt* sequences from the three species and has strong *p*-values (*p* < 1E-7) in multiple MEME runs. To determine the genomic distribution of this motif, we used the FIMO (Find Individual Motif Occurrences) tool of the MEME Suite to scan the genomes of *S. osmophilus*, *S. octosporus*, *S. cryophilus,* and *S. pombe*. The FIMO hits were filtered using the *p*-value cutoff of 3E-5 because all occurrences of the motif in the *cnt* sequences as reported by MEME can pass this cutoff.

### Telomere-related analyses

The analysis of telomeric repeat units present in the HiFi reads was performed using the software TweenMotif (https://download.cnet.com/TweenMotif/3000-2054_4-75325319.html) (Box *et al*. 2008). Incomplete repeat units at the end of the HiFi reads were filtered out.

The telomerase RNA gene (*ter1*) of *S. osmophilus* was identified by synteny and similarity (Ard *et al*. 2014; Kannan *et al*. 2015).

### Transposon-related analyses

Automated whole-genome de-novo transposable element (TE) annotation was performed using the Extensive de novo TE Annotator (EDTA) version 2.0.0 (Ou *et al*. 2019). Candidate transposons identified by EDTA were filtered by removing the ones overlapping with coding genes unrelated to transposons and the ones lacking any transposon-related protein domains that can be identified by TEsorter version 1.3 (Zhang *et al*. 2022). No intact DNA transposons, non-LTR retrotransposons, or Ty1/Copia superfamily LTR retrotransposons were found. The only intact transposons identified were four Ty3/Gypsy superfamily retrotransposons designated as Tosmo1-1, Tosmo1-2, Tosmo2-1, and Tosmo3-1 (Tosmo1-1 and Tosmo1-2 are identical in sequence) (Table S18). The LTR sequences and internal sequences of these intact retrotransposons were used as curated sequence input to perform another round of EDTA annotation. EDTA-annotated LTRs and internal sequences were filtered by removing the ones overlapping with coding genes unrelated to transposons and verified by TEsorter analysis and manual inspection of sequence alignments. For LTRs judged to be incomplete based on sequence alignment, we added flanking sequences, performed sequence alignment against, and manually evaluated where their borders can be extended. If an internal sequence and a neighboring LTR were of the same orientation and the interval between them was no longer than 5 bp, they were merged into a retrotransposon. Retrotransposon sequences generated by the merging operation were verified by manual inspection of sequence alignments. The start and end coordinates of LTRs and retrotransposons are listed in Table S18. In that table, LTRs are classified into four types (A, B, C, and D) based on a phylogenetic analysis (Figure S9A). For brevity, we used the midpoint coordinates of LTRs and retrotransposons to denote their locations in Figure S9 (the same location information is also provided in Table S18). The sequences of Tosmo1-1, Tosmo2-1, and Tosmo3-1 (including 50-bp upstream and 50-bp downstream flanking sequences) have been deposited at GenBank under accession numbers OP263985–OP263987.

The only copy of full-length Tcry1 retrotransposon in *S. cryophilus*, Tcry1-1, has an annotated length of 5055 bp (Rhind *et al*. 2011; Tong *et al*. 2019). Rhind et al. 2011 annotated two identical LTRs of 374 bp for Tcry1-1. Close inspection indicated that the 40-bp sequence immediately downstream of the annotated Tcry1-1 sequence should be part of the 3′ LTR, and the lengths of the 5′ LTR and the 3′ LTR should be 406 bp and 414 bp respectively. The length difference between the two LTRs is due to a 8-bp sequence (ATTTTCCC, at positions 393–400 of the LTR) being tandemly duplicated in the 3′ LTR. Apart from this duplication, the two LTRs are identical in sequence. The currently annotated version of Tcry1-1 does not have a TSD, whereas the revised version of Tcry1-1 has a perfect TSD of 5 bp (TTTAA). In the currently annotated version of Tcry1-1, the inverted repeats at the two ends of the LTR are 3-bp long (5’-TGT…ACA-3’), whereas in the revised version of Tcry1-1, the inverted repeats are 5-bp long (5’-TGTCA…TGACA-3’) and are identical to the inverted repeats in Tosmo1, Tosmo2, Tosmo3, and Tj1. The revised LTR annotation is also supported by the presence in the *S. cryophilus* genome a closely related solo LTR (NW_013185624.1 coordinates 2876278–2875872, minus strand), which is 407 bp long and 94% identical to the 406-bp 5′ LTR of the revised version of Tcry1-1 and possesses a perfect TSD of 5 bp (TGAAT). The revised annotation of Tcry1-1 (including 50-bp upstream and 50-bp downstream flanking sequences) (ACQJ02000037.1 coordinates 13333–18527, minus strand) has been deposited at the Third Party Annotation (TPA) section of GenBank under the accession number BK061829.

Retrotransposon sequences used for phylogenetic analysis were sequences of accession numbers given in the legend of Figure 4. The accession numbers of most of the non-fission-yeast retrotransposons were obtained from GyDB (https://gydb.org/) (Llorens *et al*. 2011). The exceptions are PpatensLTR2, Tcn1, and grasshopper. The accession number of PpatensLTR2 (GQ294565) was from (Novikova *et al*. 2010). The same publication provided an accession number for Tcn1 (XM_571377). XM_571377 is the mRNA of the predicted gene CNF03140 in the genome of *Cryptococcus neoformans* var. *neoformans* strain JEC21 (Loftus *et al*. 2005). The CNF03140 gene encompasses the coding sequences of Gag and Pol of Tcn1 but contains a wrongly annotated intron, presumably due to the inability of the gene prediction procedure to accommodate the programmed frameshift between the coding sequences of Gag and Pol. We manually annotated the Tcn1 sequence at the CNF03140 locus. The annotation of this copy of Tcn1 (including 50-bp upstream and 50-bp downstream flanking sequences) (AE017346.1 coordinates 926449–934423, minus strand) has been deposited at the Third Party Annotation (TPA) section of GenBank under the accession number BK061830.

The grasshopper retrotransposon was first discovered in strains of the phytopathogenic fungus *Magnaporthe grisea* that infect members of the plant genus *Eleusine* (Dobinson *et al*. 1993). Based on current taxonomy, these *Eleusine*-infecting strains should belong to the fungal species *Pyricularia oryzae* (Couch and Kohn 2002; Zhang *et al*. 2016). Dobinson et al. deposited a 5233-bp sequence of grasshopper at GenBank (M77661.1), which contains the 5′ LTR and the coding sequences of Gag and Pol but lacks the 3′ LTR. The length of grasshopper was reported to be approximately 8 kb (Dobinson *et al*. 1993). Thus, M77661.1 is missing more than 2 kb of the sequence of grasshopper. We performed BLASTN search using M77661.1 as query and identified 72 high-scoring hits (query length coverage = 100% and identity > 99.5%) in the genome of the *Eleusine*-infecting *Pyricularia oryzae* strain MZ5-1-6 (GenBank assembly accession GCA_004346965.1) (Gómez Luciano *et al*. 2019). Compared to M77661.1, these BLASTN hits all have an extra nucleotide after the 33rd nucleotide downstream of the stop codon of the Gag ORF. As a result, unlike the situation of M77661.1, where the Pol ORF is in the +1 frame relative to the Gag ORF, in these BLASTN hits, the Pol ORF is in the −1 frame relative to the Gag ORF. It is likely that the missing nucleotide in M77661.1 is due to a sequence error. Manual inspection of the six top BLASTN hits, which are identical to each other and differ from M77661.1 by 5 substitutions and 1 indel, and their flanking sequences showed that these six BLASTN hits are each part of a full-length and intact copy of grasshopper with identical LTRs at 5′ and 3′ ends. The six copies have nearly identical sequences (lengths ranging from 7614 to 7629 bp and all pairwise identities >= 99.8%). Five of the six copies have perfect 5-bp TSDs. We deposited the annotation of a representative copy (7615 bp long, TSD = TAAAT) with 50-bp upstream and 50-bp downstream flanking sequences (CP034205.1 coordinates 4282303–4290017, minus strand) at the Third Party Annotation (TPA) section of GenBank under the accession number BK061831.

Alignment of the amino acid sequences of RT and IN was performed using MAFFT v7.149b with the E-INS-i algorithm (Katoh and Standley 2013). Aligned sequences were examined manually using Jalview version 2.11.2.3 (Waterhouse *et al*. 2009). Maximum likelihood trees were calculated using IQ-TREE version 2.0.3 (Minh *et al*. 2020). The best-fitted substitution model was selected using ModelFinder (Kalyaanamoorthy *et al*. 2017) as implemented in IQ-TREE. Ten independent IQ-TREE runs were performed and the tree with the best log-likelihood value was chosen. 1000 ultrafast bootstraps (UFBoot) were performed using the ‘-bb’ command and 1000 Shimodaira–Hasegawa approximate likelihood ratio test (SH-aLRT) replicates were performed using the ‘-alrt’ command (Guindon *et al*. 2010; Minh *et al*. 2013). Rooting and visualization of trees were performed using FigTree version 1.4.3 (http://tree.bio.ed.ac.uk/software/figtree/).

Transcriptional start sites in the 5′ LTRs of Tosmo1-1, Tosmo2-1, and Tosmo3-1, and Tcry1-1 were predicted based on multiple sequence alignment that includes the LTRs of Tf1 and Tf2. The “pretzel” structures were identified by visual inspection of the RNA secondary structures predicted using the MXfold2 web server (http://www.dna.bio.keio.ac.jp/mxfold2/) (Sato *et al*. 2021). The alignment showing the conserved sequence context of the in-frame stop codons and the alignment showing the conservation of sequences upstream of PPTs were visualized using ggmsa (Zhou *et al*. 2022) and manually adjusted in Adobe Illustrator.

The sliding-window analysis was conducted using SimPlot++ v1.3 (Samson *et al*. 2022). The following parameters were used: Distance model = identity, Window length = 40, Step = 5, Strip gap = No Strip Gap, and Plot refresh rate = every window. For the comparison between Tosmo1, Tosmo2, and Tosmo3, an MAFFT-generated alignment of their sequences was given as input to SimPlot++ and three pairwise comparison plots were created. For the comparison between Tf1 and Tf2, an MAFFT-generated alignment of their sequences was given as input to SimPlot++ and a pairwise comparison plot was created. The plots generated by SimPlot++ were manually adjusted in Adobe Illustrator.

### *wtf* genes-related analyses

*wtf* genes in another *S. osmophilus* strain CBS 15792 have been identified and annotated in our recently published study (De Carvalho *et al*. 2022). We calculated the average nucleotide identity (ANI) between the genomes of CBS 15792 and the type strain CBS 15793^T^ using OrthoANIu (Yoon *et al*. 2017). As a comparison we calculated ANIs between genomes of six representative pure-lineage *S. pombe* strains (JB22, JB760, JB869, JB758, JB837, and JB864) (https://db.cngb.org/search/project/CNP0001878/) (Tusso *et al*. 2022). OrthoANIu analysis results are shown in Table S19. The synteny between the nuclear genome assemblies of CBS 15792 and CBS 15793^T^ was analyzed using nucmer in mummer-4.0.0 (Marçais *et al*. 2018). The smallest of the 11 nuclear contigs in the CBS 15792 genome assembly, tig00007777_pilon_x4, which has a length of 12408 bp and does not contain any *wtf* genes, was not aligned to the CBS 15793^T^ genome by nucmer. *wtf* genes in the CBS 15793^T^ genome that are syntenic to *wtf* genes in the CBS 15792 genome were identified as follows: for each *wtf* gene in the CBS 15792 genome, its flanking non-repetitive protein-coding genes were extracted, and their counterparts in the CBS 15793^T^ genome were identified by BLASTP analysis of the amino acid sequences of gene products. The thus defined genomic regions in the CBS 15793^T^ genome were inspected for the presence of *wtf* genes. Annotated protein-coding genes falling into these regions and sharing sequence similarities with *wtf* genes in CBS 15792 were classified as *wtf* genes. For regions lacking any annotated protein-coding genes resembling *wtf* genes, BLASTN analysis was performed to identify *wtf* pseudogenes. *wtf* pseudogenes not annotated by the MAKER pipeline were given systematic names but were not included in the protein-coding genes listed in Table S4. The coordinates of *wtf* genes (including *wtf* pseudogenes) in CBS 15793^T^ are listed in Table S20. Pairwise nucleotide identities were calculated using Sequence Demarcation Tool Version 1.2 (SDTv1.2) (Muhire *et al*. 2014) and the default Needleman-Wunsch algorithm (as implemented in MUSCLE) was used.

### Mating-type region-related analyses

Sequences of mating-type regions of *S. pombe*, *S. octosporus*, *S. cryophilus*, and *S. osmophilus* (provided as GenBank format files in Supplementary Files S1–S4) were obtained as follows: the *S. pombe* sequence was derived from the reference genome sequence by replacing the chromosome 2 region spanning coordinates 2129208-2137121 with the sequence of the “mating type region” contig (GenBank accession FP565355) and by replacing the *M* cassette sequence at the *mat1* locus with the *P* cassette sequence; the *S. octosporus* sequence was from the Tong et al. genome assembly (http://bifx-core.bio.ed.ac.uk/~ptong/genome_assembly/) (Tong *et al*. 2019); the *S. cryophilus* sequence was derived from the Tong et al. genome assembly by replacing the *M* cassette sequence at the *mat1* locus with the *P* cassette sequence and by changing a nucleotide in the Pc-coding sequence that causes a premature stop codon to the nucleotide in the reference *S. cryophilus* genome; the *S. osmophilus* sequence was from the genome assembly generated in this study. For analyses of the mating-type loci of *S. japonicus*, we used sequences of PCR-cloned *mat1* locus and donor region (GenBank accessions JQ735907 and JQ735908) (Yu *et al*. 2013). We note that the coding sequence of *S. japonicus* Pi in these sequences differs from the coding sequence of Pi in the reference *S. japonicus* genome. A 1-bp deletion in the reference *S. japonicus* genome causes a frameshift in the coding sequence of Pi.

The synteny plot was generated using clinker and clustermap.js (Gilchrist and Chooi 2021). Pairwise amino acid identities were calculated using Sequence Demarcation Tool Version 1.2 (SDTv1.2) (Muhire *et al*. 2014) and the default Needleman-Wunsch algorithm (as implemented in MUSCLE) was used. Amino acid sequence alignments of cassette-encoded proteins and nucleotide sequence alignment of the cassettes and the SAS region were generated using MAFFT (Katoh and Standley 2013) and visualized using Jalview (Waterhouse *et al*. 2009).

Annotations of H1, H2, H3, and inverted repeats (IRs) in the *S. pombe* sequence are according to GenBank accession FP565355. Annotations of H1, H2, and H3 in the *S. japonicus* sequences are according to GenBank accession JQ735908. H1, H2, H3, and IRs in *S. octosporus*, *S. cryophilus*, and *S. osmophilus* sequences were annotated by manually inspecting the self-to-self sequence comparison results generated by YASS (Noé and Kucherov 2005). The structures of the Pi–Mi complexes were predicted using AlphaFold-Multimer with default parameters (Evans *et al*. 2022). The predicted structures were visualized using the Mol* Viewer (https://molstar.org/) (Sehnal *et al*. 2021). Among the structures predicted, we chose the one with the highest confidence score. The interface area of each complex was calculated using the PDBePisa web server (https://www.ebi.ac.uk/pdbe/pisa/) (Krissinel and Henrick 2007).

Transposon-related sequences in the donor regions of *S. octosporus*, *S. cryophilus*, and *S. osmophilus* were identified by BLASTN analysis using the LTRs and the internal sequences of full-length transposons Tf1, Tf2, Tosmo1, Tosmo2, Tosmo3, Tcry1, and Tj1 as queries. Multiple sequence alignment shown in Figure 5C was generated by adding INT-related sequences found in the donor regions into an alignment of internal sequences of Tcry1-1, Tosmo1-1, Tosmo2-1, and Tosmo3-1 using the --addfragments option of MAFFT, and was visualized using seqvisr v0.2.6 (https://github.com/vragh/seqvisr) (Raghavan 2021).

### Cbp1 family-related analyses

TBLASTN and BLASTP searches using 21 fission yeast Cbp1 family proteins as queries were performed using the BLAST web server at NCBI (https://blast.ncbi.nlm.nih.gov/Blast.cgi). Terminal inverted repeats (TIRs), a common characteristic of DNA transposons, were searched by manually inspecting the self-to-self sequence comparison results generated by YASS (Noé and Kucherov 2005). Sequence alignment and maximum likelihood tree construction were performed as described above. Pairwise amino acid identities were calculated using Sequence Demarcation Tool Version 1.2 (SDTv1.2) (Muhire *et al*. 2014) and the default Needleman-Wunsch algorithm (as implemented in MUSCLE) was used. Local synteny plots were generated using clinker and clustermap.js (Gilchrist and Chooi 2021).

### Mitogenome-related analyses

Protein-coding genes in the mitogenome were annotated using the MFannot mitogenome annotator based on genetic code 4 (https://megasun.bch.umontreal.ca/apps/mfannot/) (Prince *et al*. 2022). Mitochondrial introns were classified using RNAweasel (https://megasun.bch.umontreal.ca/apps/rnaweasel/) (Prince *et al*. 2022). Mitochondrial tRNA genes were annotated based on the predictions of tRNAscan-SE using the sequence source option “Other mitochondrial” (http://trna.ucsc.edu/tRNAscan-SE/) (Lowe and Chan 2016) and the predictions of RNAweasel. Initiator formylmethionyl-tRNA gene and elongator methionyl-tRNA gene were distinguished based on similarity to tRNA genes in annotated mitogenomes of other fission yeast species. rRNA genes were annotated based on cross-species conservation revealed by BLASTN analysis. The *rnpB* (RNase P RNA) gene was annotated based on the result of RNAweasel analysis and manually adjusted based on homology to *S. octosporus rnpB*. Gene boundaries and exon-intron junctions were manually examined and adjusted when necessary. The sequence and annotation of the *S. osmophilus* mitogenome have been deposited at GenBank under the accession number OP310968.

The circular graphic map of the mitogenome was generated as follows: The coordinates of genes and DHEs were transformed into a tab-delimited text file using custom scripts. The GC content of sliding windows with window size = 35 bp and step size = 5 bp was calculated using bedtools nuc v2.30.0 (https://bedtools.readthedocs.io/en/latest/) (Quinlan and Hall 2010) and the calculation result was transformed into a tab-delimited text file using custom scripts. These text files were given as input to Circos v0.69 (Krzywinski *et al*. 2009). The backgrounds representing the GC content ranges of 0–70% and 70–100% were defined in the “plot.conf” file of Circos. The resulting Circos plot was manually modified in Adobe Illustrator. Phylogenetic analysis was performed in the same manner as described earlier for the nuclear genome. Codon usage was analyzed using Geneious Prime (Dotmatics).

Plots showing the synteny between the mitogenomes of fission yeast species were generated using the graphics module of the JCVI utility libraries (https://github.com/tanghaibao/jcvi/wiki/MCscan-%28Python-version%29) (Tang *et al*. 2015). As input for JCVI, we manually constructed a BED file containing the coordinates of the genes and a “blocks” file containing the matching relationships between genes in different species. The plots were merged and adjusted using Adobe Illustrator. Local synteny plots for the mitogenome region containing *trnI(cau)* and the nuclear genome region containing *til1* were generated using clinker and clustermap.js (Gilchrist and Chooi 2021) and manually adjusted using Adobe Illustrator. Pairwise amino acid identities were calculated using Sequence Demarcation Tool Version 1.2 (SDTv1.2) (Muhire *et al*. 2014) and the default Needleman-Wunsch algorithm (as implemented in MUSCLE) was used. The sliding-window analysis was conducted using SimPlot++ v1.3 (Samson *et al*. 2022). The following parameters were used: Distance model = identity, Window length = 40, Step = 5, Strip gap = No Strip Gap, and Plot refresh rate = every window.

DHEs were identified using a GC scanning approach based on the GC-rich characteristics of *S. octosporus* DHEs. GC-rich regions were found using bedtools nuc (window size = 35 bp, step size = 1 bp, GC content > 75%) and then inspected visually for the potential to form double-hairpin structures. The sequence alignment of DHEs and the accompanying arc diagram were generated using ggmsa (Zhou *et al*. 2022) and manually adjusted using Adobe Illustrator.

### Data Availability Statement

Sequencing data have been deposited at NCBI SRA (BioProject: PRJNA870673). Genome assemblies have been submitted to GenBank (accession number OP310968 for the mitogenome and submission number SUB1225547 for the nuclear genome).

## Results and Discussion

### Assembly of the genome of the *S. osmophilus* type strain CBS 15793^T^

To assemble the genome of the *S. osmophilus* type strain CBS 15793^T^, we generated 430.07 Mb (∼35× coverage assuming a 12 Mb genome) of PacBio HiFi read data using the PacBio Sequel II platform, and 4.86 Gb (∼405× coverage assuming a 12 Mb genome) of Illumina paired-end read data using the Illumina NovaSeq 6000 platform (Table S1). After contig assembly, polishing, and filtering (Figure S1A), we obtained a total of six contigs (Figure 1D and Table S2). One of the six contigs is the mitochondrial genome, which will be described in detail in a later section. Among the other contigs are three megabase-sized contigs (5.57 Mb, 3.50 Mb, and 2.35 Mb respectively). The total length of these three contigs (11.42 Mb) is similar to the total lengths of nuclear genome contigs in the reference genomes of the other four fission yeast species (12.57 Mb, 11.27 Mb, 11.52 Mb, and 11.13 Mb for *S. pombe*, *S. octosporus*, *S. cryophilus*, and *S. japonicus*, respectively) (Wood *et al*. 2002; Rhind *et al*. 2011). The six ends of these three contigs are either telomeric repeat sequences (described in detail in a later section) or sequences of 18S–5.8S–25S rRNA genes (rDNA) (Figure 1E), which are the types of sequences found at or next to chromosome ends in other fission yeast species. Within each contig, one centromere was identified based on synteny of centromere-flanking genes (Figure 1E). These observations suggest that like other fission yeast species, *S. osmophilus* has three chromosomes and these three contigs correspond to the three chromosomes. By convention, we numbered the three chromosomes from the longest to the shortest and designated the short arm of each chromosome as the left arm.

The rDNA repeat arrays are not fully assembled in the reference genomes of *S. pombe*, *S. octosporus*, *S. cryophilus*, or *S. japonicus*. Cutting-edge sequencing technologies available today still do not allow the full assembly of long arrays of rDNA repeats. In the recently published telomere-to-telomere human genome assembly, the three longest rDNA arrays are not fully assembled and have to be modeled instead (Nurk *et al*. 2022). In our *S. osmophilus* genome assembly, there are approximately two copies, five copies, and one copy of rDNA repeat unit at the left end of chromosome 2 contig, the left end of chromosome 3 contig, and the right end of chromosome 3 contig, respectively (Figure 1E), suggesting that like the situation in *S. octosporus* and *S. cryophilus* (Tong *et al*. 2019), *S. osmophilus* has three rDNA arrays. None of the rDNA arrays is fully assembled as no telomeric repeats are present at their outer ends. Based on Illumina read depth coverage, we estimated that there are a total of about 185 copies of rDNA repeat units in the *S. osmophilus* nuclear genome.

The remaining two contigs in our *S. osmophilus* assembly are less than 100 kb long. They both contain a few copies of rDNA repeat units at one end and telomeric repeats at the other end. They share a nearly identical (99.98% identity) 15.3-kb sequence between rDNA and telomeric repeats but differ in where the rDNA/non-rDNA junction falls within an rDNA repeat unit (Figure S1B). We named them rDNA-distal contig 1 and 2 (Figure 1D). The Illumina read depth coverage of the two types of rDNA/non-rDNA junctions showed an approximately 2:1 ratio (Figure S1C), suggesting that rDNA-distal contig 1 may correspond to sequences located outside of the distal ends of two rDNA arrays and rDNA-distal contig 2 may correspond to the sequence distal to the other rDNA array.

The quality of our *S. osmophilus* genome assembly was assessed in multiple ways. First, we found that 99.45% of PacBio HiFi reads and 99.62% of cleaned Illumina reads could be mapped onto the genome assembly, indicative of its completeness. Second, mapping results using both PacBio HiFi reads and Illumina reads showed even depth coverage across chromosomes, with the only exception being the rDNA repeats (Figure S2A), suggesting that there are no copy number errors of repeat regions outside of the rDNA arrays. Third, variant calling using HiFi reads only identified three SNPs and 32 indels and variant calling using Illumina reads only identified one SNP and 12 indels. The two variant sets overlap by one SNP and 6 indels and all of them are in rDNA repeats, likely due to sequence variations in the unassembled rDNA repeat units. Thus, the genome assembly is virtually error-free at the base level. Fourth, we analyzed the presence or absence of 1315 Benchmarking Universal Single Copy Orthologs (BUSCO) (ascomycota_odb9 gene set) (Simão *et al*. 2015) in our *S. osmophilus* genome assembly and found that 1222 (92.9%) of them are classified as “complete and single copy” and 29 (2.2%) of them are classified as “complete and duplicated” (Table S3). This level of BUSCO completeness is similar to those of published genome assemblies of the other four fission yeast species (Table S3 and Figure S2B). Together, these results demonstrate that we have obtained a highly contiguous, complete, and correct genome assembly of the type strain of *S. osmophilus*.

### Gene annotation and orthogroup analysis

To facilitate gene annotation, we obtained transcriptome data of vegetatively growing cells of the *S. osmophilus* type strain CBS 15793^T^ using both Illumina short read-based RNA-seq and long-read cDNA sequencing on the Oxford Nanopore Technologies (ONT) platform (Table S1). High-confidence transcript sequences were generated from these transcriptome data using the FLAIR (Full-Length Alternative Isoform analysis of RNA) workflow (Tang *et al*. 2020), and were provided to the annotation pipeline MAKER as EST evidence (Campbell *et al*. 2014). By incorporating additional evidence including ab initio prediction and annotated protein sequences of other four fission yeast species, protein-coding genes of *S. osmophilus* were predicted (Figure S3A). Upon further manual inspection and adjustment, a total of 5098 protein-coding genes were annotated in the three chromosomal contigs (Figure S3C and Table S4). We also annotated nuclear genes encoding functional non-coding RNAs including rRNAs, tRNAs, snRNAs, snoRNAs, and four other functional RNAs (*srp7*, *mrp1*, *rrk1*, and *ter1*) (Figures S3B–S3C and Tables S5–S9).

To establish evolutionary relationships between protein-coding genes of different fission yeast species, we inferred orthologous gene groups (orthogroups, OGs) encompassing nuclear-encoded protein-coding genes of five fission yeast species using proteinortho, a tool that takes into account both similarity and synteny (Lechner *et al*. 2011, 2014). 4873 (95.6%) of the 5098 *S. osmophilus* genes fall into 4864 orthogroups that contain genes of at least one other fission yeast species (Table S10 and Figure S4A). 4122 (80.9%) *S. osmophilus* genes fall into 4120 orthogroups that contain genes in all five fission yeast species (Table S10 and Figure S4A).

We assigned functional annotations to *S. osmophilus* protein-coding genes by giving them the gene product descriptions of the orthologs in other fission yeast species, especially *S. pombe* (Figure S4B). For genes without proteinortho-detected orthologs in other fission yeast species and for genes whose orthologs in other fission yeast species have the gene product description of “hypothetical protein”, we resorted to the tool eggNOG-mapper to obtain functional annotations (Cantalapiedra *et al*. 2021) (Figure S4B). In total, descriptive annotations other than “hypothetical protein” were obtained for 4868 (95.5%) *S. osmophilus* protein-coding genes (Table S4).

### Species phylogeny and evolutionary rate difference

We used the assembled nuclear genome of the *S. osmophilus* type strain to examine the phylogenetic position of *S. osmophilus*. First, we performed a maximum likelihood analysis using the amino acid sequences of 1060 “complete and single-copy” BUSCO genes present in all five fission yeast species as well as the outgroup species *Saitoella complicata*, which also belongs to the *Taphrinomycotina* subphylum of *Ascomycota* (Liu *et al*. 2009) (Figure S5A and Table S11). The sequences of these genes were generated by BUSCO analysis (Simão *et al*. 2015), which only uses genome sequences as input and is independent of genome annotations. The maximum likelihood tree shows that *S. osmophilus* shares a more recent common ancestor with *S. octosporus* than with other fission yeast species (Figure S5A).

We used the BUSCO-gene-inferred phylogeny to estimate the divergence time between *S. osmophilus* and *S. octosporus* (Figure 1F). The divergence times of the *S. japonicus*–*S. pombe* split (207.2 million years ago), the *S. pombe*–*S. octosporus* split (108.2 million years ago), and the *S. octosporus*–*S. cryophilus* split (29.4 million years ago) reported in a recently published timetree of *Ascomycota* were used as time calibration nodes (Shen *et al*. 2020). This analysis showed that *S. osmophilus* and *S. octosporus* diverged about 15.7 millions years ago. This divergence time is similar to the divergence time between humans and orangutans (genus *Pongo*) (Kumar *et al*. 2017).

To corroborate the BUSCO-based results, we used the 4085 orthogroups containing single-copy orthologs in all five fission yeast species (1:1:1:1:1 orthologs) to calculate pairwise amino acid identities between orthologous proteins from two different fission yeast species (Table S12). Mean pairwise identities were calculated for all species pairs (Figure S5B). Consistent with the BUSCO-based species tree, the orthologs in *S. osmophilus* and *S. octosporus* have the highest mean pairwise identity of 90.52%.

Interestingly, the maximum likelihood tree shows that the evolutionary rate of *S. octosporus* is 51% higher than that of *S. osmophilus* since their divergence from each other (0.0622 substitutions per site for *S. octosporus* vs. 0.0412 substitutions per site for *S. osmophilus*) (Figure S5A). Consistent with this observation, the mean pairwise identity between *S. octosporus* and *S. cryophilus* orthologs (85.51%) is lower than that between *S. osmophilus* and *S. cryophilus* orthologs (87.06%) (Figure S5B). We plotted histograms of pairwise identities of the 4085 orthogroups and observed unimodal distributions of identity values, with the distribution for the *S. octosporus*–*S. cryophilus* comparison slightly shifted to smaller values compared to that for the *S. osmophilus*–*S. cryophilus* comparison (Figure S5C). To directly analyze the differences in identity values for the two comparisons, we calculated the differences in pairwise identities for orthologs belonging to the same orthogroup and plotted histograms of the differences (Figure S5D). The difference values show unimodal distributions not centered at zero, confirming overall differences in identity values. For 74% of the 4085 orthogroups, the pairwise identity between *S. osmophilus* and *S. cryophilus* orthologs is larger than that between *S. octosporus* and *S. cryophilus* orthologs (Table S12).

It is known that evolutionary rates of species are inversely correlated with their generation times (Wu and Li 1985; Smith and Donoghue 2008; Thomas *et al*. 2010). It is possible that *S. octosporus* has a shorter generation time or spends less time in quiescence than *S. osmophilus* in native habitats and therefore has evolved faster. The second possibility is consistent with the known natural habitats of these two species (Brysch-Herzberg *et al*. 2022): the primary habitat of *S. octosporus* is honeybee honey, which can be available all year round in a honeycomb, whereas the main habitat of *S. osmophilus* is bee bread of solitary bees, which only lasts for a maximum of a few weeks before being fully consumed by the larva in a nest chamber.

During evolution, both gene sequence divergences and changes in gene order accumulate over time. To visualize how the gene order in the nuclear genome of *S. osmophilus* differs from those in other fission yeast species, we generated pairwise synteny plots based on single-copy orthologs (Figures 1G–1J). Consistent with the species phylogeny inferred from sequence divergences, *S. osmophilus* shares the highest level of synteny with *S. octosporus*. There are only a few large-scale chromosomal rearrangements between these two species. One of these rearrangements is a reciprocal translocation with breakpoints falling into two centromeres (described in more details below).

### Centromeres of *S. osmophilus* and their comparison with centromeres of other fission yeasts

Centromeres in the *S. osmophilus* genome were identified based on the conserved synteny of centromere-flanking protein-coding genes (Ács-Szabó *et al*. 2018; Tong *et al*. 2019) (Table S13). For *cen1*, *cen2*, and *cen3* of *S. osmophilus*, the lengths of the regions between the two closest flanking protein-coding genes are approximately 56 kb, 57 kb, and 72 kb, respectively (Figure 2A), together accounting for 1.6% of the total length of the chromosomal contigs. These regions are the most prominent low-GC-content regions in the nuclear genome and are also the regions with the highest densities of tRNA genes (Figure 1E).

**Figure 2.**
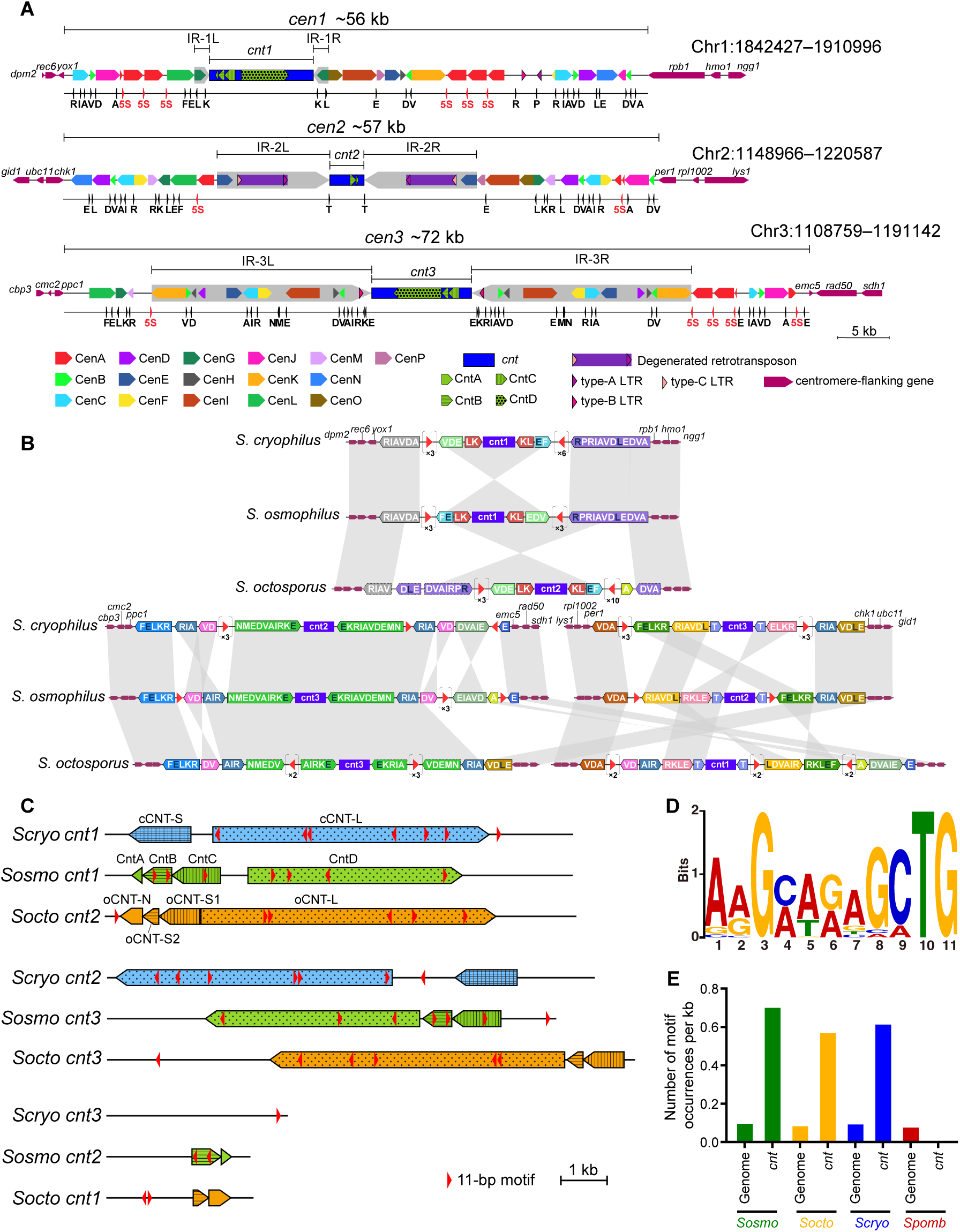
Centromeres of *S. osmophilus*. (A) Diagrams depicting the organization of *S. osmophilus* centromeres. For each centromere, features are shown on two lines. Shown on the top line are centromere-flanking genes (thin maroon arrows), centromeric repeats (colored arrows), the central core (*cnt*, dark blue rectangle), inverted repeats flanking the central core (IRs, thick grey arrows), retrotransposon (purple rectangle), and long-terminal repeats (LTRs, colored arrowheads with black border). Shown on the bottom line are 5S rRNA genes (red arrowheads) and tRNA genes (black arrowheads with single-letter codes of cognate amino acids). These diagrams and all features within them are drawn to scale. (B) Diagrams depicting the synteny of tRNA genes and 5S rRNA genes in the centromeres of *S. cryophilus*, *S. osmophilus*, and *S. octosporus*. tRNA genes are denoted using single-letter codes of cognate amino acids. Isoacceptor tRNAs that accept the same amino acid but have different anticodons are distinguished by the color of the letter (white: tRNA-ArgACG, tRNA-GluTTC, and tRNA-LeuCAA; black: tRNA-ArgTCG, tRNA-GluCTC, and tRNA-LeuAAG). Syntenic tRNA gene clusters are denoted by colored arrows. The orientations of tRNA gene clusters are defined as follows: among the letters representing the tRNA genes in a tRNA gene cluster, the letter occurring earliest in the alphabet is selected and the orientation of its corresponding tRNA gene is taken as the orientation of the tRNA gene cluster. The orientations of partial tRNA gene clusters (dashed line border) are set to be the same as the orientations of their corresponding full clusters. 5S rRNA genes are shown as red arrowheads. A cluster of tandemly arranged 5S rRNA genes is shown as an arrowhead in brackets, with the number of 5S rRNA genes in the cluster shown below. Features are not drawn to scale. (C) Diagrams depicting the arrangements of *cnt* repeats in the *cnt* regions of *S. cryophilus*, *S. osmophilus*, and *S. octosporus*. Orthologous *cnt* regions are grouped together as in (B). The orientations of the *cnt* regions of *S. osmophilus* are as in (A). The orientations of the *cnt* regions of the other two species are set in a way so as to best show the similarity of the arrangements of *cnt* repeats. The locations and orientations of the cCNT-S and cCNT-L repeats of *S. cryophilus* and the oCNT-S and oCNT-L repeats of *S. octosporus* are as defined previously (Tong *et al*. 2019). oCNT-S is split into oCNT-S1 and oCNT-S2. oCNT-N is a type of repeat identified in this study. Orientations of oCNT-N and the four types of *cnt* repeats of *S. osmophilus* are set in a way so as to best show the similarity of the arrangements of *cnt* repeats. The *cnt* regions and repeats within them are shown to scale in these diagrams. The distribution of the MEME-identified 11-bp motif (red arrowhead, not drawn to scale) is also shown in the diagrams. (D) Sequence logo of the MEME-identified 11-bp motif. (E) The density of the 11-bp motif (number of motif occurrences per kb) within the *cnt* regions and across the entire nuclear genome of each of four fission yeast species.

To investigate the organization of *S. osmophilus* centromeres, we performed all-to-all comparison of the sequences of the three centromeric regions and found that like the situation in other fission yeast species (Rhind *et al*. 2011; Tong *et al*. 2019), these regions are extensively occupied by repetitive sequences (Figure 2A and Table S14). Similar to the centromeres of *S. octosporus*, *S. cryophilus*, and *S. pombe* (Tong *et al*. 2019), in each *S. osmophilus* centromere, a central core (*cnt*, blue rectangles in Figure 2A) is flanked by a pair of inverted repeats (IRs, thick grey arrows in Figure 2A). The lengths of *cnt1*, *cnt2*, and *cnt3* are 10.1 kb, 3.1 kb, and 9.7 kb, respectively, and the lengths of IRs flanking them are 1.5 kb, 11.1 kb, and 21.4 kb, respectively. Sequence similarity exists not only within the same centromere, but also between different centromeres. We identified 20 types of repeats present in more than one centromere (colored arrows in Figure 2A). Four types locate exclusively within *cnt* regions and are designated CntA– CntD (Figures 2A and 2C, Table S14). The other 16 types locate outside of *cnt* regions and are designated CenA–CenP (Figure 2A and Table S14). We will refer to these repeats outside of *cnt* regions as peri-core repeats.

There are 13 5S rRNA genes in the centromeres of *S. osmophilus* (Figure 2A), accounting for 13% of the 5S rRNA genes in the nuclear genome (Table S5). Nine of them form three clusters, with each cluster containing three tandemly arranged 5S rRNA genes. The other four 5S rRNA genes are situated alone. *S. octosporus* and *S. cryophilus* have higher number of 5S rRNA genes in their centromeres (25 and 20 respectively), and they each have multiple types of 5S rRNA gene-associated repeats (termed Five-S-Associated Repeats, FSARs), some of which harbor protein-coding sequences (Tong *et al*. 2019). In *S. osmophilus*, there is only one type of centromeric repeats associated with 5S rRNA genes: the CenA repeats. CenA repeats are usually flanked by two tandemly arranged 5S rRNA genes (Figure 2A) and likely result from the homogenizing effect of 5S rRNA gene-mediated recombination. CenA repeats do not share sequence similarity with FSARs in *S. octosporus* and *S. cryophilus* and do not contain protein-coding sequences.

There are 96 tRNA genes in the centromeres of *S. osmophilus* (Figure 2A), accounting for 33% of the tRNA genes in the nuclear genome (Table S6). This high level of tRNA gene enrichment in the centromeres is similar to the situations in *S. octosporus*, *S. cryophilus*, and *S. pombe* (Tong *et al*. 2019). Two types of peri-core repeats are flanked on both sides by tRNA genes (CenB and CenD) and many other types of peri-core repeats are flanked on one side by tRNA genes (Figure 2A), suggesting that tRNA genes may contribute to the formation of these repeats through tRNA gene-mediated recombination. There are two degenerated full-length retrotransposons and four solo long terminal repeats (LTRs) in the centromeres of *S. osmophilus* (Figure 2A) (see a later section for detailed description of *S. osmophilus* retrotransposons). Overall, centromeres of *S. osmophilus* are not rich in retrotransposon-related sequences. This is similar to the situation in *S. octosporus* and *S. cryophilus* (Tong *et al*. 2019).

Chromatin immunoprecipitation followed by next-generation sequencing (ChIP-Seq) analyses in *S. pombe*, *S. octosporus*, and *S. cryophilus* have shown that *cnt* regions are assembled in CENP-A^Cnp1^ chromatin and outer centromeric repeats are assembled in H3K9me2 heterochromatin, with the CENP-A/heterochromatin transitions demarcated by the most *cnt*-proximal tRNA gene clusters or LTRs (Tong *et al*. 2019). We speculate that similar situations may occur in *S. osmophilus*.

It has been shown that *S. octosporus* and *S. cryophilus*, but not *S. pombe*, share synteny of tRNA genes and 5S rRNA genes in centromeres (Tong *et al*. 2019). We compared centromeric tRNA genes and 5S rRNA genes of *S. osmophilus*, *S. octosporus*, and *S. cryophilus* (Figure 2B). The number, types, order, and orientation of centromeric tRNA genes are strongly conserved among these three species. Synteny of tRNA genes often extends across a long cluster of tRNA genes. There can be as many as 12 tRNA genes in a syntenic cluster. The small number of breakdowns of synteny appear to be mainly caused by intra-centromeric rearrangements. For example, tRNA gene order difference between *S. osmophilus cen1* and its orthologous centromere in *S. cryophilus* can be explained by a single inversion event that may have occurred through recombination between 5S rRNA genes. Compared to centromeric tRNA genes, centromeric 5S rRNA genes show weaker synteny and appear to have undergone relatively frequent gain and loss events, suggesting that they are under a more relaxed constraint during evolution. No obvious cross-species sequence similarity can be detected in intervening sequences between syntenic tRNA genes and 5S rRNA genes.

*S. octosporus* and *S. pombe*, but not *S. cryophilus*, share perfect synteny of protein-coding genes flanking the three centromeres (Tong *et al*. 2019). Thus, it is thought that the situation in *S. octosporus* and *S. pombe* represents the ancestral state. However, our analysis shows that *S. osmophilus* and *S. cryophilus* share identical synteny of centromere-flanking genes (Figure 2B), suggesting that the situation in these two species may represent the state in the common ancestor of *S. osmophilus*, *S. octosporus*, and *S. cryophilus*. Regardless of which situation is the ancestral state of these three species, the break of synteny can be attributed to an inter-centromeric translocation, with one breakpoint at or near a tRNA gene cluster (the RIA cluster) situated between *cnt3* and *emc5* in *S. osmophilus* and the other breakpoint at or near another RIA cluster situated between *cnt2* and *chk1* in *S. osmophilus* (Figure 2B).

No cross-species sequence similarity is known to exist between the *cnt* regions of different fission yeast species (Tong *et al*. 2019). However, there appear to be conserved arrangements of repeats in *cnt* regions of *S. octosporus* and *S. cryophilus*, as both species contain *cnt* repeats of different lengths (termed oCNT-L and oCNT-S for the long and short *S. octosporus* repeats respectively and cCNT-L and cCNT-S for the long and short *S. cryophilus* repeats respectively) and they are arranged in similar ways in orthologous centromeres (Tong *et al*. 2019). Our BLASTN analysis of the sequences of the *cnt* regions of *S. osmophilus*, *S. octosporus*, and *S. cryophilus* revealed no cross-species sequence similarity. Interestingly, we identified previously unrecognized sequence similarity among the three *cnt* regions of *S. octosporus*. No *cnt* repeats were previously known to exist in *cnt1* of *S. octosporus*. We found that a repeat of approximately 480 bp, which we named oCNT-N (N for new), is present in *cnt1* and *cnt2* of *S. octosporus* (Figures 2C and S6). In addition, a part of oCNT-S is also present in *cnt1*. Therefore, we split oCNT-S into two parts: oCNT-S1 and oCNT-S2. oCNT-S1 is present in *cnt2* and *cnt3*, and oCNT-S2 in present in all three *cnt* regions (Figures 2C and S6).

There are four types of *cnt* repeats in *S. osmophilus*: CntA–D (Figures 2C and S7 and Table S14). They are arranged in a remarkably similar way to the four types of *cnt* repeats of *S. octosporus* (Figure 2C). In Figure 2C, orthologous *cnt* regions in *S. cryophilus*, *S. osmophilus*, and *S. octosporus* are grouped together to highlight cross-species similarities in *cnt* repeat distribution and positioning. CntA is present in *cnt1* and *cnt2* of *S. osmophilus* and mirrors the distribution of oCNT-N in *S. octosporus*. CntB is present in all three *cnt* regions of *S. osmophilus* and resembles oCNT-S2 in distribution. CntC is located in *cnt1* and *cnt3* of *S. osmophilus* and matches the locations of oCNT-S1. CntD is the longest *cnt* repeat in *S. osmophilus*. It is present in *cnt1* and *cnt3*, whose orthologous *cnt* regions in *S. octosporus* and *S. cryophilus* harbor the long repeats oCNT-L and cCNT-L, respectively. Within each *cnt* region of *S. osmophilus*, the relative positioning of different *cnt* repeats always perfectly matches that of their counterparts in *S. octosporus*. The conserved *cnt* repeat arrangements suggest that inter-*cnt* sequence similarities have been selected for during evolution and can be maintained long after cross-species *cnt* sequence similarity is no longer detectable.

Even though no cross-species sequence similarities in *cnt* regions were found by BLASTN analysis, we wondered whether it is possible that short conserved sequence motifs, which cannot be detected by BLASTN, may be enriched in *cnt* regions. To test this possibility, we applied the motif discovery tool MEME (Multiple Em for Motif Elicitation) (Bailey *et al*. 2009). A 11-bp motif was found in the *cnt* regions of *S. osmophilus*, *S. octosporus*, and *S. cryophilus* (Figure 2D). For each of these three species, the density of the occurrences of the 11-bp motif in the *cnt* regions is about seven times of the average density across the nuclear genome (Figure 2E and Table S15). This motif is present in all nine *cnt* regions of these three species (Figure 2C), but is absent in the *cnt* regions of *S. pombe* (Figure 2E and Table S15). It is possible that this motif may be of functional relevance.

### Telomeres of *S. osmophilus* and their comparison with telomeres of other fission yeasts

Telomeres are composed of tandem arrays of short repeats synthesized by the reverse transcription enzyme telomerase, which contains a RNA subunit that serves as the template for telomeric repeat synthesis. We investigated telomeres of *S. osmophilus* and compared them to those of other fission yeast species.

In the *S. osmophilus* genome assembly, telomeric repeats were found at the three rDNA-free ends of the chromosomal contigs (Tel1L, Tel1R, and Tel2R) (Figures 3A and 3B). For the other three chromosome ends that contain rDNA repeats and are not fully assembled, telomeres are expected to locate at the far ends of rDNA-distal sequences (Tel2L, Tel3L, and Tel3R) (Figure 3A) and should correspond to the telomeric repeats in the two rDNA-distal contigs (Figure 3B).

**Figure 3.**
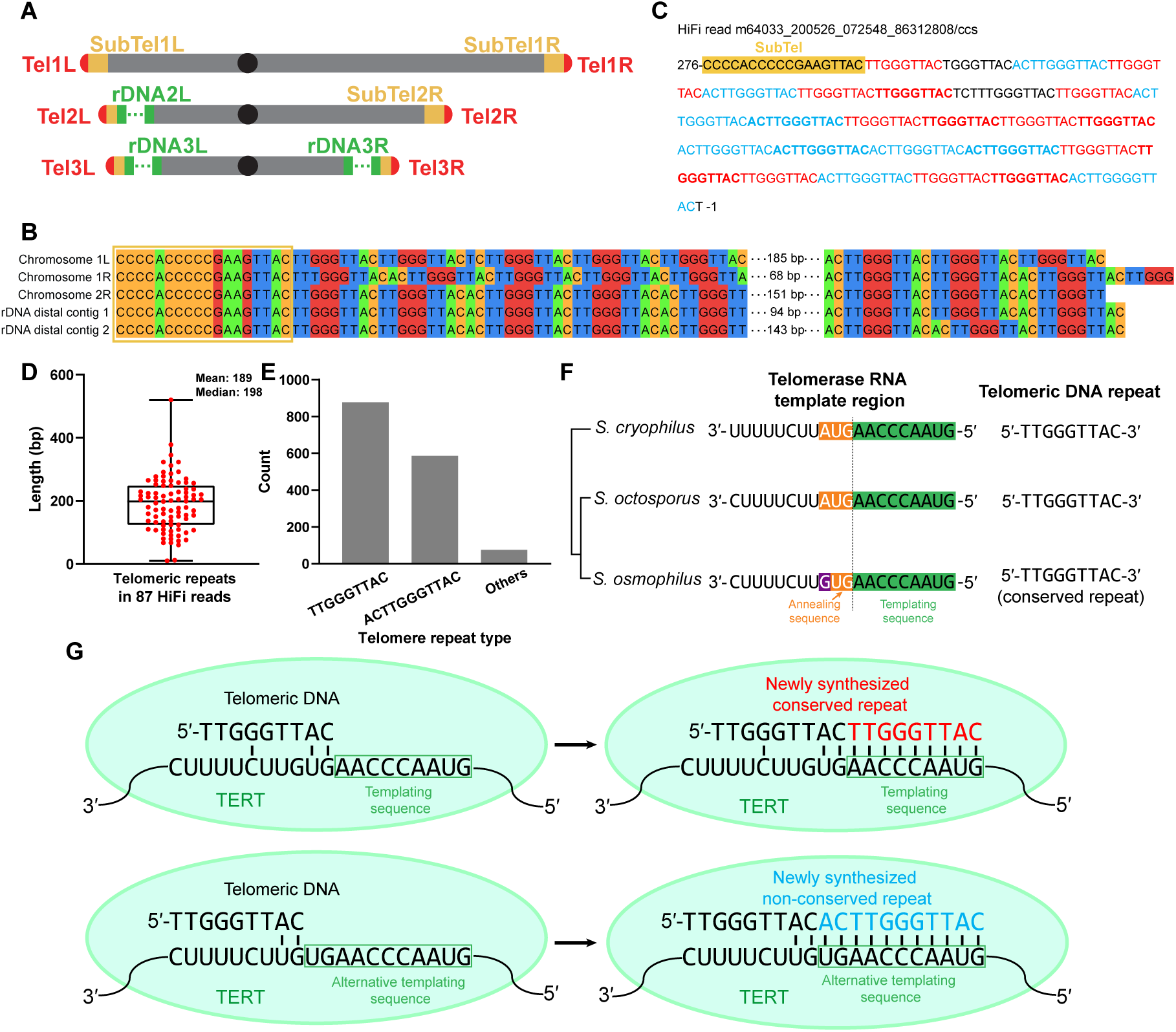
Telomeres of *S. osmophilus*. (A) Diagrams of the chromosomes in *S. osmophilus*. (B) Telomeric sequences present in the genome assembly of *S. osmophilus*. Sequences juxtaposed to telomeric repeats are boxed. (C) Telomeric sequence in a representative HiFi read containing telomeric repeats. The 9-bp conserved repeats are shown in red and the 11-bp non-conserved repeats are shown in blue. Other types of repeat units are shown in black. Neighboring repeats of the same type are distinguished by alternately using the regular font and the bold font. (D) Lengths of telomeric repeats in the 87 HiFi reads containing telomeric repeats. (E) The numbers of the conserved repeat, the non-conserved repeat, and other types of telomeric repeat units in the 87 HiFi reads containing telomeric repeats. (F) The template regions of the telomerase RNAs of three fission yeast species and the corresponding telomeric DNA repeats. The nucleotide that distinguishes *S. osmophilus* from the other two species is highlighted in purple. A schematic species tree is shown at left. (G) Models explaining why *S. osmophilus* has two types of telomeric repeat units. Watson-Crick base pairs are denoted by vertical lines. Telomerase reverse transcriptase (TERT) is the protein component of telomerase.

The telomeres in *S. octosporus* and *S. cryophilus* are composed of near-perfect arrays of the same type of 9-bp telomeric repeat unit: TTGGGTTAC (Rhind *et al*. 2011; Kannan *et al*. 2015) (Figures S8A and S8B). Manual inspection of the telomeric sequences in the *S. osmophilus* genome assembly revealed that there are two main types of telomeric repeat units, 9-bp TTGGGTTAC and 11-bp ACTTGGGTTAC (Figure 3B). To exclude the possibility of assembly artifacts, we examined the HiFi reads and confirmed that these two types of repeat units indeed co-exist in the same telomeric sequence (Figure 3C and Table S16). The 9-bp repeat unit is identical to the telomeric repeat unit in *S. octosporus* and *S. cryophilus*. We hereafter refer to the 9-bp repeat unit as the conserved repeat and the 11-bp repeat unit as the non-conserved repeat. We identified a total of 87 HiFi reads containing telomeric repeats (Table S16). The lengths of telomeric repeats in these HiFi reads show a normal distribution centered around approximately 200 bp (Figure 3D). The conserved repeat and the non-conserved repeat account for 57% and 38% of the repeat units in the HiFi reads, respectively, and other types of repeat units together only account for 5% of the repeat units (Figure 3E and Table S17).

To understand why *S. osmophilus* differs from *S. octosporus* and *S. cryophilus* in the telomeric repeat units, we compared the telomerase RNA genes (*ter1*) of these three species and paid special attention to the template regions of telomerase RNAs (Figures 3F, S8C, and S8D). The template region in a telomerase RNA usually consists of two parts: a “templating sequence” corresponding to a full telomeric repeat unit and serving as the template for reverse transcription, and on its 3′ side an “annealing sequence” that is identical to the 5′ part of the “templating sequence” and base pairs with the 3′ end of a telomeric DNA that has just finished one round of reverse transcription (Peska *et al*. 2020, 2021; Fajkus *et al*. 2021). The telomerase RNA genes of *S. osmophilus*, *S. octosporus* and *S. cryophilus* are well conserved (Figures 3F and S8D). *S. octosporus* and *S. cryophilus* have the same nine-nucleotide templating sequence (3′- AACCCAAUG-5′). This sequence is strictly conserved in *S. osmophilus* (Figure 3F). However, there is a notable difference in the annealing sequence. *S. octosporus* and *S. cryophilus* have the same three-nucleotide annealing sequence 3′-AUG-5′, which is identical to the 5′ trinucleotide of the templating sequence. In *S. osmophilus*, this sequence is 3′-GUG-5′. Consequently, the templating sequence of *S. osmophilus* telomerase RNA is flanked on its 3′ side by two tandem copies of the dinucleotide sequence 3′-UG-5′ (Figures 3F). We propose that either of the two 3′- UG-5′ sequences can base pair with the 3′-most dinucleotide of a telomeric DNA (Figure 3G). If the templating-sequence-proximal 3′-UG-5′ forms the base pairs, a 9-bp conserved telomeric DNA repeat will be synthesized in the ensuing round of reverse transcription; on the other hand, if the distal 3′-UG-5′ forms the base pairs, the proximal 3′-UG-5′ becomes part of the templating sequence and a 11-bp non-conserved repeat will be synthesized in the next round of reverse transcription.

### Retrotransposons in *S. osmophilus* and their evolutionary relationships with other retrotransposons

We surveyed the transposon content of the *S. osmophilus* genome using the Extensive de-novo TE Annotator (EDTA) pipeline (Ou *et al*. 2019). Similar to the situations in other fission yeast species (Bowen *et al*. 2003; Rhind *et al*. 2011; Tusso *et al*. 2022), the only type of intact transposons found in the *S. osmophilus* genome is Ty3/Gypsy superfamily long terminal repeat (LTR) retrotransposons (see Materials and Methods). We identified a total of four full-length and intact retrotransposons (Figure 1E and Table S18). They each contain two identical LTRs at their 5′ and 3′ ends, indicative of recent transposition. Two of these retrotransposons are identical in sequence, and we named them Tosmo1-1 and Tosmo1-2. The other two were named Tosmo2-1 and Tosmo3-1. The lengths of Tosmo1, Tosmo2, and Tosmo3 retrotransposons are 5098 bp, 5083 bp, and 5068 bp, respectively. The lengths of their LTRs are 395 bp, 392 bp, and 395 bp, respectively. Tosmo1-1, Tosmo1-2, and Tosmo2-1 but not Tosmo3-1 are flanked by perfect 5-bp target site duplications (TSDs) (Table S18). Using the sequences of these four retrotransposons to guide a new round of EDTA-based analysis, we were able to identify 15 degenerated retrotransposons and 63 solo LTRs in the *S. osmophilus* genome (Table S18). Phylogenetic analysis classified the LTRs in the *S. osmophilus* genome into four types (Figure S9A). Tosmo1 and Tosmo3 have type-A LTRs and Tosmo2 has type-B LTRs. Retrotransposons and solo LTRs in the *S. osmophilus* genome are not strongly enriched in any particular genomic regions (Figures 1E and S9B).

We performed comparative analysis of fission yeast retrotransposons, including the three types of *S. osmophilus* retrotransposons identified here and *S. pombe* Tf1 and Tf2, *S. cryophilus* Tcry1, and *S. japonicus* Tj1 (Figure 4A). Tf1 and Tf2 are extensively characterized retrotransposons (Levin *et al*. 1990; Weaver *et al*. 1993; Esnault and Levin 2015; Maxwell 2020). A survey of retrotransposons in a diverse set of *S. pombe* isolates found that full-length elements are mostly Tf1 or Tf2 and the rest are hybrid elements resulting from recombination between Tf1 and Tf2 (Tusso *et al*. 2022). Tcry1 is the only type of full-length retrotransposon known to exist in *S. cryophilus*. There is a single copy of Tcry1 in the genome of the only available strain of *S. cryophilus* (Rhind *et al*. 2011; Tong *et al*. 2019). We noticed that the downstream boundary of that copy of Tcry1 has been mis-annotated and its length should be 5095 bp, 40 bp longer than previously reported (see Materials and Methods). *S. japonicus* has a large variety of retrotransposons (Rhind *et al*. 2011; Chapman *et al*. 2022). However, the retrotransposons in the *S. japonicus* reference genome tend to have sequence errors (Rhind et al. 2011). Thus, we included in our analysis only one type of *S. japonicus* retrotransposon, Tj1, because a reliable sequence of a full-length Tj1 retrotransposon is available (Guo *et al*. 2015), even though there is no full-length Tj1 in the *S. japonicus* reference genome (Rhind et al. 2011). No full-length retrotransposons have been found in *S. octosporus* (Rhind *et al*. 2011; Tong *et al*. 2019).

**Figure 4.**
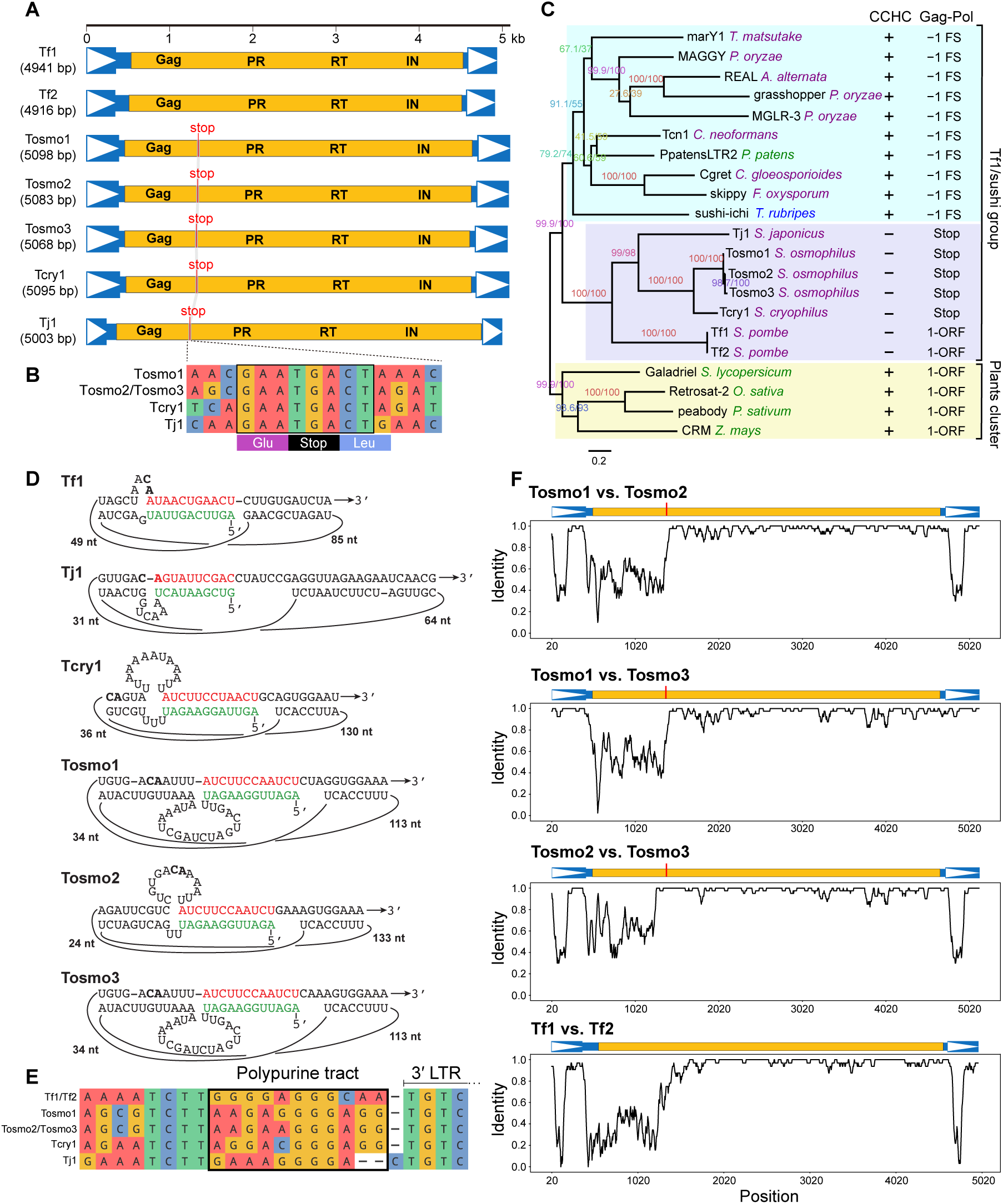
Identification of *S. osmophilus* retrotransposons and comparative analysis of fission yeast retrotransposons. (A) Diagrams showing the structures of 7 fission yeast retrotransposons. Blue-colored regions and yellow-colored regions correspond to non-coding and coding sequences, respectively, and are drawn to scale. Boxed arrowheads denote LTRs. Gag, PR, RT, and IN indicate sequences coding for Gag, protease, reverse transcriptase, and integrase, respectively. The vertical red line with “stop” indicates the in-frame stop codon between the coding sequences of Gag and Pol proteins. (B) Alignment of the nucleotide sequences of and surrounding the in-frame stop codons in Tosmo1, Tosmo2, Tosmo3, Tcry1, and Tj1. The eight strictly conserved nucleotides are boxed. (C) Maximum likelihood tree constructed using the amino acid sequences of the RT-IN region, which corresponds to amino acids 374–1256 of the Tf1 ORF. The tree was rooted using “Plants cluster” retrotransposons as outgroup. Branch labels are the SH-aLRT support value (%) and the UFBoot support value (%) calculated by IQ-TREE. The names of the species are colored as follows: blue for animal species, green for plant species, and purple for fungal species. Three major branches in the tree are highlighted by shades. The column with the header “CCHC” indicates whether a CCHC motif is present (+) or absent (−) in the Gag ORF of a retrotransposon. The column with the header “Gag-Pol” indicates which strategy is used to express the Gag-Pol fusion protein: −1 frameshift (−1 FS), in-frame stop codon (Stop), or a single ORF (1-ORF). We note that grasshopper was thought to use +1 frameshift (Gao *et al*. 2003), but our analysis showed that it uses −1 frameshift (see Materials and Methods). The accession numbers are as follows: *Tricholoma matsutake* marY1 (AB028236), *Pyricularia oryzae* MAGGY (L35053), *Alternaria alternata* REAL (AB025309), *Pyricularia oryzae* grasshopper (BK061831), *Pyricularia oryzae* MGLR-3 (AF314096), *Cryptococcus neoformans* Tcn1 (BK061830), *Physcomitrella patens* PpatensLTR2 (GQ294565), *Colletotrichum gloeosporioides* Cgret (AF264028), *Fusarium oxysporum* skippy (L34658), *Takifugu rubripes* sushi-ichi (AF030881), *S. japonicus* Tj1 (KT447435), *S. osmophilus* Tosmo1 (OP263985), *S. osmophilus* Tosmo2 (OP263986), *S. osmophilus* Tosmo3 (OP263987), *S. cryophilus* Tcry1 (BK061829), *S. pombe* Tf1 (M38526), *S. pombe* Tf2 (L10324), *Solanum lycopersicum* Galadriel (AF119040, nucleotides 12865–19060, minus strand), *Oryza sativa* Retrosat-2 (AF111709, nucleotides 25889–38686), *Pisum sativum* peabody (AF083074), *Zea mays* CRM (AY129008). (D) The “pretzel” structures that can be formed by the 5′ UTR regions of the retrotransposon RNAs. Self-primers situated at the 5′ ends of RNAs are highlighted in green. PBSs situated downstream of the 5′ LTRs are highlighted in red. The last two nucleotides of the 5′ LTR, which are always CA, are highlighted in bold. (E) Alignment of the nucleotide sequences of and surrounding the polypurine tracts (PPTs) in Tf1, Tf2, Tosmo1, Tosmo2, Tosmo3, Tcry1, and Tj1. The nucleotides of the PPTs are boxed. (F) Sliding-window analysis showing that the mosaic nature of the diversity between Tosmo1, Tosmo2, and Tosmo3 is similar to the pattern of diversity between Tf1 and Tf2.

Tf1, Tf2, Tosmo1, Tosmo2, Tosmo3, Tcry1, and Tj1 each encode four proteins: the structural protein Gag, the protease (PR), the reverse transcriptase (RT), and the integrase (IN) (Figure 4A). In Tf1 and Tf2, all proteins are encoded within a single open reading frame (ORF). In contrast, in Tosmo1, Tosmo2, Tosmo3, Tcry1, and Tj1, an in-frame stop codon separates the coding sequence for Gag and the coding sequence for the Pol polyprotein that encompasses PR, RT, and IN (Figure 4A). Such an in-frame stop codon is a known strategy employed by a number of retroviruses and retrotransposons to express a Gag-Pol fusion protein through programmed stop codon readthrough and to ensure that the level of the Gag-Pol fusion protein is lower than that of the Gag protein (Yoshinaka *et al*. 1985; Matthews *et al*. 1997; Leng *et al*. 1998; Gao *et al*. 2003; Forbes *et al*. 2007; Guo *et al*. 2015). The in-frame stop codon in Tj1 has been shown to be essential for its transposition activity (Guo *et al*. 2015).

We compared the sequences of and surrounding the in-frame stop codon in Tosmo1, Tosmo2, Tosmo3, Tcry1, and Tj1 (Figure 4B). Interestingly, a 8-bp sequence, which includes the stop codon (TGA), the preceding codon (GAA for glutamate), and the first two nucleotides of the following codon (CTA or CTG for leucine), is strictly conserved, suggesting not only a common ancestry, but also selective constraints preventing divergence. TGA is known to be the most readthrough-permissive stop codon (Firoozan *et al*. 1991; Jungreis *et al*. 2011; Cridge *et al*. 2018). The dinucleotide AA immediately upstream of a stop codon promotes readthrough in *Saccharomyces cerevisiae* (Tork *et al*. 2004). Several types of sequences immediately downstream of the TGA stop codon are known to favor readthrough in various eukaryotic organisms, including the nucleotide C (Li and Rice 1993; Bonetti *et al*. 1995; Jungreis *et al*. 2011; Cridge *et al*. 2018), the dinucleotide CT (Stiebler *et al*. 2014), and the tetranucleotide CTAG (Loughran *et al*. 2014). Thus, Tosmo1, Tosmo2, Tosmo3, Tcry1, and Tj1 appear to exploit readthrough mechanisms conserved in eukaryotic organisms to regulate the expression of the Gag-Pol fusion protein.

Ty3/Gypsy retrotransposons have been phylogenetically classified into two main branches: chromovirus and non-chromovirus (Marín and Lloréns 2000; Llorens *et al*. 2009, 2011; Neumann *et al*. 2019). Chromovirus retrotransposons usually contain a chromodomain at the C-terminus of the integrase, whereas non-chromovirus retrotransposons lack the chromodomain. Tf1, Tf2, Tosmo1, Tosmo2, Tosmo3, Tcry1, and Tj1 all contain a chromodomain (Figure S10A) and are thus chromovirus retrotransposons. Tf1 is the founding member of a sublineage of chromovirus retrotransposons, termed the Tf1/sushi group (Butler *et al*. 2001; Goodwin and Poulter 2001; Neumann *et al*. 2019). A common feature of Tf1/sushi retrotransposons is that they initiate reverse transcription using a self-priming mechanism first discovered in studies of Tf1 (Levin 1995, 1996; Lin and Levin 1997b; Butler *et al*. 2001; Goodwin and Poulter 2001). The Tf1/sushi group is equivalent to the “Fungi/Vertebrates cluster” of chromovirus retrotransposons in the classification system of the Gypsy Database of Mobile Genetic Elements (GyDB, https://gydb.org/) (Llorens *et al*. 2009, 2011). The term “Fungi/Vertebrates cluster” is a misnomer as Tf1/sushi retrotransposons are present not only in fungi and vertebrates, but also in a number of non-seed plant species (Novikova *et al*. 2010; Neumann *et al*. 2019). Previously published phylogenetic analyses have shown that Tf1/sushi retrotransposons form a monophyletic group and the closest relatives of Tf1/sushi retrotransposons are plant chromovirus retrotransposons that use host tRNAs to prime reverse transcription (Marín and Lloréns 2000; Goodwin and Poulter 2001; Levin 2007; Llorens *et al*. 2009; Novikova *et al*. 2010; Neumann *et al*. 2019). These plant chromovirus retrotransposons are referred to as the “Plants cluster” of chromovirus retrotransposons in GyDB (Llorens *et al*. 2009, 2011).

We performed a phylogenetic analysis of Tf1, Tf2, Tosmo1, Tosmo2, Tosmo3, Tcry1, and Tj1 together with other representative chromovirus retrotransposons using amino acid sequences encompassing the two most conserved proteins encoded by retrotransposons, RT and IN (excluding the less conserved chromodomain at the C-terminus of IN) (Figure 4C). This analysis showed that Tf1, Tf2, Tosmo1, Tosmo2, Tosmo3, Tcry1, and Tj1 form a highly supported monophyletic clade, which is sister to all other Tf1/sushi retrotransposons, suggesting that these fission yeast retrotransposons descent from an ancestral retrotransposon existing in the common ancestor of fission yeasts. Supporting their shared origin is the fact that the Gag proteins of these seven fission yeast retrotransposons do not contain a Cys-X_2_-Cys-X_4_-His-X_4_-Cys (CCHC) motif, which is widely present in the Gag proteins of retrotransposons and retroviruses (Covey 1986; Llorens *et al*. 2008), including the Gag proteins of other chromovirus retrotransposons (Figure 4C and Figure S11). As the common ancestor of fission yeasts existed more than 200 million years ago (Rhind *et al*. 2011; Shen *et al*. 2020), these fission yeast Tf1/sushi retrotransposons may have co-evolved with fission yeasts for over 200 million years.

Intriguingly, the RT-IN phylogeny is incongruent with the species tree of fission yeasts, as retrotransposons in *S. osmophilus* and *S. cryophilus* share a more recent common ancestor with *S. japonicus* Tj1 than with *S. pombe* Tf1 and Tf2. On the other hand, this phylogeny is consistent with the fact that Tf1 and Tf2 have a single ORF whereas Tosmo1, Tosmo2, Tosmo3, Tcry1, and Tj1 have an in-frame stop codon at the Gag-Pol junction. Tf1/sushi retrotransposons of non-fission-yeast species predominantly express the Gag-Pol fusion protein through programmed −1 frameshifting at the Gag-Pol junction, whereas the outgroup “Plants cluster” retrotransposons usually have a single ORF (Figure 4C) (Gao *et al*. 2003). We speculate that the ancestor of the seven fission yeast retrotransposons may be a single-ORF retrotransposon. The in-frame-stop-codon retrotransposons would have arisen from this ancestor before the divergence of fission yeast species and would later have been lost in the lineage leading to *S. pombe*. The single-ORF retrotransposons would have been lost in the lineage leading to *S. osmophilus* and *S. cryophilus*. The situation in *S. japonicus* awaits further characterization of its retrotransposon repertoire.

The self-priming of Tf1 requires a three-duplex RNA structure formed by the 5’ untranslated region (UTR) of the Tf1 mRNA (Lin and Levin 1997a) (Figure 4D). The central duplex is formed between the self-primer at the 5’ end of the mRNA (shown in green in Figure 4D) and the primer-binding site (PBS) located downstream of the 5’ LTR (shown in red in Figure 4D). The other two duplexes are each formed between a PBS-adjacent sequence and a 5’ LTR sequence located downstream of the self-primer. This type of RNA structure, dubbed the “pretzel” structure (Butler *et al*. 2001), can also be formed by the mRNAs of Tf2 and several other Tf1/sushi retrotransposons (Lin and Levin 1997a; b; Butler *et al*. 2001; Goodwin and Poulter 2001). We inspected the sequences of Tosmo1, Tosmo2, Tosmo3, Tcry1, and Tj1 and found that their mRNAs all have the potential to form the “pretzel” structure (Figure 4D), suggesting that they share a similar self-priming mechanism with Tf1.

During reverse transcription of retrotransposons and retroviral mRNAs, the synthesis of plus-strand cDNA is initiated from an RNA primer that corresponds to a purine-rich sequence located upstream and adjacent to the 3’ UTR (Wilhelm and Wilhelm 2001; Rausch and Le Grice 2004; Rausch *et al*. 2017). This sequence is termed the polypurine tract (PPT) (Majors and Varmus 1981). We inspected PPT sequences and PPT-flanking sequences of Tf1, Tf2, Tosmo1, Tosmo2, Tosmo3, Tcry1, and Tj1 (Figure 4E). Interestingly, even though their PPT sequences do not show strong conservation, the four nucleotides immediately upstream of the PPT are exclusively TCTT in these fission yeast retrotransposons. This feature is not conserved in other Tf1/sushi retrotransposons (Figure S10B). Thus, the PPT-adjacent TCTT sequence is a characteristic feature of these fission yeast retrotransposons. As the plus-strand primer of Tf1 is known to begin at the third nucleotide of the TCTT sequence (Atwood-Moore *et al*. 2005; Esnault and Levin 2015), the TTCT sequence may help define the 5’ end of the plus-strand primer. In addition, a T-rich sequence immediately upstream of the PPT, termed T-box or U-box, is important for plus-strand priming of several retroviruses and *Saccharomyces cerevisiae* retrotransposon Ty1 (Ilyinskii and Desrosiers 1998; Noad *et al*. 1998; Robson and Telesnitsky 1999; Wilhelm *et al*. 1999b). It is possible that the TCTT sequence is similarly important for plus-strand priming of these fission yeast retrotransposons.

To more comprehensively understand the relationship between Tosmo1, Tosmo2, and Tosmo3, we performed a sliding-window analysis of nucleotide identity using SimPlot++ (Figure 4F) (Samson *et al*. 2022). Consistent with the RT-IN tree, Tsomo1, Tosmo2, and Tosmo3 are very similar to each other in the Pol coding sequence (overall nucleotide identity > 96%). However, strikingly high sequence diversity exists outside of the Pol coding sequence. High sequence diversity is located in two separate regions: first, Tosmo2 strongly differs from Tosom1 and Tosmo3 in an approximately 140-bp segment in the U3 region of the LTR, with the downstream boundary of the segment only 9 bp away from the U3-R junction (Figure S10C); and second, Tosmo1, Tosmo2, and Tosmo3 differ substantially from each other in an approximately 780-bp region beginning from the 5’ UTR sequence downstream of the PBS and extending to the C-terminal region of the Gag coding sequence. The second diversity region extends about 140 bp further downstream to the beginning of the Pol coding sequence for the Tosmo1-Tosmo2 comparison and the Tosmo1-Tosmo3 comparison. At the amino acid level, the identities of Gag proteins are 34%, 32%, and 55% for Tosmo1-Tosmo2, Tosmo1-Tosmo3, and Tosmo2-Tosmo3 comparisons, respectively (Figure S11). Such mosaic patterns of diversity are remarkably similar to the situation of *S. pombe* Tf1 and Tf2, which share 98% nucleotide identity in the Pol coding sequence but are highly different from each other in a segment of the U3 region and in the Gag coding sequence and the adjacent 5’ UTR sequence (Figures 4F, S10D, and S11) (Weaver *et al*. 1993; Kelly and Levin 2005). We hypothesize that this similarity may not be coincidental but rather due to non-random formation of diversity regions and/or selective advantages conferred by certain diversity patterns.

As has been proposed before for Tf1 and Tf2 (Kelly and Levin 2005), the mosaic patterns of diversity between Tsomo1, Tosmo2, and Tosmo3 likely result from inter-element recombination between divergent retrotransposons. Inter-element recombination may happen either through recombination between chromosomally integrated retrotransposons, or through extrachromosomal reverse-transcription-related recombination (Kelly and Levin 2005; Drost and Sanchez 2019). The recombination breakpoint near the U3-R junction may be a reflection of reverse-transcription-related recombination happening at the minus-strand transfer step of reverse transcription, when the minus-strand cDNA that has been synthesized to the 5′-end of the retrotransposon RNA, i.e. the beginning of the R region in the 5′ LTR, is transferred to base pair with the R region in the 3′ portion of either the same RNA molecule or another RNA molecule co-packaged in the same virus-like particle (Wilhelm *et al*. 1999a; Drost and Sanchez 2019).

The U3 region of the LTR serves as the promoter for the transcription of the retrotransposon mRNA and contains cis-regulatory sequences such as the TATA box and transcription factor-binding sites (Bilanchone *et al*. 1993; Benachenhou *et al*. 2013). In *S. pombe*, a nucleosome positioned at the U3 region of the Tf2 retrotransposon by chromatin remodelers Fft2 and Fft3 prevents the transcription of full-length Tf2 transcript in unstressed cells (Persson *et al*. 2016). The diversification of the U3 sequence in the retrotransposons in *S. pombe* and *S. osmophilus* may widen transcriptional regulatory mechanisms and thus confer adaptability. The Gag proteins of retroviruses and retrotransposons are known to be the target of host defense mechanisms and self-restriction mechanisms (Sanz-Ramos and Stoye 2013; Saha *et al*. 2015; Cottee *et al*. 2021). The diversification of the Gag sequence in the retrotransposons in *S. pombe* and *S. osmophilus* may be beneficial to the evolutionary survival of these retrotransposons.

### Comparisons of the *wtf* genes in CBS 15792 and CBS 15793^T^

*wtf* (short for <u>w</u>ith <u>Tf</u> LTRs) genes, which represent the largest gene family in *S. pombe*, are selfish spore killer genes (Hu *et al*. 2017; Nuckolls *et al*. 2017). Our recently published study showed that *wtf* genes also exist in *S. octosporus*, *S. osmophilus*, and *S. cryophilus* (De Carvalho *et al*. 2022). That study identified 42 *wtf* genes (including 11 pseudogenes) in a draft genome assembly of the *S. osmophilus* strain CBS 15792. CBS 15792 and CBS 15793^T^ share an average nucleotide identity (ANI) of 99.73% (Table S19). As a comparison, ANIs between REF-clade *S. pombe* strains estimated to have diverged within the last few thousand years are around 99.86%, and ANIs between NONREF-clade *S. pombe* strains are around 99.38% (Table S19) (Tao *et al*. 2019). Consistent with the high nucleotide identity between CBS 15792 and CBS 15793^T^, 41 of the 42 *wtf* genes in CBS 15792 have syntenic counterparts in CBS 15793^T^ (Table S20 and Figure S12A). The *wtf31* gene in CBS 15792 does not have a syntenic counterpart in CBS 15793^T^. This presence–absence polymorphism is likely owing to recombination mediated by tandemly arranged 5S rRNA genes flanking the *wtf31* gene in CBS 15792 (Figure S12B).

Among the 41 pairs of syntenic *wtf* genes in CBS 15792 and CBS 15793^T^, two pairs show inter-strain differences in whether a gene is an active gene or a pseudogene (Table S20). *wtf7* in CBS 15792 is a pseudogene with a premature stop codon in exon 4, whereas its syntenic counterpart, *wtf41* in CBS 15793^T^, does not have the premature stop codon and is an active gene (Figure S12C). Conversely, *wtf33* in CBS 15792 is an active gene, but its syntenic counterpart, *wtf26* in CBS 15793^T^, is a pseudogene owing to a frameshifting 1-bp deletion in exon 2 (Figure S12D).

The within-pair nucleotide identities of 22 (54%) syntenic pairs are 100% and the within-pair nucleotide identities of 15 (37%) pairs are lower than 100% but higher than 98% (Table S20). Manual inspection of the other four pairs whose within-pair nucleotide identities fall below 98% showed that ectopic recombination is the likely cause of their higher levels of inter-strain divergence. Figure S12E shows one of these four pairs, *wtf13* in CBS 15793^T^ and its syntenic counterpart *wtf22* in CBS 15792. They share an overall nucleotide identity of 96.60%. However, within an approximately 250-bp region, the nucleotide identity between these two genes is only 85.36%. In this region, *wtf13* in CBS 15793^T^ is highly similar to *wtf15* in CBS 15793^T^ (97.98% identity), and *wtf22* in CBS 15792 is highly similar to *wtf32* in CBS 15793^T^ (98.33% identity) (Figure S12E). Thus, either *wtf13* in CBS 15793^T^ or *wtf22* in CBS 15792 may have undergone ectopic recombination in this region. These results suggest that ectopic recombination drives fast divergence of *wtf* genes in *S. osmophilus*.

### Cross-species comparisons of the mating-type region

Haploid fission yeast cells exist in one of two mating types, P (plus) or M (minus). Whether the mating type is P or M is determined by which of two cassettes, *P* or *M*, is present at the mating-type locus *mat1* (Klar *et al*. 2014). Homothallic fission yeast strains can switch mating type by replacing the cassette at *mat1* with a different cassette residing at a donor locus. There are two heterochromatin-silenced donor loci, *mat2-P* and *mat3-M*, which are adjacent to each other and are a few genes away from *mat1* in *S. pombe*, *S. octosporus*, and *S. cryophilus* (Rhind *et al*. 2011). The cassettes in the three *mat* loci are all flanked by homology boxes (H1 box on one side and H2 box on the other side), which serve as recombination substrates during mating type switching.

We examined the synteny of the mating-type region in different fission yeast species (Figure 5A). We did not include *S. japonicus* in this analysis because the mating-type region in the reference *S. japonicus* genome is misassembled (Yu *et al*. 2013). For ease of comparison, we chose to use sequences in which the cassette at *mat1* is the *P* cassette (GenBank format files of sequences shown in Figure 5A are provided as Supplementary Files S1–S4). Sequences are oriented such that *mat1* is on the left and the donor region is on the right. Genes flanking the *mat1* locus show strong synteny across *S. pombe*, *S. octosporus*, *S. cryophilus*, and *S. osmophilus*. Notably, *nvj2* and *sui1*, which are the third and fourth genes on the left side of *mat1* in all four species, are among the genes that show conserved proximity to the mating-type locus across the *Ascomycota* phylum (Riley *et al*. 2016; Hanson and Wolfe 2017). With the exception of *SPBC23G7.10c*, all genes locating in the 17-kb interval between *mat1* and *mat2-P* (the *L*-region) in *S. pombe* are syntenically conserved in *S. octosporus*, *S. cryophilus*, and *S. osmophilus*. In *S. pombe*, *mat2-P* and *mat3-M* have the same orientation (H2 on the left and H1 on the right) and are separated by a 11-kb interval (the *K*-region). In contrast, in *S. octosporus*, *S. cryophilus*, and *S. osmophilus*, *mat2-P* and *mat3-M* are in opposite orientations, with their H1 boxes facing each other, and are only about 600 bp apart (Figure 5A). This is likely the ancestral configuration as *mat2-P* and *mat3-M* of *S. japonicus* are also arranged in this way (Yu *et al*. 2013). In all four species, the donor region is separated from the closest neighboring genes by inverted repeats (IRs), which have been shown to be the boundaries separating the donor region heterochromatin from the neighboring euchromatin in *S. pombe*, *S. octosporus*, and *S. cryophilus* (Noma K *et al*. 2001; Tong *et al*. 2019). On the right side of the donor region, gene synteny is maintained between *S. octosporus*, *S. cryophilus*, and *S. osmophilus* but is broken between these three species and *S. pombe*.

**Figure 5.**
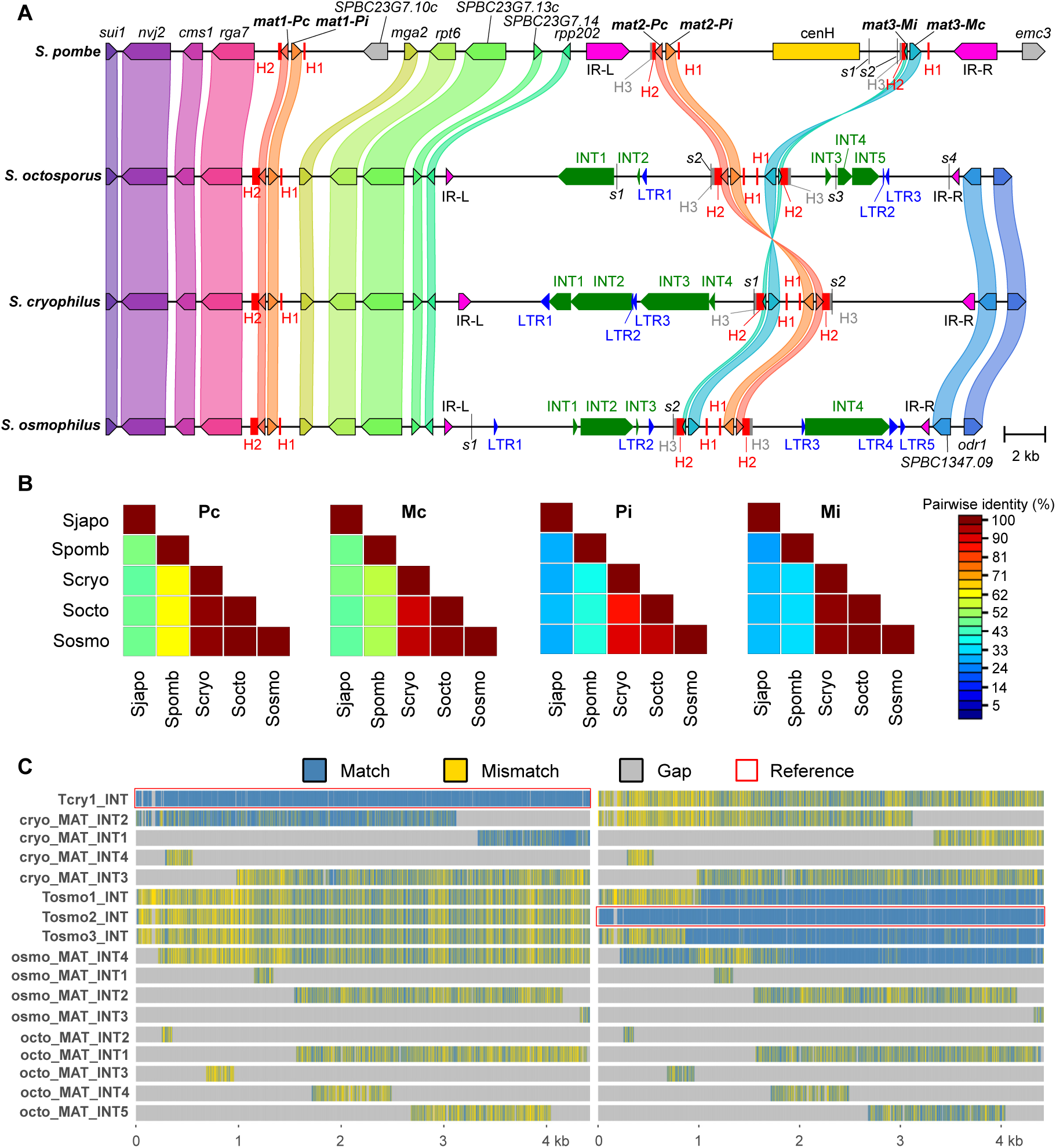
Cross-species comparisons of the mating-type region. (A) Synteny of the mating-type regions in *S. pombe*, *S. octosporus*, *S. cryophilus*, and *S. osmophilus*. For the two genes on the right side of the donor loci in *S. octosporus*, *S. cryophilus*, and *S. osmophilus*, the gene names shown are the names of the orthologous genes in *S. pombe*. Sequences sharing similarities with the LTRs and the internal sequences (INTs) of full-length retrotransposons are denoted by blue arrowheads and green arrows, respectively. The 8-bp Atf1-binding sites named *s1*, *s2*, etc. are denoted by black vertical lines. (B) Color matrices showing the pair-wise amino acid identities of mating-type cassette-encoded proteins of different fission yeast species. Color matrices were generated using Sequence Demarcation Tool Version 1.2 (SDTv1.2) (Muhire *et al*. 2014). (C) Multiple sequence alignment of the internal sequences (INTs) of Tcry1, Tosmo1, Tosmo2, Tosmo3, and INT-related sequences in the donor regions of *S. octosporus*, *S. cryophilus*, and *S. osmophilus*. Multiple sequence alignment was visualized using seqvisr v0.2.6 (https://github.com/vragh/seqvisr) (Raghavan 2021). The reference sequences used for visualization are the internal sequences of Tcry1 (left) and Tosmo2 (right).

In *S. pombe*, the *P* cassette encodes two proteins Pc and Pi and the *M* cassette encodes two proteins Mc and Mi (Kelly *et al*. 1988). Genes encoding Pc, Pi, and Mc but not Mi are annotated in the reference genomes of *S. octosporus*, *S. cryophilus*, and *S. japonicus* (Rhind 2011). An Mi-encoding gene has been annotated in the PCR-cloned mating-type donor region of a non-reference strain of *S. japonicus* (GenBank: JQ735908.1) (Yu *et al*. 2013). Our inspection showed that Mi-encoding genes exist in *S. osmophilus*, *S. octosporus* and *S. cryophilus* (Figure 5A). Pairwise comparisons and multiple sequence alignments of the amino acid sequences of Pc, Mc, Pi, and Mi showed that all four proteins are highly conserved among *S. osmophilus*, *S. octosporus*, and *S. cryophilus* (Figures 5B and S13A–D). Pc and Mc of these three species are moderately similar to their counterparts in *S. pombe* and *S. japonicus*, whereas Pi and Mi of these three species show rather low overall similarities to their homologs in *S. pombe* and *S. japonicus* (Figures 5B and S13A–D).

In *S. pombe*, Pi and Mi interact with each other in the zygote (Vještica *et al*. 2018). We predicted a heterodimeric structure of the *S. pombe* Pi–Mi complex using AlphaFold-Multimer (PDB format file provided as Supplementary File S5) (Evans *et al*. 2022). Pi contains a homeodomain in its C-terminal region (Kelly *et al*. 1988). In the predicted structure of the Pi–Mi heterodimer, the N-terminal region of Pi (amino acids 1–82), which consists of three α-helices, forms an intermolecular four-helix bundle with the 42-amino-acid Mi, which folds into a single α-helix (Figure S13E). Analysis of the predicted structure using the SOCKET2 web application showed that the hydrophobic core of the four-helix bundle forms knobs-in-holes packing typical of coiled coils (Kumar and Woolfson 2021). Pi and Mi of *S. osmophilus* are predicted by AlphaFold-Multimer to form a heterodimer with a structure resembling that of the *S. pombe* Pi–Mi complex (Figure S13F and S13G; PDB format file provided as Supplementary file S6). The Pi–Mi interface areas are 1335 Å^2^ and 1297 Å^2^ for the *S. pombe* complex and the *S. osmophilus* complex, respectively. Residues at the Pi–Mi interfaces are mostly hydrophobic, but also include a few charged residues that form interhelical salt bridges. These interface residues are substantially conserved in *S. osmophilus*, *S. octosporus*, *S. cryophilus*, and *S. pombe*, but not in *S. japonicus* (Figures S13C and S13D). AlphaFold-Multimer did not predict a confident heterodimeric structure for Pi and Mi of *S. japonicus*. Thus, Pi and Mi of *S. japonicus* may not interact or may interact in a different manner.

We also examined the conservation of cis-acting sequences at the three mating-type loci using BLASTN analysis and multiple sequence alignment. The H1 boxes in all five fission yeast species share similar sequences (Figures S14A and Table S21) (Rhind *et al*. 2011; Yu *et al*. 2013). The H2 boxes in *S. osmophilus*, *S. octosporus*, *S. cryophilus*, and *S. pombe* are highly similar in the most cassette-proximal 75 nucleotides, which partially overlap with the 3’ coding sequences of Pc and Mi (Figure S14B and Table S22). As a result of this overlap, Pc and Mi of *S. osmophilus*, *S. octosporus*, *S. cryophilus* share the same 12 amino acids in their C termini (Figures S13A and S13D). The H2 box in *S. japonicus* shares no obvious similarity with the H2 boxes in other species. For the H3 boxes, which locate next to the H2 boxes in the donor region, cross-species similarity is only detectable between *S. osmophilus* and *S. octosporus* (Table S23). No obvious cross-species similarity was detected for the inverted repeats flanking the donor regions (Table S24). Two cis-acting sequences called switch-activating sites (SAS1 and SAS2) are present in a 110-bp *L*-region segment immediately adjacent to H1 in *S. pombe* (Arcangioli and Klar 1991). Sequence alignment suggests that this *L*-region segment is conserved across five fission yeast species (Figure S14A) (Rhind *et al*. 2011; Yu *et al*. 2013). SAS1 in *S. pombe* harbors three TA(A/G)CG motifs, which are binding sites for the trans-acting factor Sap1 (Ghazvini *et al*. 1995). SAS1 counterparts in *S. osmophilus*, *S. japonicus*, *S. octosporus*, and *S. cryophilus* contain one, one, two, and two TA(A/G)CG motifs, respectively (Figure S14A).

In *S. pombe*, two 8-bp Atf1-binding sites (ATGACGTA) located in the donor region, called *s1* and *s2*, act redundantly with the RNAi pathway to promote the establishment of the donor region heterochromatin (Thon *et al*. 1999; Nickels *et al*. 2022). Our inspection of the donor region sequences of the other four fission yeast species showed that there are two, two, and four ATGACGTA sequences in the donor regions of *S. cryophilus*, *S. osmophilus*, and *S. octosporus*, respectively (Figure 5A). Interestingly, the ATGACGTA sequences closest to the donor cassettes in these three species share similar locations and surrounding sequences (Figure 5A and Figure S14C), suggesting that they have a common evolutionary origin and are maintained by purifying selection. The donor region of *S. japonicus* lacks such Atf1-binding sites, presumably because its positioning inside a centromere allows it to depend on neighboring centromeric sequences for heterochromatin establishment (Yu *et al*. 2013).

The donor regions of *S. octosporus* and *S. cryophilus* are known to harbor transposon remnants (Rhind *et al*. 2011; Tong *et al*. 2019). We found that transposon-related sequences also exist in the donor region of *S. osmophilus* (Figure 5A). We used the LTRs and the internal sequences (INTs) of full-length retrotransposons Tf1, Tf2, Tosmo1, Tosmo2, Tosmo3, Tcry1, and Tj1 as queries to perform BLASTN analysis of the donor regions of these three species. Based on the BLASTN results, we separately annotated sequences with similarities to LTRs and sequences with similarities to INTs (Figure 5A, Table S25, and Supplementary Files S1–S4). The annotated transposon-related sequences in the donor regions of *S. cryophilus*, *S. octosporus*, and *S. osmophilus* are most closely related to Tcry1, Tosmo1, Tosmo2, or Tosmo3 (Table S25). Multiple sequence alignment showed that INT2 and INT1 of *S. cryophilus* are substantially more similar to the internal sequence of Tcry1 than to internal sequences of Tosmo1, Tosmo2, and Tosmo3 and are likely to be two fragments of a retrotransposon that has suffered an internal deletion during degeneration (Figure 5C). INT4 of *S. osmophilus*, which is a nearly full-length internal sequence, is more similar to Tosmo2 than to other retrotransposons (Figure 5C). The close relationships of these INT sequences to full-length retrotransposons present in the same fission yeast species suggest that they result from retrotransposon insertions occurring after species divergence.

### The evolutionary trajectories of Cbp1 family proteins

Cbp1 family proteins are transposase-derived proteins known to be present in *S. pombe*, *S. octosporus*, and *S. cryophilus*, but not *S. japonicus* (Casola *et al*. 2008; Rhind *et al*. 2011; Upadhyay *et al*. 2017). There are three Cbp1 family proteins in *S. pombe*: Cbp1 (also known as Abp1), Cbh1, and Cbh2 (Irelan *et al*. 2001). They bind to the LTRs of retrotransposons in the *S. pombe* genome and repress the transcription of retrotransposons (Cam *et al*. 2008). In their absence, replication forks stall and collapse at LTRs (Zaratiegui *et al*. 2011). It has been proposed that the advent of Cbp1 family proteins in the common ancestor of *S. pombe*, *S. octosporus*, and *S. cryophilus* resulted in lower diversities of retrotransposons in these three fission yeast species compared to *S. japonicus* and a switch from a retrotransposon-rich centromere structure to a retrotransposon-free or -poor centromere structure (Rhind *et al*. 2011). Intrigued by their unique evolutionary origins and important roles in genome defense and maintenance, we decided to take advantage of the *S. osmophilus* genome to better understand the evolutionary origins and dynamics of Cbp1 family proteins.

*S. octosporus* and *S. cryophilus* each have six Cbp1 family proteins, encoded by six genes sharing synteny between these two species (Upadhyay *et al*. 2017) (Figures 6A and S15A–E). We found that *S. osmophilus* also has six Cbp1 family proteins and that their coding genes share synteny with genes encoding Cbp1 family proteins in *S. octosporus* and *S. cryophilus* (Figures 6A and S15A–E). We named these six proteins Cbt1–Cbt6 (Cbt stands for <u>Cb</u>p1 family domesticated transposase). They mostly share the same domain organization with *S. pombe* Cbp1 family proteins (Figure 6B). The exception is Cbt2, which lacks the N-terminal tandem helix-turn-helix (HTH) domains. For all three *S. pombe* Cbp1 family proteins, the eponymous catalytic triads of their DDE/D transposase domains are degenerated, with at least one of the three acidic residues mutated to a non-charged residue (Figure 6B). Interestingly, in four of the six Cbt proteins (Cbt2, Cbt3, Cbt4, and Cbt5), the catalytic triad residues are either DDE or DDD (Figures 6B and 6C), suggesting the possibility that these proteins may still retain some catalytic activities. Notable examples of domesticated transposases retaining catalytic activities include domesticated *piggyBac* transposases acting in programmed genome rearrangements in ciliates (Baudry *et al*. 2009; Cheng *et al*. 2010), vertebrate RAG proteins involved in V(D)J recombination (Huang *et al*. 2016), and two domesticated transposases involved in mating-type switching in *Kluyveromyces lactis* (Barsoum *et al*. 2010; Rajaei *et al*. 2014).

**Figure 6.**
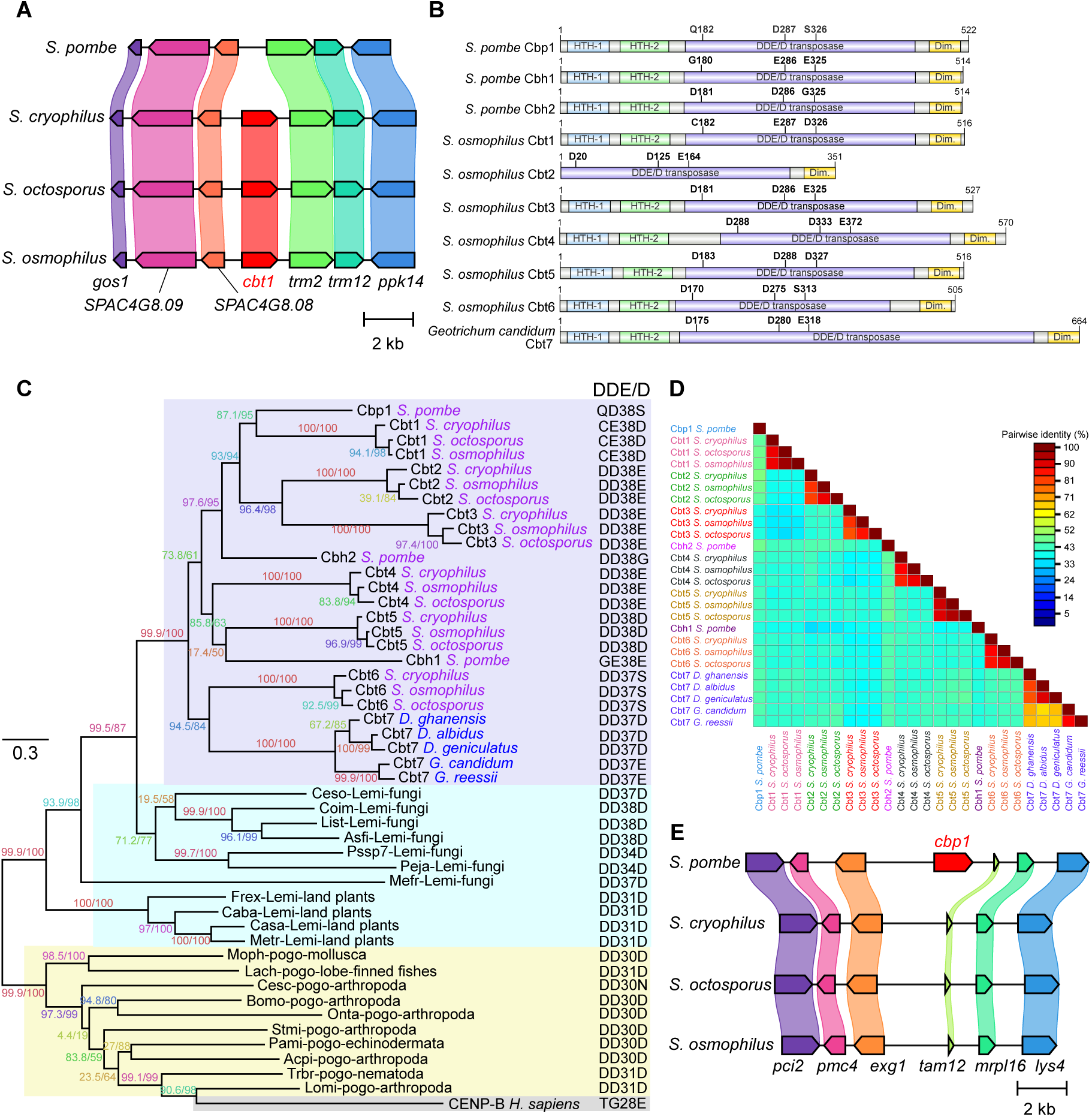
Evolutionary origins and trajectories of Cbp1 family proteins. (A) Diagram showing the local synteny at the genomic region containing the *cbt1* gene. Gene names shown are the names of the genes in *S. pombe*. The *S. cryophilus* orthologs of *S. pombe trm2* and *trm12* were manually annotated because they are missing in the gene annotation of the *S. cryophilus* reference genome. (B) Diagrams showing the domain organization of representative Cbp1 family proteins. Domain boundaries are based on the NMR structure of the N-terminal region of Cbp1 (Kikuchi *et al*. 2002), the crystal structure of the dimerization domain of human CENP-B (Tawaramoto *et al*. 2003), the AlphaFold-predicted structure of full-length Cbp1 (https://alphafold.ebi.ac.uk/entry/P49777) (Varadi *et al*. 2022), and sequence alignment. The three residues of the catalytic triad and their positions are shown on top of each domain diagram. These diagrams were generated using IBS 2.0 (Xie *et al*. 2022). (C) Maximum likelihood tree constructed using the amino acid sequences of Cbp1 family proteins of *Schizosaccharomyces* and *Dipodascaceae* (shaded in purple), human CENP-B (shaded in grey), and transposases of representative *Lemi* family transposons (shaded in blue) and *pogoR* family transposons (shaded in yellow). For Cbp1 family proteins, the names of the species are colored as follows: purple for *Schizosaccharomyces* species and blue for *Dipodascaceae* species. The tree was rooted using *pogoR* transposases (including human CENP-B) as outgroup. Branch labels are the SH-aLRT support value (%) and the UFBoot support value (%) calculated by IQ-TREE. The column with the header “DDE/ D” shows the catalytic triad residues and the number of amino acids between the last two residues of the catalytic triad. The accession numbers for the Cbp1 family proteins and human CENP-B are as follows: *S. pombe* Cbp1/SPBC1105.04c (NP_596460), *S. cryophilus* Cbt1/SPOG_05545 (XP_013022104), *S. octosporus* Cbt1/SOCG_02041 (XP_013015989), *S. osmophilus* Cbt1/SOMG_01668, *S. cryophilus* Cbt2/SPOG_05395 (XP_013022085), *S. octosporus* Cbt2/SOCG_00294 (XP_013018168), *S. osmophilus* Cbt2/SOMG_01142, *S. cryophilus* Cbt3/SPOG_05336 (XP_013024138), *S. octosporus* Cbt3/SOCG_02703 (XP_013016648), *S. osmophilus* Cbt3/SOMG_05008, *S. pombe* Cbh2/SPBC14F5.12c (NP_596738), *S. cryophilus* Cbt4/SPOG_05273 (XP_013022933), *S. octosporus* Cbt4/SOCG_00803 (XP_013018677), *S. osmophilus* Cbt4/SOMG_03711, *S. pombe* Cbh1/SPAC9E9.10c (NP_594583), *S. cryophilus* Cbt5/SPOG_05200 (XP_013025055), *S. octosporus* Cbt5/SOCG_00860 (XP_013018734), *S. osmophilus* Cbt5/SOMG_03771, *S. cryophilus* Cbt6/SPOG_05660 (XP_013023607), *S. octosporus* Cbt6/SOCG_04915 (XP_013019516), *S. osmophilus* Cbt6/SOMG_00850, *Geotrichum candidum* Cbt7 (KAF5117802), *Galactomyces reessii* Cbt7 (QVTG01000070.1 coordinates 42,175–44,130), *Dipodascus ghanensis* Cbt7 (JAKKCX010000005.1 coordinates 397,487–399,661), *Dipodascus geniculatus* Cbt7 (PPOV01000003.1 coordinates 283,093–285,156, minus strand), *Dipodascus albidus* Cbt7 (PPJE01000012.1 coordinates 37,750–39,864), and human CENP-B (NP_001801). Names and nucleotide sequences of *Lemi* family and *pogoR* family transposons were from Table S1 of Gao et al. 2020. Amino acid sequences of *Lemi* family transposases and *pogoR* family transposases were translated from the nucleotide sequences. (D) A color matrix showing the pair-wise amino acid identities between Cbp1 family proteins. This color matrix was generated using Sequence Demarcation Tool Version 1.2 (SDTv1.2) (Muhire *et al*. 2014). (E) Diagram showing the local synteny at the genomic region containing the *cbp1* gene. Gene names shown are the names of the genes in *S. pombe*.

Cbp1 family proteins originated from DNA transposons related to the *pogo* element of *Drosophila melanogaster* (Casola *et al*. 2008). The domestication of *pogo*-related transposons has occurred independently many times during evolution, with the most famous example being the emergence of the centromere-associated protein CENP-B in the common ancestor of mammals (Tudor *et al*. 1992; Casola *et al*. 2008; Mateo and González 2014; Dupeyron *et al*. 2020; Gao *et al*. 2020). Traditionally, *pogo*-related transposons are classified as a family belonging to the *Tc1/mariner* superfamily of DNA transposons (Plasterk *et al*. 1999). Two recent studies have elevated *pogo*-related transposons to the status of a superfamily sister to the *Tc1/mariner* superfamily (Dupeyron *et al*. 2020; Gao *et al*. 2020). The *pogo* superfamily has been divided into six main families: *Passer*, *Tigger*, *pogoR*, *Lemi*, *Mover*, and *Fot/Fot-like* (Gao *et al*. 2020). In vertebrates, four domesticated genes (*CENPB*, *TIGD3*, *TIGD4*, and *TIGD6*) are derived from *pogoR* family transposons, six domesticated genes (*CENPBD1*, *JRK*, *JRKL*, *TIGD2*, *TIGD5*, and *TIGD7*) are derived from *Tigger* family transposons, and two domesticated genes (*POGK* and *POGZ*) are derived from *Passer* family transposons (Gao *et al*. 2020). The origins of domesticated *pogo* superfamily transposons in fungi have not received the same level of scrutiny.

We used the 21 fission yeast Cbp1 family proteins as queries to perform TBLASTN searches against a collection of 1014 representative nucleotide sequences of *pogo* superfamily transposons curated by Gao et al. 2020. The top 10 hits for each query overlap extensively across the 21 queries. There are a total of 20 transposons in the top 10 hits for all queries (Table S26). They are exclusively *Lemi* family transposons present in fungal species, including one *Saccharomycotina* species (*Lipomyces starkeyi*) and more than 10 *Pezizomycotina* species belonging to the classes of *Eurotiomycetes*, *Dothideomycetes*, and *Leotiomycetes*. The amino acid identities between queries and top hits are in the range of 32–39% and similarities are found throughout entire query sequences (Table S26). Thus, we conclude that Cbp1 family proteins likely originated from *Lemi* family transposons.

We also used the 21 fission yeast Cbp1 family proteins as queries to perform BLASTP searches against the NCBI nr database excluding *Schizosaccharomyces*. Consistent with the TBLASTN results described above, transposases of *Lemi* family transposons are among the high-scoring hits (Table S27). Interestingly, for a majority of the queries, the highest scoring hit is either KAF5117802.1 or KAF7497890.1, two nearly identical proteins (amino acid identity 99.4%) from the *Saccharomycotina* species *Geotrichum candidum* (teleomorph *Galactomyces candidus*) (Table S27). These two proteins are from two different strains (LMA-317 and LMA-40) of *G. candidum* (Perkins *et al*. 2020). BLASTP searches using these two *G. candidum* proteins as queries to search against the NCBI nr database excluding *G. candidum* identified fission yeast Cbp1 family proteins as the highest scoring hits and transposases of known *Lemi* family transposons as lower-ranking hits. Thus, fission yeast Cbp1 family proteins and these two *G. candidum* proteins share higher similarities than they do with transposases of known *Lemi* family transposons.

To determine whether KAF5117802.1 and KAF7497890.1 are proteins encoded by transposons, we retrieved their coding sequences and flanking regions from the genome assemblies of LMA-317 and LMA-40. We could not find terminal inverted repeats (TIRs), the hallmark of DNA transposons, in nearby flanking sequences. Sequence alignment showed that in LMA-317 and LMA-40, not only the coding sequences but also the flanking regions containing multiple neighboring genes are nearly identical (nucleotide identity 97.8% in a 24-kb region). Thus, these two proteins are encoded by the same gene locus in the two *G. candidum* strains. No additional copies of this gene could be found in the two genome assemblies. The lack of TIRs, the existence as a single-copy gene, and the identical genome locations in two divergent strains all suggest that this gene is a domesticated transposon.

Using KAF5117802.1 as query to perform TBLASTN search against the NCBI whole-genome shotgun contigs (wgs) database led to the identification of closely related homologs in four fungal species belonging to the same family (*Dipodascaceae*) as *G. candidum*: *Dipodascus albidus*, *Dipodascus geniculatus*, *Dipodascus ghanensis*, and *Galactomyces reessii*. Similarities between the query and hits were found throughout the length of the query and the amino acid identities ranged from 62% to 85%. The local synteny is largely conserved between *G. candidum* and these four species (Figure S15F). *D. albidus* and *D. geniculatus* are estimated to have diverged about 112 million years ago from *G. candidum* (Shen *et al*. 2020). Thus, the gene encoding KAF5117802.1 has been residing at the same locus for over 100 million years. Together, our observations suggest that this gene originated from a transposon domestication event occurring more than 100 million years ago. Because it encodes a protein with high similarity to fission yeast Cbp1 family proteins, we named this gene *CBT7*.

We performed a phylogenetic analysis using the amino acid sequences of 21 fission yeast Cbp1 family proteins, five Cbt7 proteins from *Dipodascaceae* species, human CENP-B, and 21 transposases of representative transposons belonging to *Lemi* and *pogoR* families (Figure 6C). Consistent with the BLAST results, fission yeast Cbp1 family proteins and *Dipodascaceae* Cbt7 proteins form a highly supported monophyletic clade, which we hereafter refer to as the Cbp1 family clade. Transposases of *Lemi* family transposons from the *Saccharomycotina* species *Lipomyces starkeyi* and several *Pezizomycotina* species form a sister clade to the Cbp1 family clade. These two clades are nested within a larger monophyletic clade that includes transposases of other *Lemi* family transposons. Thus, phylogenetic analysis lends support to the idea that Cbp1 family proteins are derived from *Lemi* family transposases. As expected, *pogoR* family transposases, which form a monophyletic group including human CENP-B, are sister to *Lemi* family transposases and Cbp1 family proteins.

Within the Cbp1 family clade, the 18 proteins from *S. osmophilus*, *S. octosporus*, and *S. cryophilus* form six highly supported monophyletic clusters, corresponding to Cbt1–Cbt6 (Figure 6C). The phylogenetic relationships between the three proteins in each of these six clusters are always congruent with the species tree. The catalytic triad residues and the number of amino acids between the last two residues of the catalytic triad are strictly conserved within each cluster (Figure 6C). Pairwise identities of proteins within each cluster are in the range of 86%–96% for the *S. osmophilus*–*S. octosporus* comparison, in the range of 82%–92% for the *S. osmophilus*–*S. cryophilus* comparison, and in the range of 79%–91% for the *S. octosporus*–*S. cryophilus* comparison (Figures 6D). These values are largely in line with those of single-copy orthologs conserved across all five fission yeasts species (Figure S5C), suggesting that since the divergence of *S. osmophilus*, *S. octosporus*, and *S. cryophilus* about 29 million years ago, Cbt1–Cbt6 have evolved at rates similar to conserved proteins under purifying selection. Together, these observations indicate that the domestication event(s) leading to the emergence of Cbt1– Cbt6 must have occurred earlier than the divergence of these three species. Furthermore, the conservation of Cbt1–Cbt6 proteins suggests that the presence of full-length retrotransposons in *S. osmophilus* and *S. cryophilus* but not *S. octosporus* cannot be attributed to interspecific differences in Cbp1 family proteins.

Cbt7 proteins from the five *Dipodascaceae* species also form a monophyletic cluster (Figure 6C). Branches within the Cbt7 cluster have longer lengths than branches within Cbt1– Cbt6 clusters, consistent with longer species divergence time (112 million years vs. 29 million years). The seven Cbt clusters and the three *S. pombe* proteins reside on 10 long branches indicative of substantial divergence. Indeed, pairwise identities between any two proteins not affiliated with the same long branch are below 50% (Figure 6D). Intriguingly, fission yeast Cbp1 family proteins form a paraphyletic group, with the *Dipodascaceae* Cbt7 cluster nested in it. The fission yeast Cbt6 proteins are sister to the *Dipodascaceae* Cbt7 proteins. Such phylogenetic relationships suggest that horizontal evolutionary events are likely responsible for the presence of Cbp1 family proteins in both *Dipodascaceae* species and *Schizosaccharomyces* species, which are estimated to have diverged about 563 million years ago (Shen *et al*. 2020).

The three *S. pombe* Cbp1 family proteins do not group together by themselves. Instead, Cbp1 is sister to the Cbt1 cluster and Cbh2 is sister to a group including Cbp1 and Cbt1–Cbt3 clusters. Cbh1 is sister to the Cbt5 cluster but this grouping has poor support. Because genes encoding the three *S. pombe* Cbp1 family proteins do not share synteny with genes encoding Cbt1–Cbt6 proteins (Upadhyay *et al*. 2017) (Figures 6E and S15G–H), the relationships between Cbp1 and Cbt1 and between Cbh1 and Cbt5 are unlikely to be orthologous. The intermingling of *S. pombe* Cbp1 family proteins with Cbt1–Cbt5 clusters suggests that, if the three *S. pombe* Cbp1 family proteins arose through gene duplications, the duplication events occurred before the divergence of *S. pombe* from the common ancestor of *S. osmophilus*, *S. octosporus*, and *S. cryophilus* about 108 million years ago. Similarly, if Cbt1–Cbt6 proteins resulted from gene duplications, most duplications events probably occurred in the common ancestor of the four fission yeast species.

We propose two possible evolutionary scenarios to explain the phylogenetic pattern of the Cbp1 family clade. In the first scenario, all proteins belonging to this clade originated from a single transposon domestication event occurring more than 100 million years ago in the common ancestor of the four *Schizosaccharomyces* species. A series of gene duplications resulted in nine or more paralogs dispersed in the genome. One paralog underwent horizontal transfer into the common ancestor of the five *Dipodascaceae* species. Subfunctionalization or neofunctionalization of the paralogs was accompanied by rapid evolution (Lynch and Conery 2000; Scannell and Wolfe 2008), contributing to the long branch lengths. After the divergence of *S. pombe* from the common ancestor of *S. osmophilus*, *S. octosporus*, and *S. cryophilus*, independent gene loss events in the two lineages resulted in the remaining paralogs being completely non-overlapping between the two lineages. In the second scenario, multiple related but nonidentical *Lemi* family transposons invaded an ancestral *Schizosaccharomyces* species more than 100 million years ago, and one of these transposons also invaded an ancestral *Dipodascaceae* species at around the same time. Independent domestication events happened in the two ancestral species. In the ancestral *Schizosaccharomyces* species, multiple domestication events led to dispersed paralogous genes. Later, gene loss events occurred as in the first scenario, resulting in the phylogenetic pattern we see today.

### Mitogenome of *S. osmophilus* and comparative analysis of fission yeast mitogenomes

Mitochondria possess a genome termed the mitogenome. In *S. pombe*, the mitogenome is indispensable for viability in the wild-type genetic background (Chiron *et al*. 2007; Li *et al*. 2019). The analysis of intraspecific mitogenome diversity of *S. pombe* has offered insights into the evolutionary history of this species (Tao *et al*. 2019). An interspecific comparison of the mitogenomes of *S. pombe*, *S. octosporus*, and *S. japonicus* has been conducted (Bullerwell *et al*. 2003). The mitogenome of *S. cryophilus* has been sequenced but has not been subjected to in-depth analysis (Rhind *et al*. 2011). Here, we assembled the complete circular-mapping mitogenome of the *S. osmophilus* type strain CBS 15793^T^ (Figure 7A and Table S28). We annotated it and performed comparative analysis of the mitogenomes of five fission yeast species.

**Figure 7.**
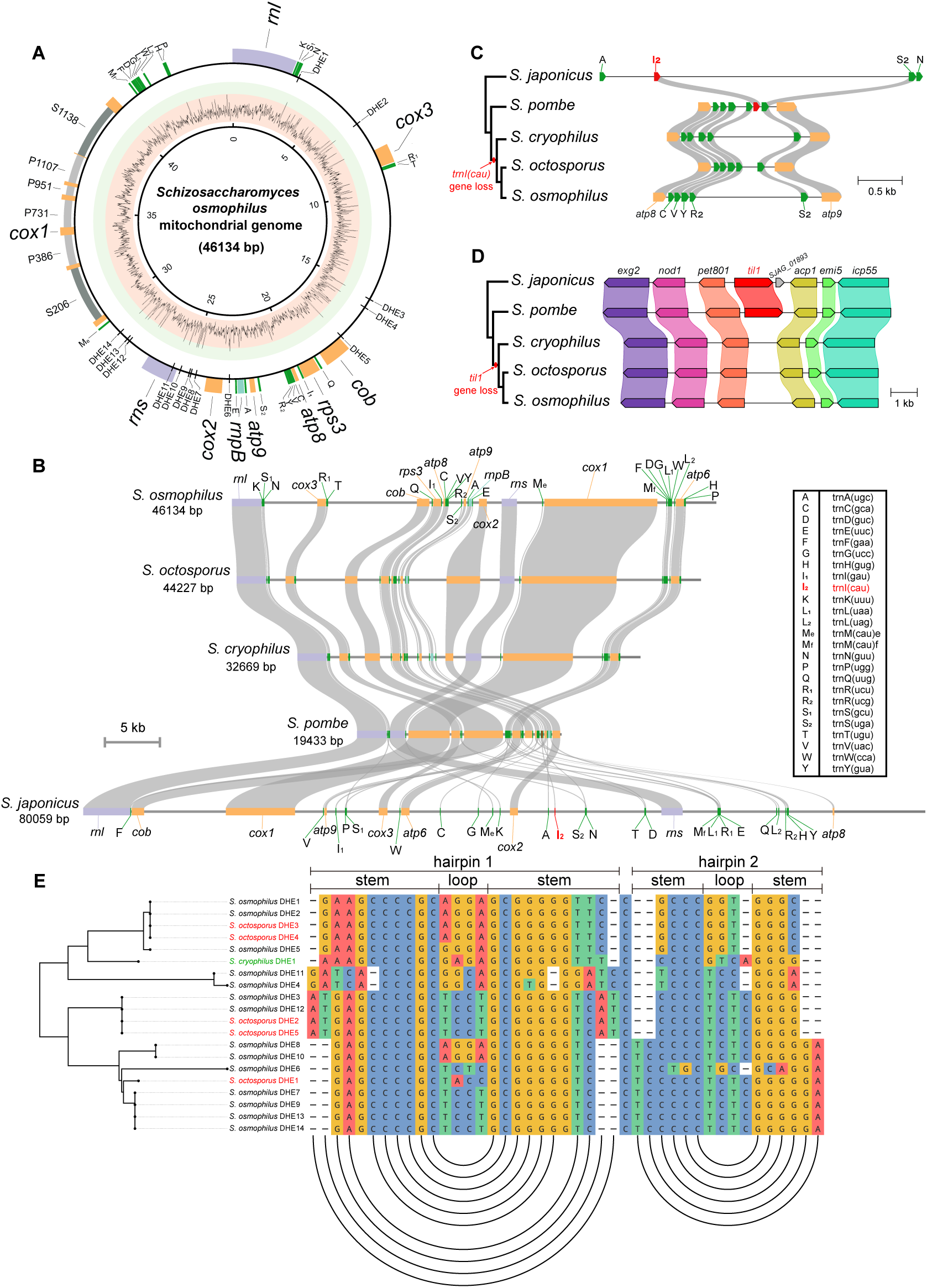
Mitogenome of *S. osmophilus* and comparative analyses of fission yeast mitogenomes. (A) Graphic map of the circular-mapping mitogenome of *S. osmophilus*. Genes are shown in the outer circle and are colored as follows: protein-coding genes, yellow; *rnpB*, blue; rRNA genes, purple; tRNA genes, green. The names of tRNA genes are abbreviated and their full names are shown at the right side of panel B. Group I introns and group II introns are colored light grey and dark grey, respectively. GC content in 35-bp sliding windows is shown as a line plot in an inner circle, where red background denotes the GC content range of 0–70% and green background denotes the GC content range of 70– 100%. (B) Gene order conservation between the mitogenomes of five fission yeast species. Genes are colored as in (A), except that the tRNA gene *trnI(cau)* (abbreviated as I2) is highlighted in red. The transcription orientation is from left to right for all genes except the *atp8* gene and three tRNA genes in the *S. japonicus* mitogenome (Bullerwell *et al*. 2003). For brevity, introns are not shown. (C) Local synteny of the mitogenome region containing the *trnI(cau)* gene (abbreviated as I2). A schematic species tree is shown at left. (D) Local synteny of the nuclear genome region containing the *til1* gene. A schematic species tree is shown at left. (E) Sequence alignment of DHEs in the mitogenomes of *S. osmophilus*, *S. octosporus*, and *S. cryophilus*. Below the alignment is an arc diagram depicting the predicted secondary structures of DHEs. In the arc diagram, each arc represents a base pair and connects the positions of two base-pairing nucleotides. At left is a neighbor-joining tree constructed using the bionj function of the APE package based on a distance matrix generated by the dist.gene function of APE (Paradis *et al*. 2004).

The mitogenome of the *S. osmophilus* type strain is 46134 bp long and contains eight protein-coding genes (*atp6*, *atp8*, *atp9*, *cob*, *cox1*, *cox2*, *cox3*, and *rps3*), the RNaseP RNA gene (*rnpB*), two rRNA genes (*rnl* and *rns*), and 24 tRNA genes (Figure 7A). All 35 genes are encoded on the same DNA strand. Such gene content and gene orientations are exactly the same as those of the reference mitogenomes of *S. octosporus* and *S. cryophilus* (Figure 7B). Moreover, the 35 genes show perfect synteny between these three species (Figure 7B). The size differences between the mitogenomes of these three species are mainly due to differences in the lengths of introns and intergenic regions (Figure S16A). Partial gene synteny conservation can be observed between these three species and *S. pombe*, whereas the *S. japonicus* mitogenome shares little synteny with the mitogenomes of the other four fission yeast species (Figure 7B). A phylogenetic tree generated using mitochondrial genes has the same topology as the species tree generated using nuclear genes, with *S. octosporus* being the closest relative of *S. osmophilus* (Figure S16B).

We analyzed the codon usage of protein-coding genes in the mitogenomes of five fission yeast species (Figure S16C). As has been observed before for many *Ascomycota* species (Bullerwell *et al*. 2003; Kamatani and Yamamoto 2007; Procházka *et al*. 2012), in all five species, there exists a strong preference for A and T at the third position of redundant codons. The exceptions are the phenylalanine codons and the tryptophan codons. The two phenylalanine codons TTT and TTC are used at roughly equal frequencies. This has been explained by a tendency to avoid frameshift-prone polyU in mRNAs (Bullerwell *et al*. 2003). Among the two tryptophan codons TGG and TGA, TGG is strongly favored, perhaps due to the lack of a tRNA that can efficiently decode the TGA codon (Bullerwell *et al*. 2003).

Similar to the situation in *S. octosporus* (Bullerwell *et al*. 2003), we found that non-intronic protein-coding genes in the mitogenomes of *S. osmophilus* and *S. cryophilus* lack the ATA codon for isoleucine (Figure S16C). In contrast, the ATA codon is used many times by non-intronic protein-coding genes in the mitogenomes of *S. pombe* and *S. japonicus*. In these two species, the ATA codon is decoded by a tRNA encoded by the mitochondrial tRNA gene *trnI(cau)* (abbreviated as I_2_ in Figures 7A–7C). This tRNA is only functional when the cytidine at the wobble position is converted to lysidine (2-lysyl cytidine), which can base pair with adenosine. This tRNA modification is catalyzed by the enzyme tRNA(Ile)-lysidine synthase (Xavier *et al*. 2012), which is encoded by the nuclear gene *til1* in *S. pombe* and *S. japonicus*. Interestingly, we found that *S. octosporus*, *S. osmophilus*, and *S. cryophilus* all lack the *trnI(cau)* gene in their mitogenomes and the *til1* gene in their nuclear genomes (Figures 7C and 7D). Thus, both genes were probably lost before the divergence of these three species about 29 million years ago. The *til1* gene is an essential gene in *S. pombe* (Kim *et al*. 2010; Hayles *et al*. 2013). One possible evolutionary scenario is as follows: the *til1* gene in the ancestral nuclear genome was partially inactivated by mutation. Mitochondrial functions were compromised due to the insufficient modification of the tRNA that decodes the ATA codon. Selective pressure led to the replacement of the ATA codon in the mitogenome by other isoleucine codons. The *til1* gene and the *trnI(cau)* gene thus became dispensable and eventually lost.

Mitochondrial introns are mobile elements and some of them harbor one or more open reading frames for intron-encoded proteins (IEPs) (Lang *et al*. 2007). There are six introns in the mitogenome of the *S. osmophilus* type strain, including four group I introns and two group II introns (Figure 7A). They all reside in the *cox1* gene. We named them using a recently proposed nomenclature that denotes the intron type with a uppercase letter (P for group I introns and S for group II introns) and denotes the location in the host gene with a number, which is the corresponding position in the equivalent gene of the *Pezizomycotina* species *Tolypocladium inflatum* (Zhang and Zhang 2019). This nomenclature is designed to facilitate comparisons between different mitogenomes. Thus, we also applied this nomenclature to mitochondrial introns of other four fission yeast species (Table S29 and Figure S16D) (Bullerwell *et al*. 2003; Tao *et al*. 2019). Five of the six *S. osmophilus cox1* introns (S206, P386, P731, P1107, and S1138), which all encode proteins, reside at locations where introns have been found in other fission yeast species (Figure S16D). The other *S. osmophilus cox1* intron (P951) does not encode a protein and is at a location where introns are not known to exist in other fission yeast species (Figure S13D).

Inspection of all known fission yeast mitochondrial introns revealed that introns at the same location but in different species not only share the same intron type (group I or group II), but also mostly share the same subgroup affiliation for group I introns (Figure S16D). In addition, IEPs encoded by introns at the same location are more similar to each other than to IEPs encoded by introns at other locations (Figures S16E and S16F). These observations suggest that fission yeast mitochondrial introns at the same location tend to share a common evolutionary origin. This is consistent with the “position class” concept proposed based on an analysis of *cox1* group I introns in fungi and other eukaryotes (Férandon *et al*. 2010).

Unexpectedly, *S. osmophilus* cox1P386 IEP shares substantially higher homology to *S. cryophilus* cox1P386 IEP (pairwise identity 80.2%) than to *S. octosporus* cox1P386 IEP (pairwise identity 53.8%) (Figure S16E). The pairwise identities between cox1P731 IEPs are also incongruent with the species tree, with the *S. octosporus* protein being more similar to the *S. cryophilus* protein than to the *S. osmophilus* protein (Figure S16E). We hypothesized that such patterns may result from interspecific introgression or horizontal transfer. Sliding-window analysis showed that sequences that may have undergone introgression or horizontal transfer are limited to within the cox1P386 intron and the cox1P731 intron (Figure S17A).

All proteins encoded by fission yeast group I introns are double-motif LAGLIDADG homing endonucleases (LHEs), which contain two LAGLIDADG domains. Previously we showed that half (7/14) of the LHEs in *S. pombe*, *S. octosporus*, *S. cryophilus*, and *S. japonicus* suffer inactivating mutations at catalytic residues and thus no longer possess the ability to generate DNA double-strand breaks that initiate intron mobilization (Tao *et al*. 2019). All three LHEs in *S. osmophilus* contain inactivating mutations at catalytic residues (Figure S17B). Interestingly, the prevalence of this type of LHE degeneration varies depending on intron location, with cox1P731 introns nearly always harbor one active LHE and cox1P720 introns never harbor an active LHE (Figure S17B). Such a position dependence of the extent of LHE degeneration suggests that introns descending from different origins tend to have different evolutionary trajectories.

Double-hairpin elements (DHEs) are potentially mobile elements first found in the mitogenomes of several *Chytridiomycota* fungal species (Paquin and Lang 1996; Paquin *et al*. 1997, 2000). The only *Ascomycota* species where DHEs have been found is *S. octosporus* (Bullerwell *et al*. 2003). There are five DHEs in the *S. octosporus* mitogenome. Four of them are located in intergenic regions and one of them resides in the *rnl* gene (Bullerwell *et al*. 2003). Our inspection of the mitogenomes of *S. osmophilus* and *S. cryophilus* led to the identification of 14 DHEs in *S. osmophilus* and one DHE in *S. cryophilus* (Figures 7A and 7E). Except for *S. osmophilus* DHE11, which is located in the *rns* gene, the other DHEs of *S. osmophilus* and *S. cryophilus* are located in intergenic regions. The DHEs of *S. osmophilus* and *S. cryophilus* have the potential to form similar double-hairpin structures as the five *S. octosporus* DHEs (Figure 7E). Remarkably, *S. osmophilus* DHE1 and DHE2 are identical in sequence to *S. octosporus* DHE3 and DHE4, *S. osmophilus* DHE3 and DHE12 are identical in sequence to *S. octosporus* DHE2 and DHE5, and *S. osmophilus* DHE7, DHE9, DHE13, and DHE14 only differ from *S. octosporus* DHE1 by two nucleotides in a loop region (Figure 7E). Such cross-species conservation patterns suggest that either the sequences of DHEs have been strictly maintained by purifying selection after the divergence of *S. octosporus* and *S. osmophilus* about 16 million years ago, or perhaps more likely, DHEs have recently undergone introgression or horizontal transfer across species barriers. As *S. osmophilus* has the highest abundance of DHEs among fission yeast species, it is an attractive model for further studying the evolution and activities of DHEs.

## Conclusion

In this study, we assembled a high-quality chromosome-level genome for the type strain of the fission yeast species *S. osmophilus* and comprehensively annotated the genes in this genome. Cross-species comparative analyses reported here demonstrate the usefulness of this genome for understanding how fission yeast genomes have evolved. We expect that this reference genome for *S. osmophilus* will facilitate future research using this species and serve as a valuable resource to support comparative genomic and population genomic studies.

## Acknowledgments

We thank Yan-Hui Xu for identifying the misannotation of Tcry1-1. We thank Fang Suo and Wen Li for helps with bioinformatic analyses and suggestions on the manuscript. N.R. is supported by a grant from the NIH (GM134300). A.L.P. is supported by funding to Robin Allshire from the Wellcome Trust (200885, 224358). Wellcome Centre for Cell Biology is supported by core funding from the Wellcome Trust (203149). M.B.-H. is supported by the Deutsche Bundesstiftung Umwelt DBU (German Federal Environmental Foundation) (34053/01-32). L.-L.D. is supported by grants from the Ministry of Science and Technology of China and the Beijing municipal government.

---

**Figure S1.**
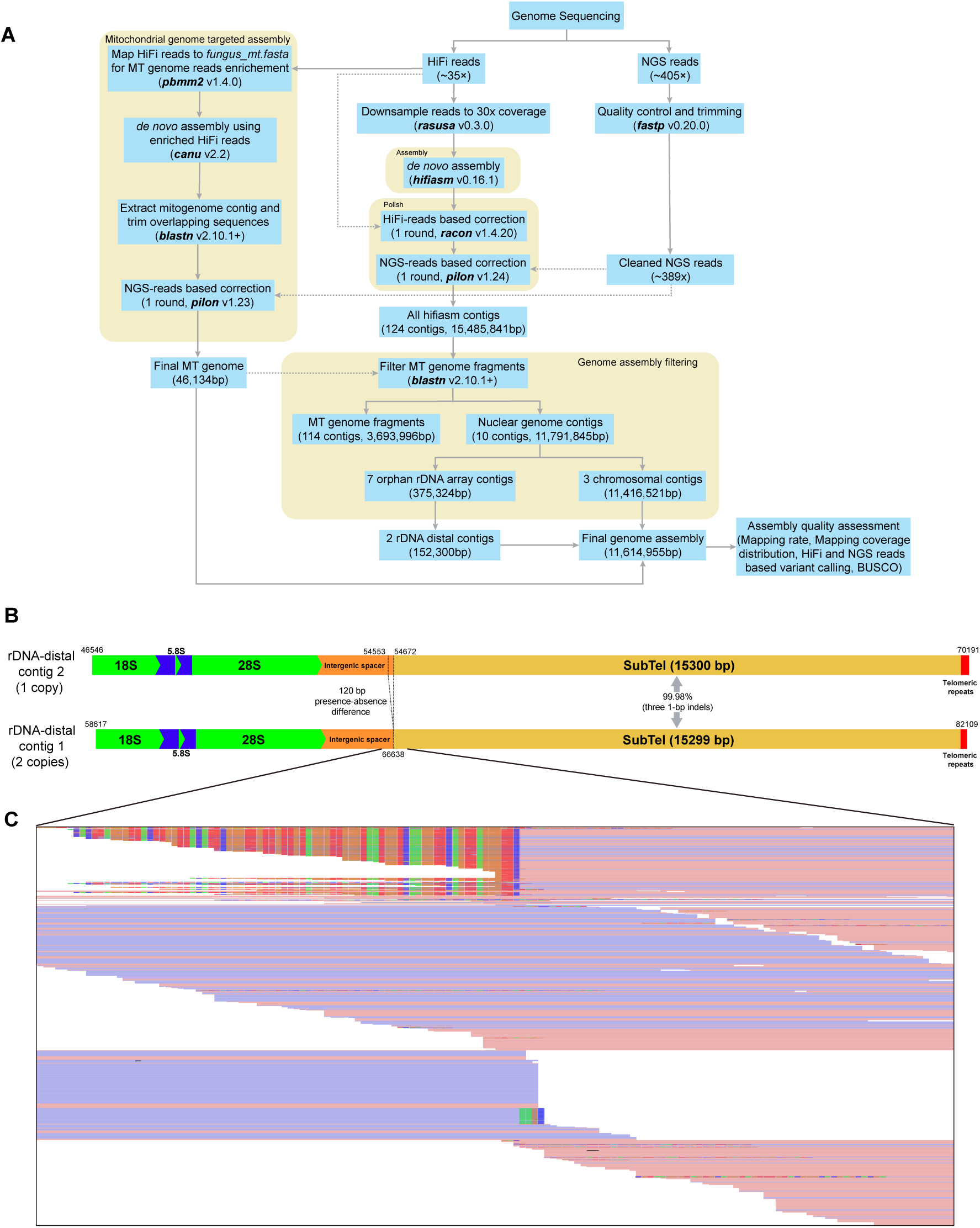
Assembling the genome of the *S. osmophilus* type strain CBS 15793^T^. (A) Workflow of genome assembly. (B) Diagrams of the two rDNA-distal contigs. (C) IGV screenshot showing the result of mapping all paired-end Illumina reads to the rDNA-distal contig-1. Reads that show perfect mapping at the rDNA/non-rDNA junction are about twice as many as reads that are soft-clipped at the rDNA/non-rDNA junction. The soft-clipped portions match the sequence at the rDNA/non-rDNA junction of the rDNA-distal contig-2.

**Figure S2.**
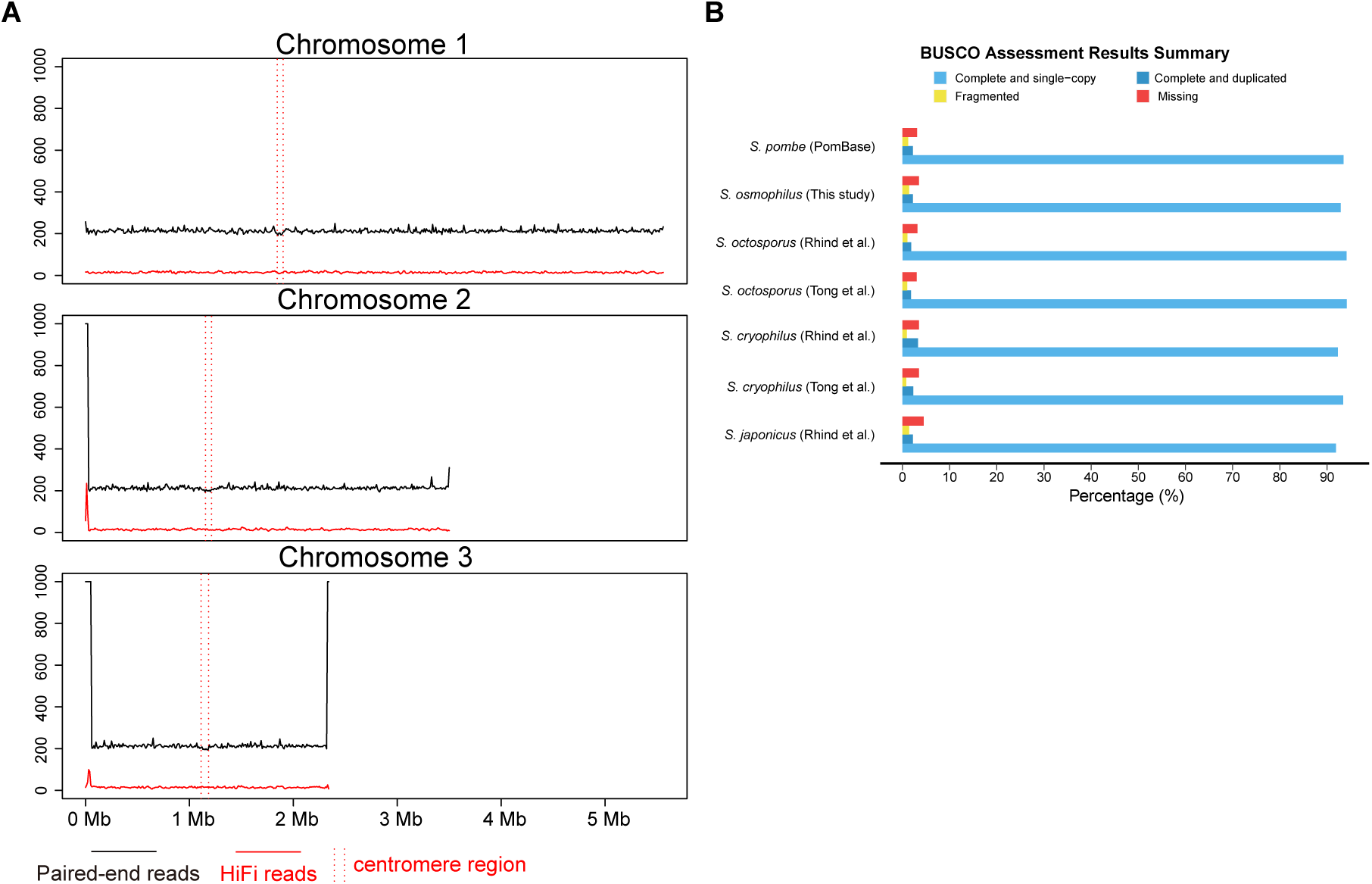
Quality assessment of the genome assembly. (A) Sliding-window analysis (10-kb window) of the depth coverage of PacBio HiFi reads (red line) and Illumina paired-end reads (black line) along the chromosomal contigs. (B) BUSCO completeness assessment of the *S. osmophilus* genome assembly and six published genomes of *S. pombe*, *S. octosporus*, *S. cryophilus*, and *S. japonicus*.

**Figure S3.**
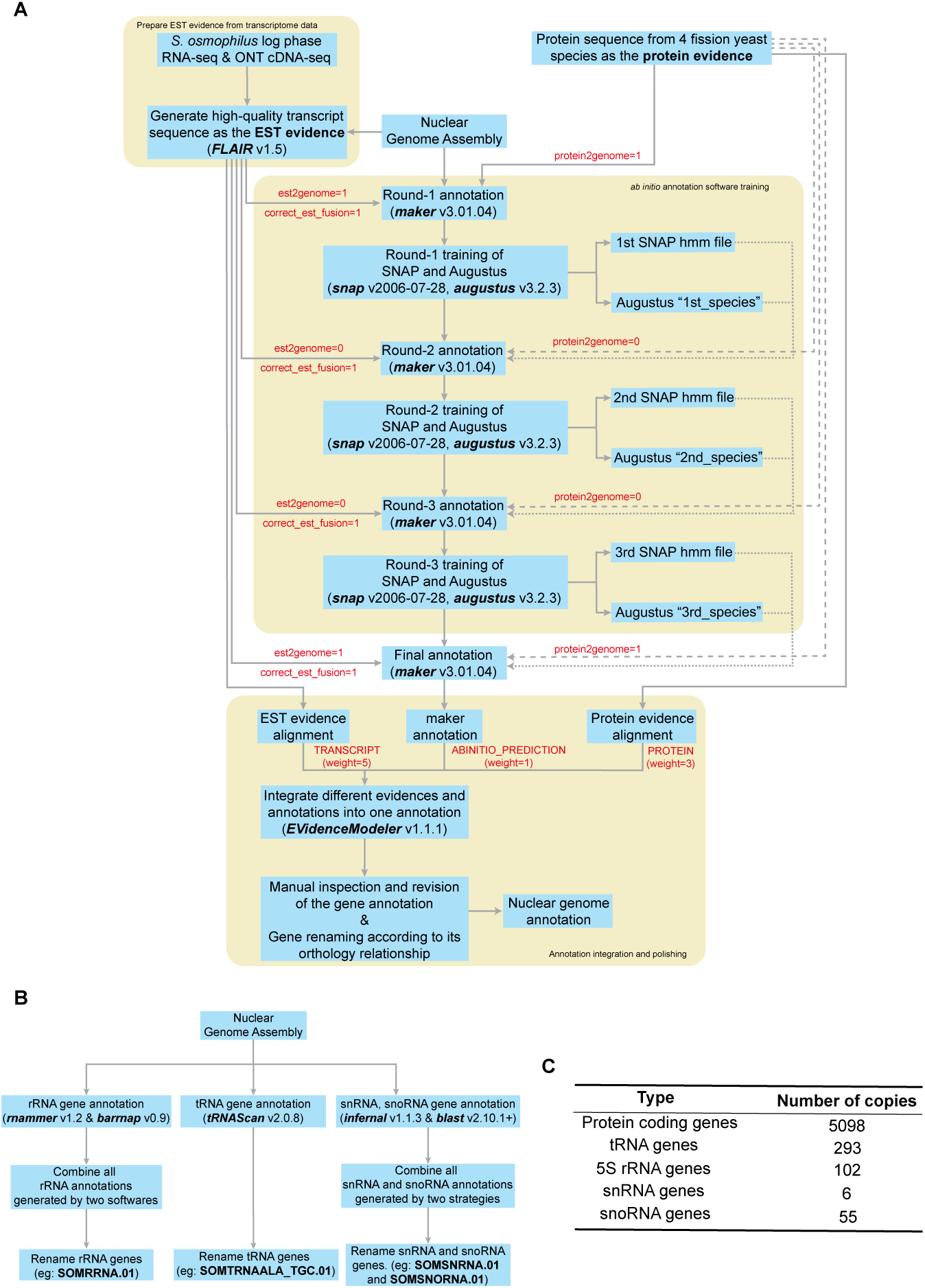
Annotating the genes in the genome assembly of the *S. osmophilus* type strain CBS 15793^T^. (A) Workflow of annotating protein-coding genes in the nuclear genome. (B) Workflow of annotating non-coding RNA genes in the nuclear genome. (C) Summary of the annotated genes in the nuclear genome.

**Figure S4.**
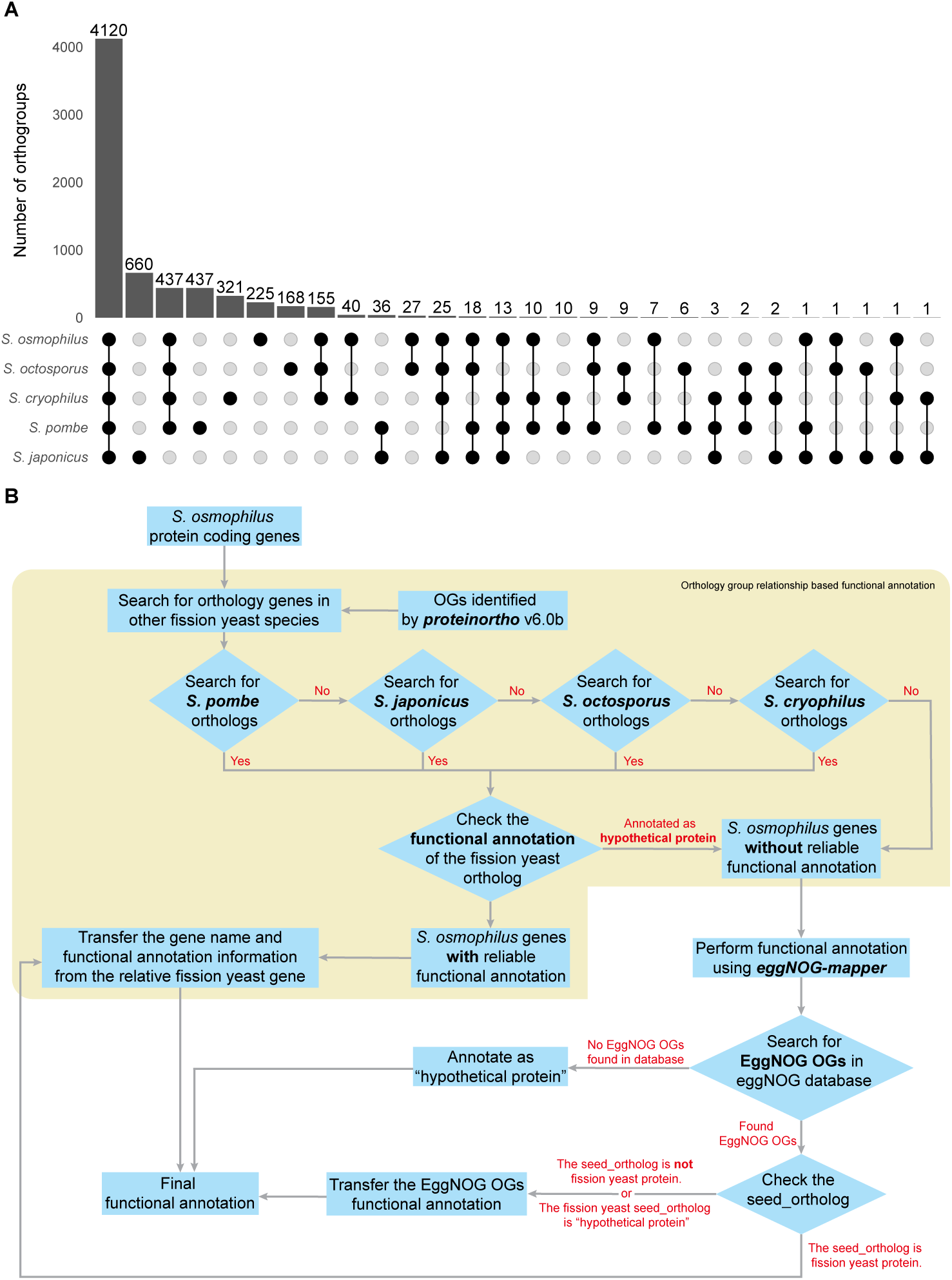
Orthogroup analysis and functional annotation of protein-coding genes. (A) UpSet plot showing the numbers of species-specific orthogroups and orthogroups shared by more than one fission yeast species. **(B)** Workflow of functionally annotating protein-coding genes in the nuclear genome.

**Figure S5.**
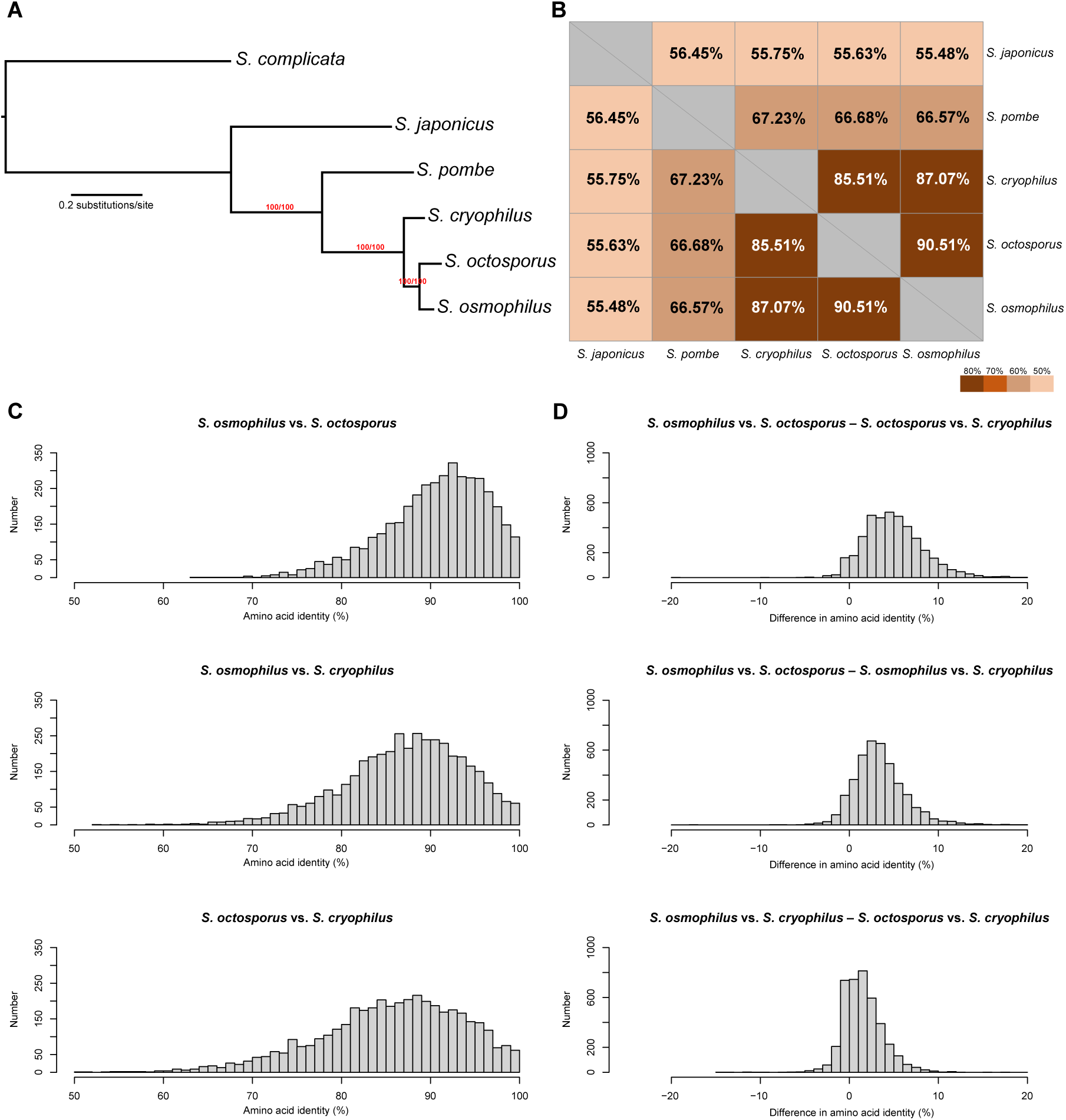
Species phylogeny analysis and evolutionary rate analysis. (A) Maximum likelihood species tree constructed using the concatenation super-matrix of amino acid sequences of 1060 “complete and single-copy” BUSCO genes that present in all fission yeast species and the outgroup species *S. complicata*. The tree is rooted using *S. complicata* as the outgroup. Branch labels are the SH-aLRT support value (%) and the UFBoot support value (%) calculated by IQ-TREE. (B) Average amino acid percentage identities of fission yeast species pairs calculated using 4085 1:1:1:1:1 single-copy orthologs defined in this study. Pairwise amino acid percentage identities were calculated using Sequence Demarcation Tool Version 1.0 (Linux version). The heatmap was generated using the R package pheatmap (https://github.com/raivokolde/pheatmap). (C) Histogram illustrating the distribution of pairwise amino acid identities of the 4085 single-copy orthologs. (D) Histogram illustrating the distribution of differences in pairwise amino acid identities of the 4085 single-copy orthologs.

**Figure S6.**
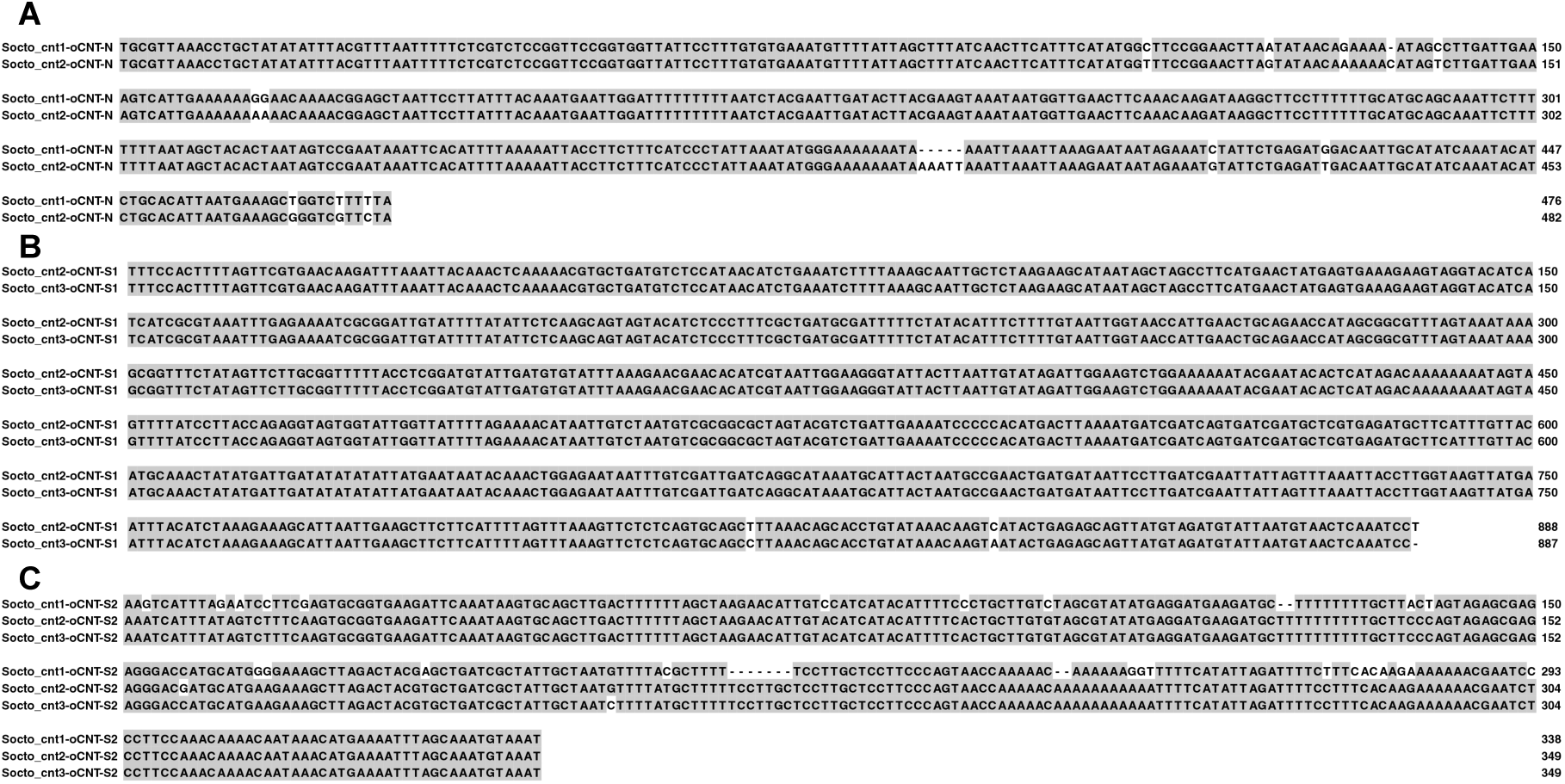
Sequence alignments of the oCNT-N, oCNT-S1, and oCNT-S2 repeats of *S. octosporus*. Nucleotides identical to the consensus are shaded in gray. (A) Alignment of the nucleotide sequences of oCNT-N. (B) Alignment of the nucleotide sequences of oCNT-S1. (C) Alignment of the nucleotide sequences of oCNT-S2.

**Figure S7:**
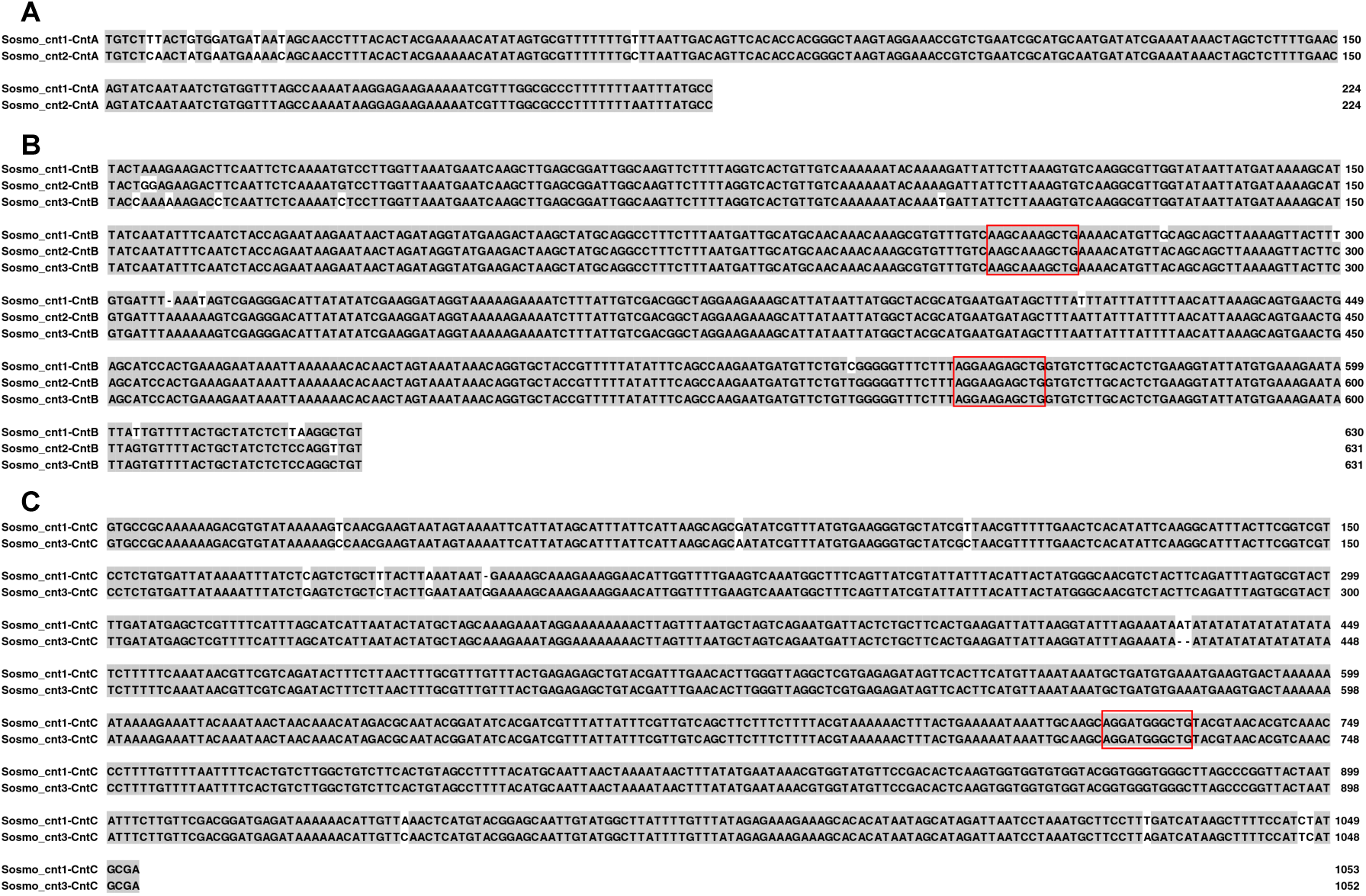
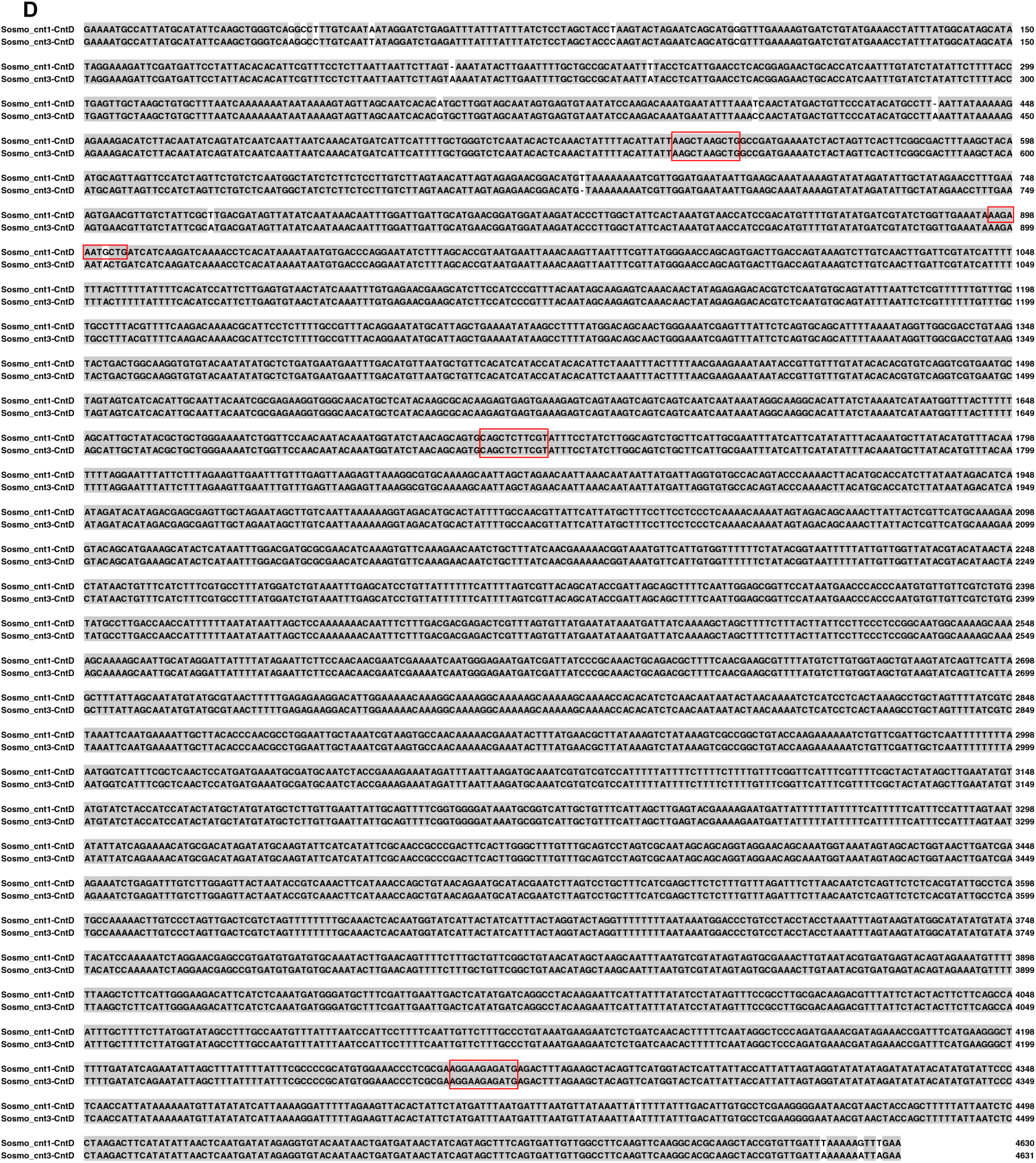
Sequence alignments of the *cnt* repeats of *S. osmophilus* and the occurrences of the 11-bp motif (boxed) in these repeats. Nucleotides identical to the consensus are shaded in gray. (A) Alignment of the nucleotide sequences of CntA. (B) Alignment of the nucleotide sequences of CntB. (C) Alignment of the nucleotide sequences of CntC. (D) Alignment of the nucleotide sequences of CntD.

**Figure S8.**
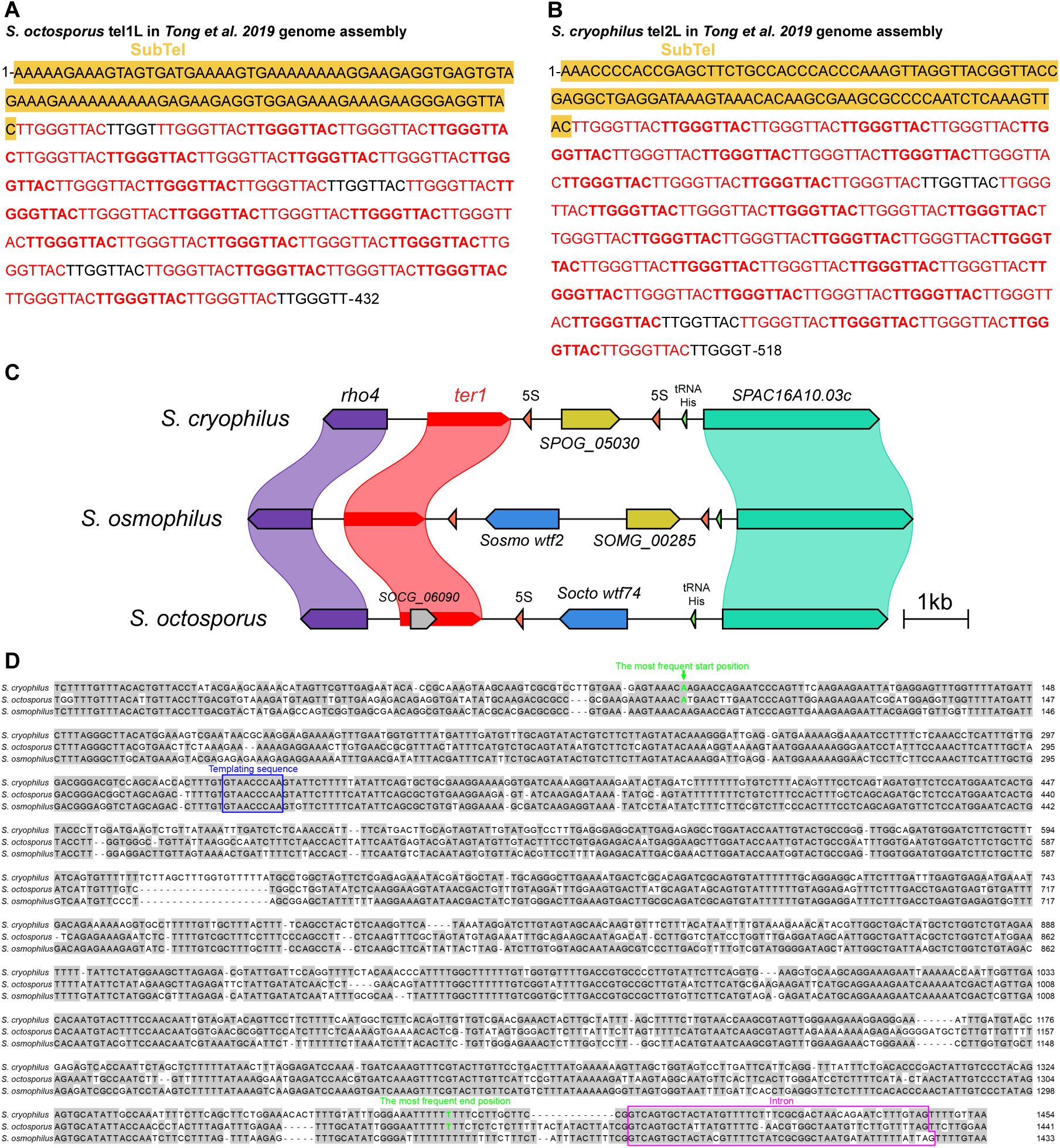
Comparative analysis of telomeric repeats and telomerase RNA genes in *S. osmophilus*, *S. octosporus*, and *S. cryophilus*. (A and B) Sequence of a telomere in the PacBio sequencing-based *S. octosporus* genome assembly (A) and a telomere in the PacBio sequencing-based *S. cryophilus* genome assembly (B) (Tong *et al*. 2019). The sequence juxtaposed to telomeric repeats are shown in a yellow background. The main type of telomeric repeats is shown in red and neighboring repeat units are distinguished by using regular and bold fonts alternately. All other types of repeat units are shown in black. (D) Local synteny of the genomic region contain *ter1* and its flanking protein-coding genes. 5S rRNA genes and tRNA genes are labeled as red and green arrows respectively. Gene names shown are the names of the genes in *S. pombe* except for three species-specific genes (*SPOG_05030*, *SOMG_00285*, and *SOCG_06090*) and two *wtf* genes (*Sosmo wtf2* and *Socto wtf74*). The start and end positions of the *ter1* genes correspond to the most frequent start and end positions of mature *ter1* transcripts shown in D. (D) Alignment of DNA sequences of the *ter1* gene and its flanking conserved sequences among *S. cryophilus*, *S. octosporus*, and *S. osmophilus*. Nucleotides identical to the consensus are shaded in gray. The templating region is indicated by a blue box. The most frequent start and end positions of mature *ter1* transcripts in *S. cryophilus* and *S. octosporus* reported in a previous study (Kannan *et al*. 2015) are highlighted in green letters. The introns (in a pink box) in the *ter1* genes of *S. cryophilus* and *S. octosporus* are as reported in a previous study (Kannan *et al*. 2015). The intron in the *ter1* gene of *S. osmophilus* are predicted based on sequence homology.

**Figure S9.**
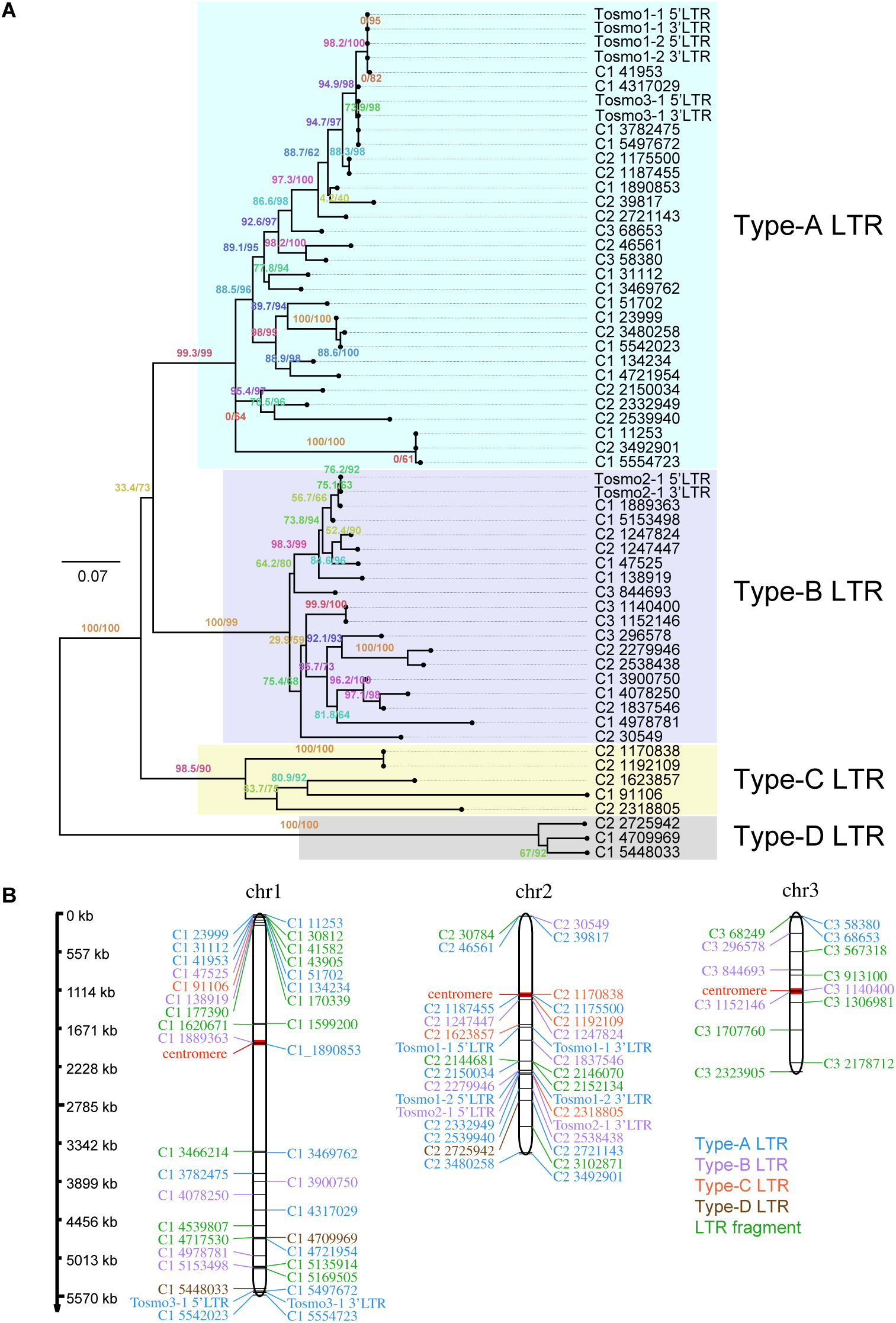
Phylogenetic relationship and chromosomal distribution of LTRs in *S. osmophilus*. (A) Maximum likelihood tree of LTRs found in the *S. osmophilus* genome. The tree was rooted by midpoint rooting. Four major branches are denoted A, B, C, and D. (B) Distribution of LTRs in the *S. osmophilus* genome.

**Figure S10.**
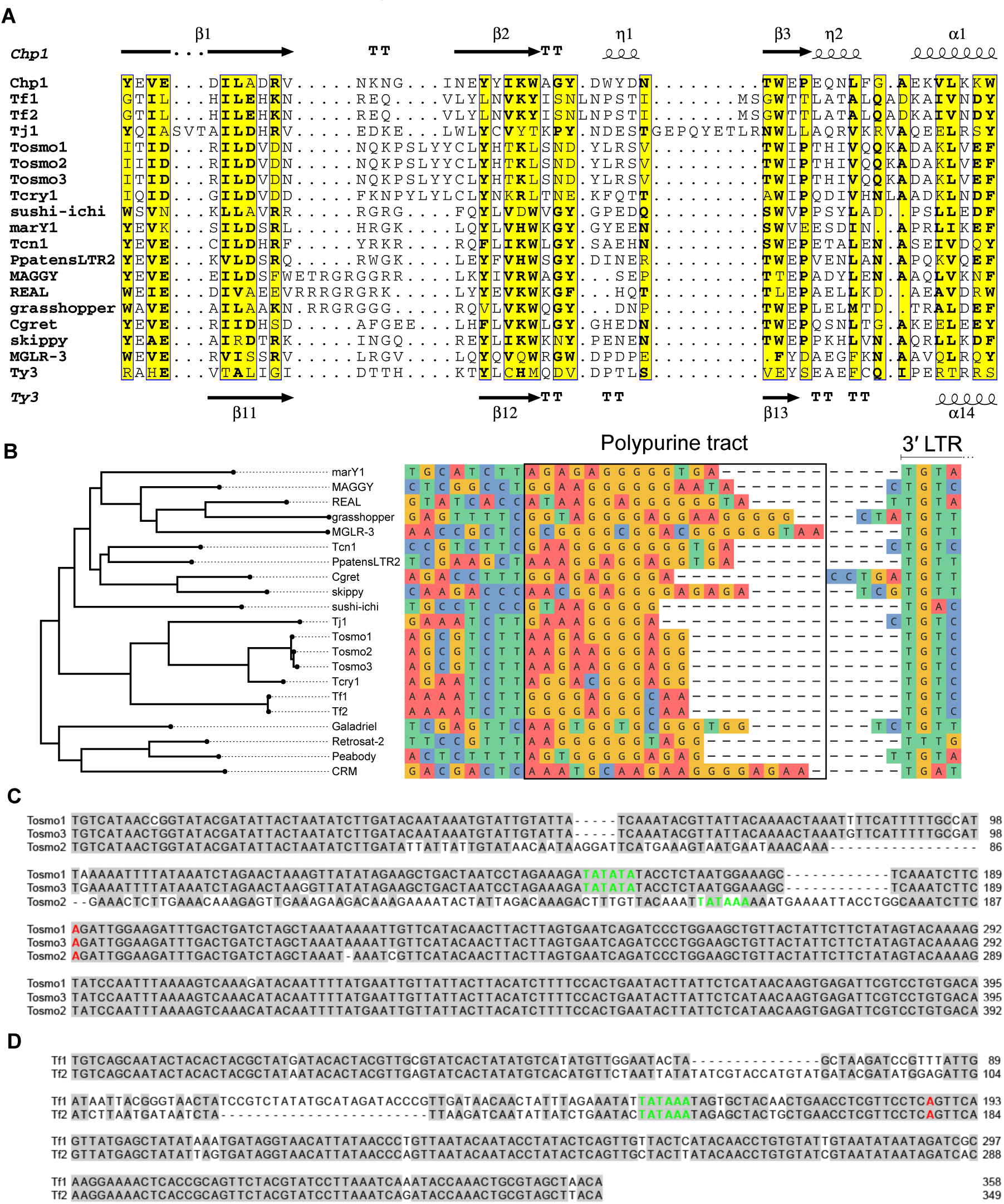
Sequence analyses of chromodomains, PPTs, and LTRs. (A) Alignment of amino acid sequences of chromodomains. Secondary structures of the chromodomains of *S. pombe* Chp1 (PDB: 3G7L) and Ty3 integrase (PDB: 7Q5B) are shown at top and bottom, respectively. (B) Alignment of the nucleotide sequences of and surrounding the polypurine tracts (PPTs). The phylogenetic tree shown on the left is the RT-IN tree shown in Figure 4C. (C) Alignment of the LTR sequences of Tosmo1, Tosmo2, and Tosmo3. Nucleotides identical to the consensus are shaded in gray. Predicted transcription start sites and TATA boxes are highlighted in red and green, respectively. Transcription start sites are at the beginning of the R region and correspond to the 5′ ends of the self-primers shown in Figure 4D. (D) Alignment of the LTR sequences of Tf1 and Tf2. Nucleotides identical to the consensus are shaded in gray. Transcription start sites and predicted TATA boxes are highlighted in red and green, respectively.

**Figure S11.**
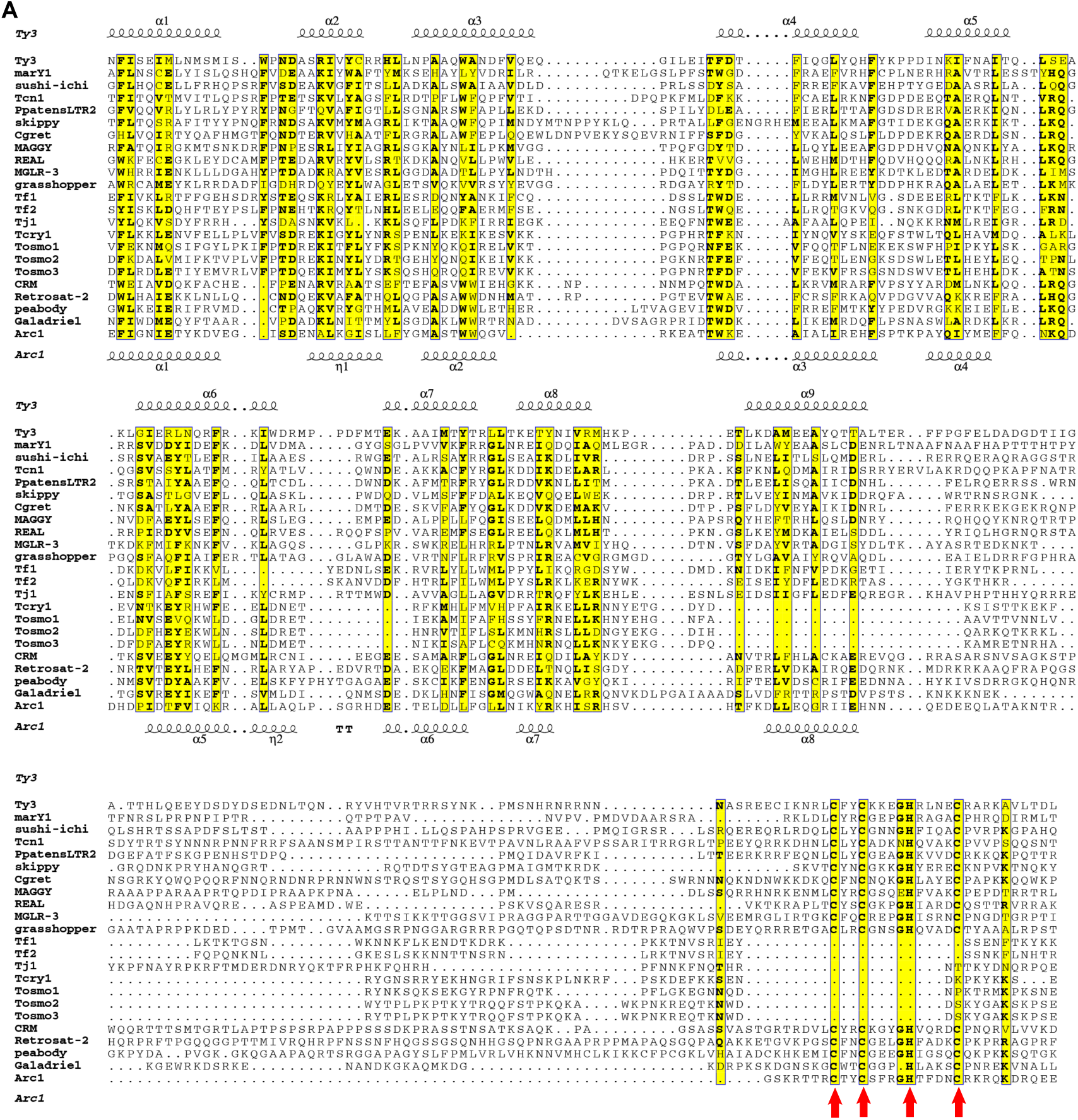
Alignment of amino acid sequences of Gag proteins. The sequences included in this alignment all begin at a stretch of sequence exhibiting similarity to helix 1 of Ty3 capsid N-terminal domain (CA-NTD). For CCHC-containing Gag proteins, the sequences end at the 10th amino acids downstream of the last cysteine of the CCHC motif; for Tosmo1, Tosmo2, Tosmo3, Tcry1, and Tj1, the sequences end at the last amino acid before the in-frame stop codon; for Tf1, the sequence ends at amino acid 245 (Teysset *et al*. 2003); for Tf2, the sequence ends at a position homologous to amino acid 245 of Tf1, and the amino acid at this position is the 4th amino acid upstream of the first β strand of the protease fold in the AlphaFold-predicted structure of the protein encoded by Tf2-3, which has a sequence identical to GenBank L10324 (https://alphafold.ebi.ac.uk/entry/P0CT36) (Varadi *et al*. 2022). Secondary structures shown at top are based on the structures of Ty3 CA-NTD (PDB 6R22) and Ty3 CA-CTD (PDB 6R23) (Dodonova *et al*. 2019). The secondary structure shown at bottom are based on the structure of *Drosophila* dArc1 (PDB 6TAS) (Erlendsson *et al*. 2020).

**Figure S12.**
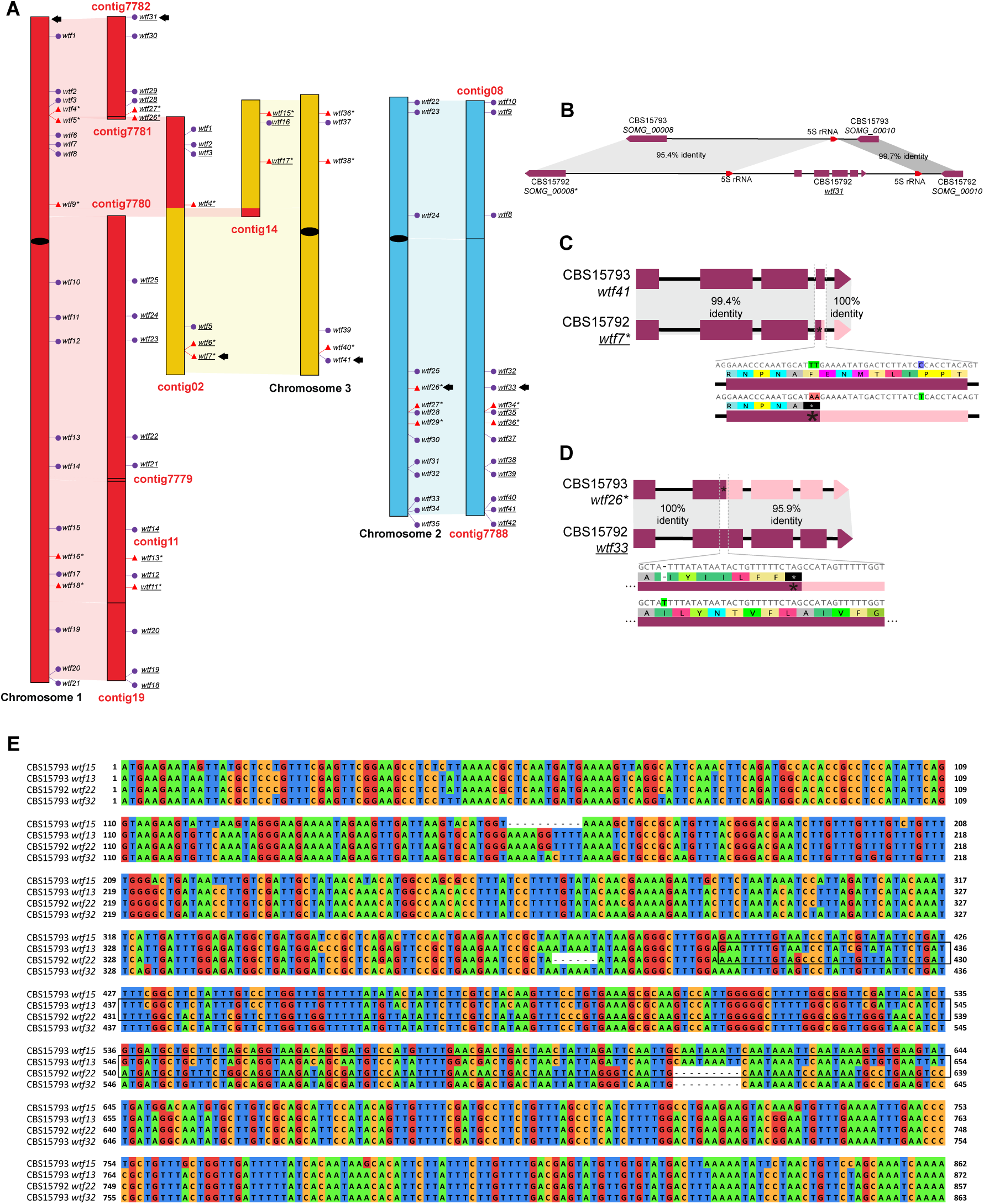
Comparisons of *wtf* genes in CBS 15792 and CBS 15793^T^. (A) The genomic distribution of *wtf* genes in CBS 15793^T^ and CBS 15792. The synteny between the three chromosomes of CBS 15793^T^ and the 10 nuclear contigs of CBS 15972 is indicated by colored connections (red, blue, and yellow for chromosomes 1, 2, and 3 of CBS 15793^T^, respectively). Centromeres in the chromosomes of CBS 15793^T^ are denoted by black ovals. The names of the 10 contigs of CBS 15972 are abbreviated such as the contig tig00007780_pilon_x4, for example, is written as contig7780. Active genes and pseudogenes are denoted by purple dots and red triangles, respectively. The 41 *wtf* genes of CBS 15793^T^ are named according to their positions in the genome. The names of the 42 *wtf* genes of CBS 15792, which are underlined, are as described (De Carvalho *et al*. 2022). The names of pseudogenes are marked with an asterisk. The presence–absence polymorphism shown in B and the differences in the status as an active gene or a pseudogene shown in C and D are highlighted using black arrows. (B) Schematic illustrating the presence–absence polymorphism of the *wtf31* gene of CBS 15792. (C) Schematic illustrating the sequence difference that results in *wtf7* of CBS 15792 being a pseudogene but its syntenic counterpart *wtf41* of CBS 15793^T^ being an active gene. (D) Schematic illustrating the sequence difference that results in *wtf33* of CBS 15792 being an active gene and its syntenic counterpart *wtf26* of CBS 15793^T^ being a pseudogene. (E) Multiple sequence alignment illustrating the sequence divergence between *wtf13* in CBS 15793^T^ and its syntenic counterpart *wtf22* in CBS 15792. Black box highlights the approximately 250-bp region where ectopic recombination may have occurred and resulted in a low nucleotide identity of 85.36% between these two syntenic genes.

**Figure S13.**
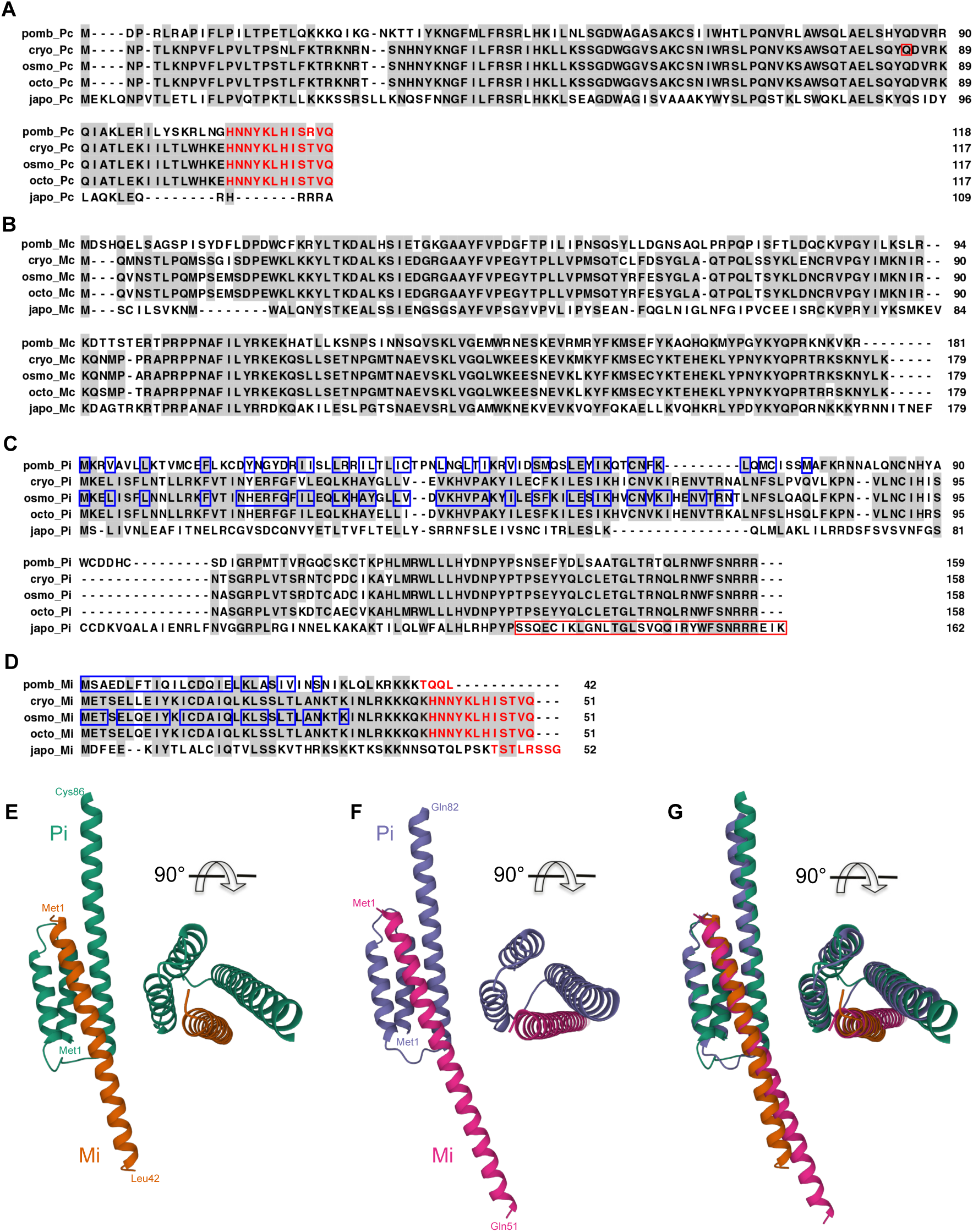
Sequences and predicted structures of mating-type cassette-encoded proteins. (A–D) Multiple sequence alignments of Pc (A), Mc (B), Pi (C), and Mi (D) of five fission yeast species. Residues identical to the consensus are shaded in gray. Residues encoded by sequences in the H2 boxes are highlighted in red letters. The Pc residue where a premature stop codon occurs in the *S. cryophilus* genome assembly reported in Tong et al. 2019 is highlighted by a red square box. The Pi residues affected by a frameshifting 1-bp deletion present in the reference *S. japonicus* genome are highlighted by a red rectangle box. Pi and Mi residues situated at the intermolecular interfaces of the AlphaFold-Multimer-predicted Pi–Mi heterodimer structures are highlighted by blue boxes. (E–F) AlphaFold-Multimer-predicted heterodimeric structures of the Pi–Mi complex of *S. pombe* (E) and the Pi–Mi complex of *S. osmophilus* (F). For clarity, only the N-terminal Mi-interacting region of Pi is shown. (H) Superposition of the structures shown in (E) and (F). The two structures were superposed using residues 1–67 of *S. pombe* Pi and residues 1–65 of *S. osmophilus* Pi.

**Figure S14.**
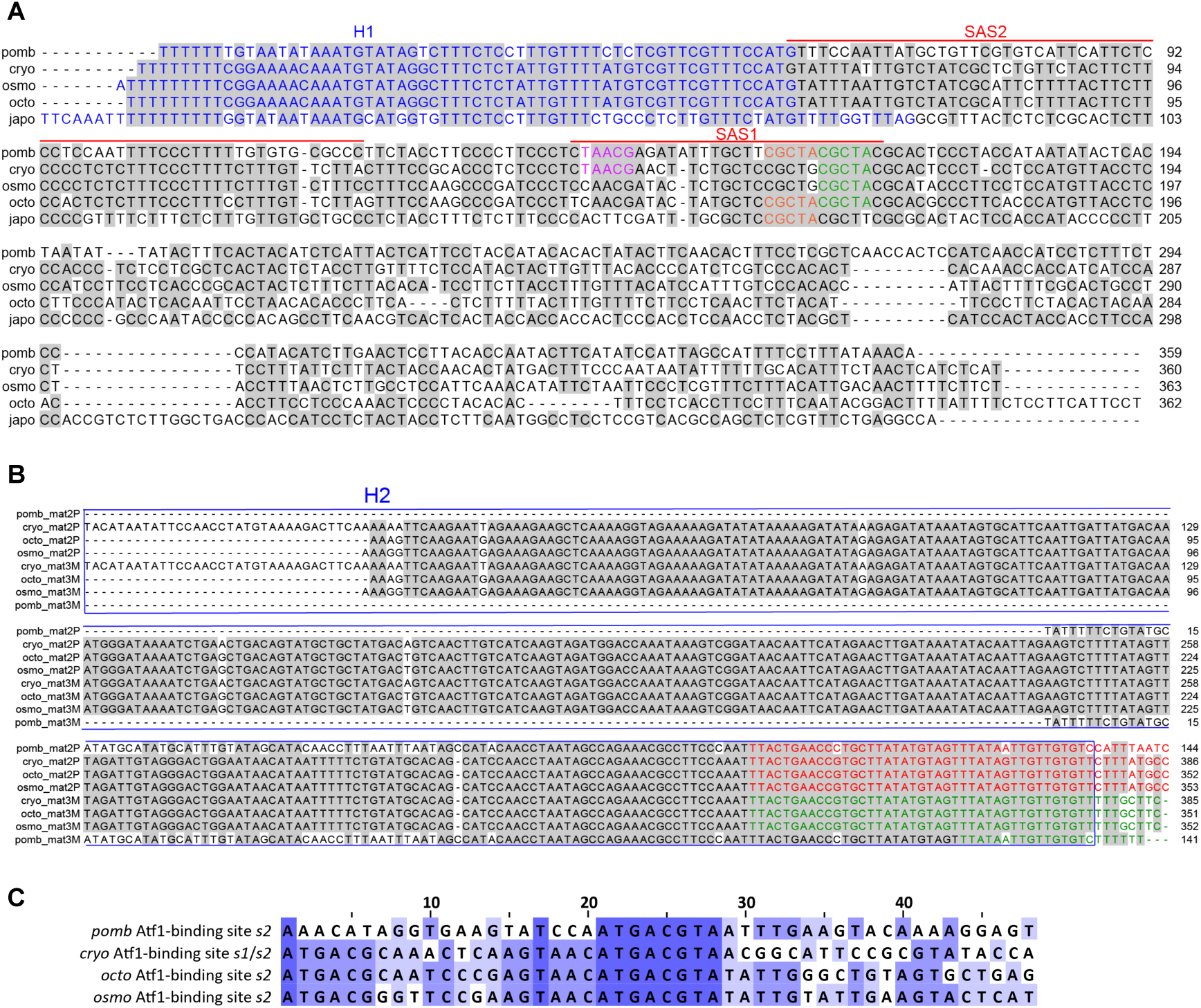
Conservation of H1, H2, the *L*-region segment containing SAS1 and SAS2, and the Atf1-binding sites. (A) Alignment of the nucleotide sequences of the 300-bp region that begins from the cassette-proximal border of H1 and extends into the *L*-region. Nucleotides identical to the consensus are shaded in gray. H1 sequences are highlighted in blue letters. SAS1 and SAS2 of *S. pombe* are highlighted by horizontal lines. Three TA(A/G)CG motifs in SAS1 of *S. pombe* are highlighted in pink, orange, and green letters. TA(A/G)CG motifs in SAS1 counterparts in *S. cryophilus*, *S. osmophilus*, *S. octosporus*, and *S. japonicus* are highlighted similarly. (B) Alignment of the nucleotide sequences of H2 boxes of *S. pombe*, *S. octosporus*, *S. cryophilus*, and *S. osmophilus*. Nucleotides identical to the consensus are shaded in gray. Coding sequences for Pc and Mi are highlighted in red letters and green letters, respectively. (C) The 48-bp sequences centered on the most cassette-proximal 8-bp Atf1-binding sites (ATGACGTA) in the donor regions of *S. pombe*, *S. cryophilus*, *S. octosporus*, and *S. osmophilus*. The *s1* and *s2* sites of *S. cryophilus* are located within the H3 boxes and therefore share the same flanking sequences. The *s2* site of *S. octosporus* is 33 bp away from the nearby H3 box and the *s2* site of *S. osmophilus* is 30 bp away from the nearby H3 box. Sequence shading is based on the level of sequence identity.

**Figure S15.**
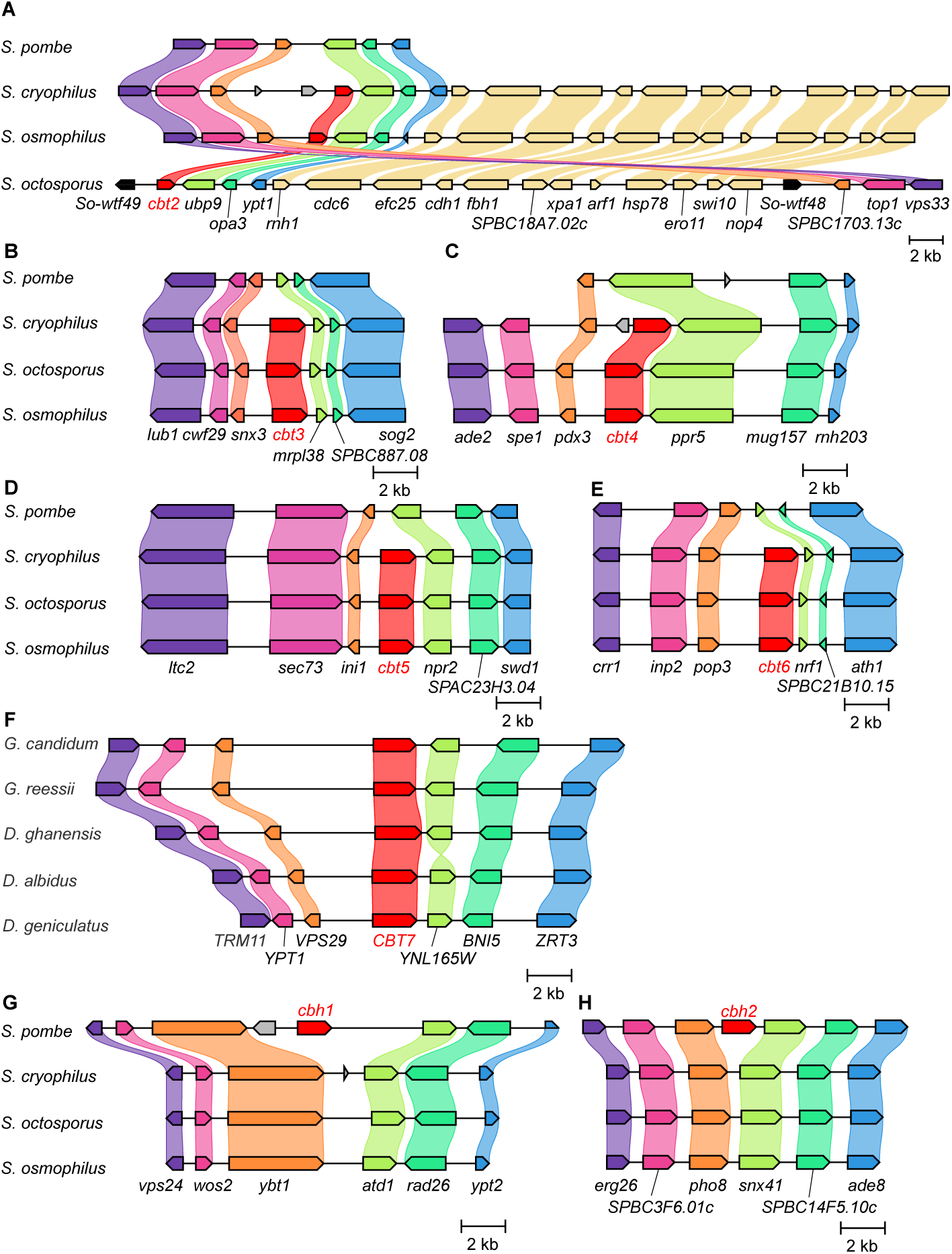
Diagrams showing the local synteny at genomic regions containing genes encoding Cbp1 family proteins, including Cbt2 (A), Cbt3 (B), Cbt4 (C), Cbt5 (D), Cbt6 (E), Cbt7 (F), Cbh1 (G), and Cbh2 (H). In (F), except for *CBT7*, gene names shown are the names of the homologous genes in *Saccharomyces cerevisiae*. In the other panels, except for *cbt1*–*cbt6* and two *wtf* genes in *S. octopous* (*wtf48*/*SOCG_00278* and *wtf49*/*SOCG_00295*), gene names shown are the names of the genes in *S. pombe*.

**Figure S16.**
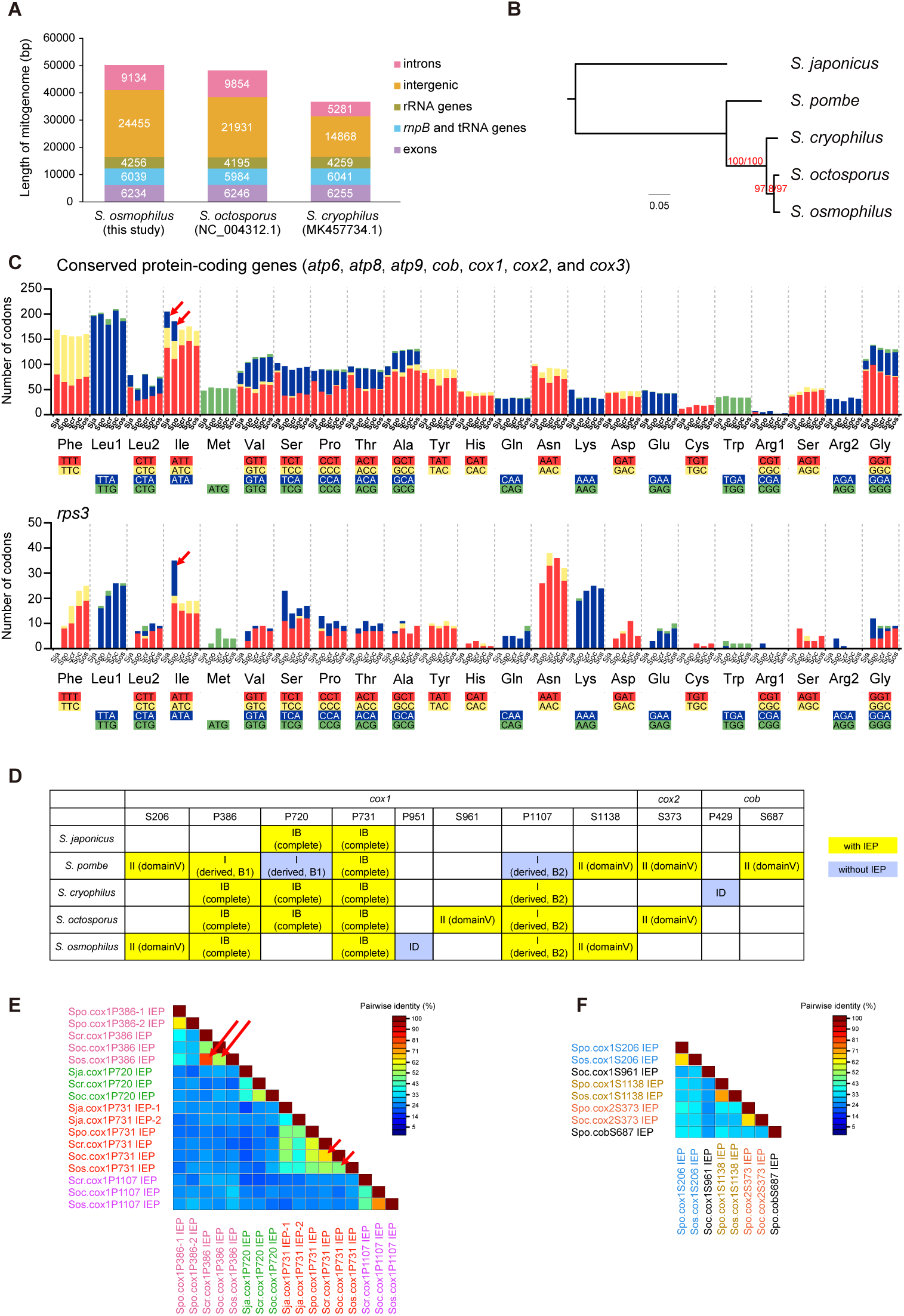
Comparative analysis of the mitogenomes of five fission yeast species. (A) Length differences between the mitogenomes of *S. octosporus*, *S. osmophilus*, and *S. cryophilus* are mainly due to variations in the lengths of introns and intergenic regions. (B) Maximum likelihood tree constructed using the amino acid sequences of the proteins encoded by the seven conserved protein-coding genes (*atp6*, *atp8*, *atp9*, *cob*, *cox1*, *cox2*, and *cox3*) in the fission yeast mitogenomes. The tree was rooted using *S. japonicus* as outgroup. Branch labels are the SH-aLRT support value (%) and the UFBoot support value (%) calculated by IQ-TREE. (C) Codon usage in the mitogenomes of five fission yeast species. Red arrows highlight the presence of ATA codons in *S. japonicus* and *S. pombe*. Sja, *S. japonicus*; Spo, *S. pombes*; Scr, *S. cryophilus*; Soc, *S. octosporus*; Sos, *S. osmophilus*. (D) Mitochondrial introns of five fission yeast species. The *S. pombe* introns listed include all introns known to exist in natural isolates of *S. pombe* (Tao *et al*. 2019). The introns of the other four species are introns present in the reference strains. RNAweasel-annotated intron type (group I or group II) and subgroup classification for group I introns are shown. Yellow and blue backgrounds indicate the presence and absence of intron-encoded proteins (IEPs), respectively. *S. pombe* has two different cox1P386 introns (cox1P386-1 and cox1P386-2). They belong to the same subgroup and both encode IEPs. (E) A color matrix showing the pair-wise amino acid identities between proteins encoded by group I introns. This color matrix was generated using Sequence Demarcation Tool Version 1.2 (SDTv1.2) (Muhire *et al*. 2014). The two long arrows highlight that the identity between Sos.cox1P386 IEP and Scr.cox1P386 IEP was higher than the identity between Sos.cox1P386 IEP and Soc.cox1P386 IEP. The two short arrows highlight that the identity between Soc.cox1P731 IEP and Scr.cox1P731 IEP was higher than the identity between Sos.cox1P731 IEP and Soc.cox1P731 IEP. (F) A color matrix showing the pair-wise amino acid identities between proteins encoded by group II introns. This color matrix was generated using Sequence Demarcation Tool Version 1.2 (SDTv1.2) (Muhire *et al*. 2014).

**Figure S17.**
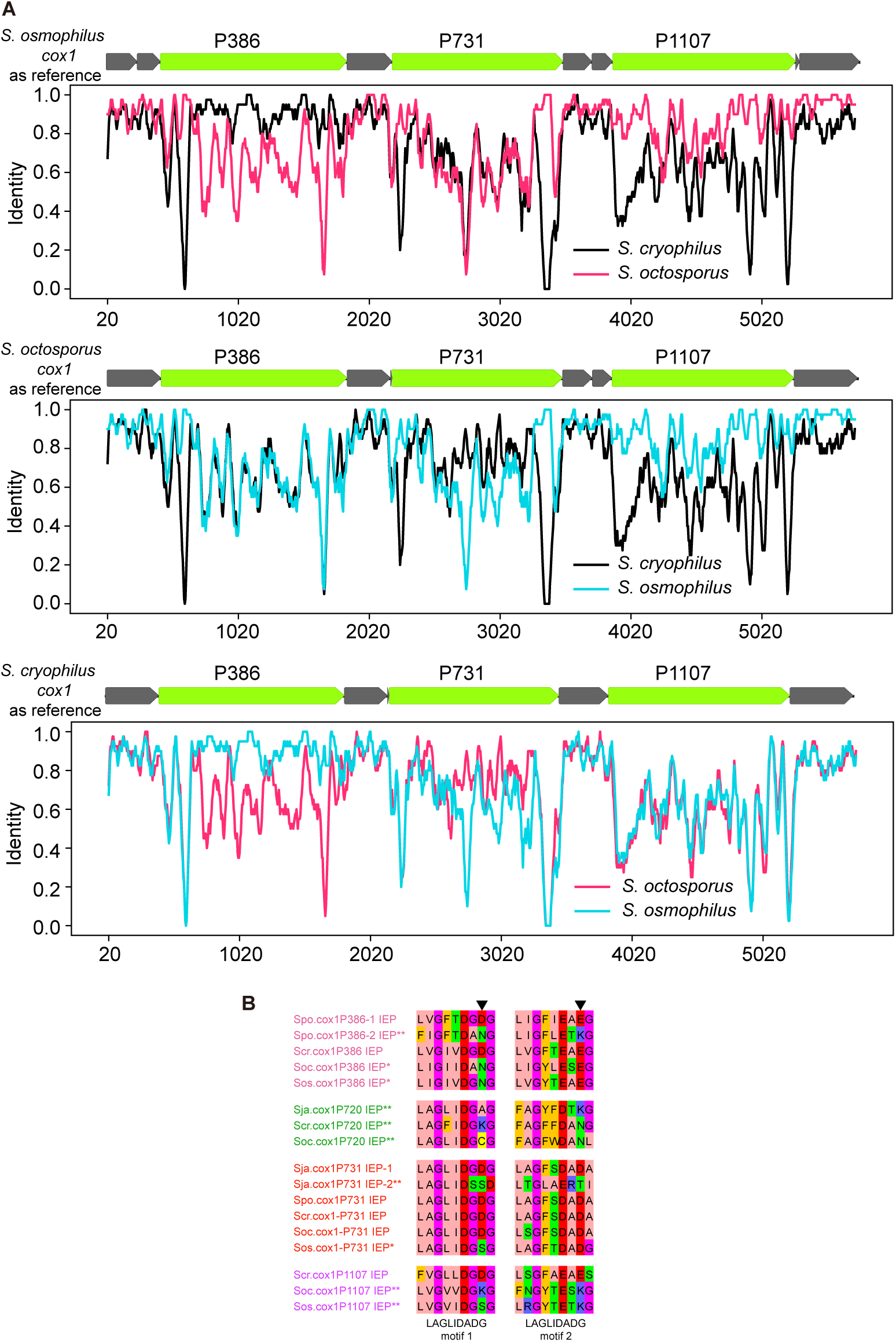
Comparative analysis of the mitogenomes of five fission yeast species. (A) Sliding-window analysis using SimPlot++ showed that the sequences that may have undergone introgression or horizontal transfer were limited within the cox1P386 intron and the cox1P731 intron. Introns that are not present in all three mitogenomes were removed before performing sequence alignment and SimPlot++ analysis. (B) The two LAGLIDADG motifs in the proteins encoded by group I introns. Black arrowheads point to the two catalytic residues, which must both be acidic residues to support DNA cleavage activities. Single asterisks indicate that the catalytic residue in one of the two LAGLIDADG motifs is not an acidic residue. Double asterisks indicate that the catalytic residue in neither LAGLIDADG motif is an acidic residue. Sja.cox1P731 intron encodes two IEPs (IEP-1 and IEP-2) and only one of them is degenerated.

## References

Ács-Szabó L., L. A. Papp, Z. Antunovics, M. Sipiczki, and I. Miklós, 2018 Assembly of Schizosaccharomyces cryophilus chromosomes and their comparative genomic analyses revealed principles of genome evolution of the haploid fission yeasts. Sci Rep 8: 14629. https://doi.org/10.1038/s41598-018-32525-9

Aoki K., K. Furuya, and H. Niki, 2017 Schizosaccharomyces japonicus: A Distinct Dimorphic Yeast among the Fission Yeasts. Cold Spring Harb Protoc 2017: pdb.top082651. https://doi.org/10.1101/pdb.top082651

Arcangioli B., and A. J. Klar, 1991 A novel switch-activating site (SAS1) and its cognate binding factor (SAP1) required for efficient mat1 switching in Schizosaccharomyces pombe. EMBO J 10: 3025–3032. https://doi.org/10.1002/j.1460-2075.1991.tb07853.x

Ard R., P. Tong, and R. C. Allshire, 2014 Long non-coding RNA-mediated transcriptional interference of a permease gene confers drug tolerance in fission yeast. Nat Commun 5: 5576. https://doi.org/10.1038/ncomms6576

Atwood-Moore A., K. Ejebe, and H. L. Levin, 2005 Specific recognition and cleavage of the plus-strand primer by reverse transcriptase. J Virol 79: 14863–14875. https://doi.org/10.1128/JVI.79.23.14863-14875.2005

Bailey T. L., M. Boden, F. A. Buske, M. Frith, C. E. Grant, et al., 2009 MEME SUITE: tools for motif discovery and searching. Nucleic Acids Res 37: W202–208. https://doi.org/10.1093/nar/gkp335

Barsoum E., P. Martinez, and S. U. Aström, 2010 Alpha3, a transposable element that promotes host sexual reproduction. Genes Dev 24: 33–44. https://doi.org/10.1101/gad.557310

Baudry C., S. Malinsky, M. Restituito, A. Kapusta, S. Rosa, et al., 2009 PiggyMac, a domesticated piggyBac transposase involved in programmed genome rearrangements in the ciliate Paramecium tetraurelia. Genes & Development 23: 2478. https://doi.org/10.1101/gad.547309

Benachenhou F., G. O. Sperber, E. Bongcam-Rudloff, G. Andersson, J. D. Boeke, et al., 2013 Conserved structure and inferred evolutionary history of long terminal repeats (LTRs). Mob DNA 4: 5. https://doi.org/10.1186/1759-8753-4-5

Bilanchone V. W., J. A. Claypool, P. T. Kinsey, and S. B. Sandmeyer, 1993 Positive and negative regulatory elements control expression of the yeast retrotransposon Ty3. Genetics 134: 685–700. https://doi.org/10.1093/genetics/134.3.685

Bonetti B., L. Fu, J. Moon, and D. M. Bedwell, 1995 The efficiency of translation termination is determined by a synergistic interplay between upstream and downstream sequences in Saccharomyces cerevisiae. J Mol Biol 251: 334–345. https://doi.org/10.1006/jmbi.1995.0438

Bowen N. J., I. K. Jordan, J. A. Epstein, V. Wood, and H. L. Levin, 2003 Retrotransposons and their recognition of pol II promoters: a comprehensive survey of the transposable elements from the complete genome sequence of Schizosaccharomyces pombe. Genome Res 13: 1984–1997. https://doi.org/10.1101/gr.1191603

Box J. A., J. T. Bunch, D. C. Zappulla, E. F. Glynn, and P. Baumann, 2008 A flexible template boundary element in the RNA subunit of fission yeast telomerase. J Biol Chem 283: 24224–24233. https://doi.org/10.1074/jbc.M802043200

Brysch-Herzberg M., A. Tobias, M. Seidel, R. Wittmann, E. Wohlmann, et al., 2019 Schizosaccharomyces osmophilus sp. nov., an osmophilic fission yeast occurring in bee bread of different solitary bee species. FEMS Yeast Res 19: foz038. https://doi.org/10.1093/femsyr/foz038

Brysch-Herzberg M., G.-S. Jia, M. Seidel, I. Assali, and L.-L. Du, 2022 Insights into the ecology of Schizosaccharomyces species in natural and artificial habitats. Antonie Van Leeuwenhoek 115: 661–695. https://doi.org/10.1007/s10482-022-01720-0

Bullerwell C. E., J. Leigh, L. Forget, and B. F. Lang, 2003 A comparison of three fission yeast mitochondrial genomes. Nucleic Acids Res 31: 759–768. https://doi.org/10.1093/nar/gkg134

Butler M., T. Goodwin, M. Simpson, M. Singh, and R. Poulter, 2001 Vertebrate LTR retrotransposons of the Tf1/sushi group. J Mol Evol 52: 260–274. https://doi.org/10.1007/s002390010154

Cam H. P., K. Noma, H. Ebina, H. L. Levin, and S. I. S. Grewal, 2008 Host genome surveillance for retrotransposons by transposon-derived proteins. Nature 451: 431–436. https://doi.org/10.1038/nature06499

Campbell M. S., C. Holt, B. Moore, and M. Yandell, 2014 Genome Annotation and Curation Using MAKER and MAKER-P. Curr Protoc Bioinformatics 48: 4.11.1–39. https://doi.org/10.1002/0471250953.bi0411s48

Cantalapiedra C. P., A. Hernández-Plaza, I. Letunic, P. Bork, and J. Huerta-Cepas, 2021 eggNOG-mapper v2: Functional Annotation, Orthology Assignments, and Domain Prediction at the Metagenomic Scale. Mol Biol Evol 38: 5825–5829. https://doi.org/10.1093/molbev/msab293

Capella-Gutiérrez S., J. M. Silla-Martínez, and T. Gabaldón, 2009 trimAl: a tool for automated alignment trimming in large-scale phylogenetic analyses. Bioinformatics 25: 1972–1973. https://doi.org/10.1093/bioinformatics/btp348

Casola C., D. Hucks, and C. Feschotte, 2008 Convergent domestication of pogo-like transposases into centromere-binding proteins in fission yeast and mammals. Mol Biol Evol 25: 29–41. https://doi.org/10.1093/molbev/msm221

Chan P. P., B. Y. Lin, A. J. Mak, and T. M. Lowe, 2021 tRNAscan-SE 2.0: improved detection and functional classification of transfer RNA genes. Nucleic Acids Res 49: 9077–9096. https://doi.org/10.1093/nar/gkab688

Chapman E., F. Taglini, and E. H. Bayne, 2022 Separable roles for RNAi in regulation of transposable elements and viability in the fission yeast Schizosaccharomyces japonicus. PLoS Genet 18: e1010100. https://doi.org/10.1371/journal.pgen.1010100

Chen S., Y. Zhou, Y. Chen, and J. Gu, 2018 fastp: an ultra-fast all-in-one FASTQ preprocessor. Bioinformatics 34: i884–i890. https://doi.org/10.1093/bioinformatics/bty560

Cheng C.-Y., A. Vogt, K. Mochizuki, and M.-C. Yao, 2010 A domesticated piggyBac transposase plays key roles in heterochromatin dynamics and DNA cleavage during programmed DNA deletion in Tetrahymena thermophila. Mol Biol Cell 21: 1753–1762. https://doi.org/10.1091/mbc.e09-12-1079

Cheng H., G. T. Concepcion, X. Feng, H. Zhang, and H. Li, 2021 Haplotype-resolved de novo assembly using phased assembly graphs with hifiasm. Nat Methods 18: 170–175. https://doi.org/10.1038/s41592-020-01056-5

Chiron S., M. Gaisne, E. Guillou, P. Belenguer, G. D. Clark-Walker, et al., 2007 Studying mitochondria in an attractive model: Schizosaccharomyces pombe. Methods Mol Biol 372: 91–105. https://doi.org/10.1007/978-1-59745-365-3_7

Cottee M. A., S. L. Beckwith, S. C. Letham, S. J. Kim, G. R. Young, et al., 2021 Structure of a Ty1 restriction factor reveals the molecular basis of transposition copy number control. Nat Commun 12: 5590. https://doi.org/10.1038/s41467-021-25849-0

Couch B. C., and L. M. Kohn, 2002 A multilocus gene genealogy concordant with host preference indicates segregation of a new species, Magnaporthe oryzae, from M. grisea. Mycologia 94: 683–693. https://doi.org/10.1080/15572536.2003.11833196

Covey S. N., 1986 Amino acid sequence homology in gag region of reverse transcribing elements and the coat protein gene of cauliflower mosaic virus. Nucleic Acids Res 14: 623–633. https://doi.org/10.1093/nar/14.2.623

Creevey C., and N. Weeks, 2021 ChrisCreevey/catsequences: Version 1.3

Cridge A. G., C. Crowe-McAuliffe, S. F. Mathew, and W. P. Tate, 2018 Eukaryotic translational termination efficiency is influenced by the 3’ nucleotides within the ribosomal mRNA channel. Nucleic Acids Res 46: 1927–1944. https://doi.org/10.1093/nar/gkx1315

De Carvalho M., G.-S. Jia, A. Nidamangala Srinivasa, R. B. Billmyre, Y.-H. Xu, et al., 2022 The wtf meiotic driver gene family has unexpectedly persisted for over 100 million years. Elife 11: e81149. https://doi.org/10.7554/eLife.81149

Dobin A., C. A. Davis, F. Schlesinger, J. Drenkow, C. Zaleski, et al., 2013 STAR: ultrafast universal RNA-seq aligner. Bioinformatics 29: 15–21. https://doi.org/10.1093/bioinformatics/bts635

Dobinson K. F., R. E. Harris, and J. E. Hamer, 1993 Grasshopper, a long terminal repeat (LTR) retroelement in the phytopathogenic fungus Magnaporthe grisea. Mol Plant Microbe Interact 6: 114–126. https://doi.org/10.1094/mpmi-6-114

Dodonova S. O., S. Prinz, V. Bilanchone, S. Sandmeyer, and J. A. G. Briggs, 2019 Structure of the Ty3/Gypsy retrotransposon capsid and the evolution of retroviruses. Proc Natl Acad Sci U S A 116: 10048–10057. https://doi.org/10.1073/pnas.1900931116

Drost H.-G., and D. H. Sanchez, 2019 Becoming a Selfish Clan: Recombination Associated to Reverse-Transcription in LTR Retrotransposons. Genome Biol Evol 11: 3382–3392. https://doi.org/10.1093/gbe/evz255

Dupeyron M., T. Baril, C. Bass, and A. Hayward, 2020 Phylogenetic analysis of the Tc1/mariner superfamily reveals the unexplored diversity of pogo-like elements. Mob DNA 11: 21. https://doi.org/10.1186/s13100-020-00212-0

Erlendsson S., D. R. Morado, H. B. Cullen, C. Feschotte, J. D. Shepherd, et al., 2020 Structures of virus-like capsids formed by the Drosophila neuronal Arc proteins. Nat Neurosci 23: 172–175. https://doi.org/10.1038/s41593-019-0569-y

Esnault C., and H. L. Levin, 2015 The Long Terminal Repeat Retrotransposons Tf1 and Tf2 of Schizosaccharomyces pombe. Microbiol Spectr 3. https://doi.org/10.1128/microbiolspec.MDNA3-0040-2014

Evans R., M. O’Neill, A. Pritzel, N. Antropova, A. Senior, et al., 2022 Protein complex prediction with AlphaFold-Multimer. BioRxiv 2021.10. 04.463034.

Fajkus P., A. Kilar, A. D. L. Nelson, M. Holá, V. Peška, et al., 2021 Evolution of plant telomerase RNAs: farther to the past, deeper to the roots. Nucleic Acids Res 49: 7680–7694. https://doi.org/10.1093/nar/gkab545

Fantes P. A., and C. S. Hoffman, 2016 A Brief History of Schizosaccharomyces pombe Research: A Perspective Over the Past 70 Years. Genetics 203: 621–629. https://doi.org/10.1534/genetics.116.189407

Férandon C., S. Moukha, P. Callac, J.-P. Benedetto, M. Castroviejo, et al., 2010 The Agaricus bisporus cox1 gene: the longest mitochondrial gene and the largest reservoir of mitochondrial group i introns. PLoS One 5: e14048. https://doi.org/10.1371/journal.pone.0014048

Firoozan M., C. M. Grant, J. A. Duarte, and M. F. Tuite, 1991 Quantitation of readthrough of termination codons in yeast using a novel gene fusion assay. Yeast 7: 173–183. https://doi.org/10.1002/yea.320070211

Forbes E. M., S. R. Nieduszynska, F. K. Brunton, J. Gibson, L. A. Glover, et al., 2007 Control of gag-pol gene expression in the Candida albicans retrotransposon Tca2. BMC Mol Biol 8: 94. https://doi.org/10.1186/1471-2199-8-94

Formenti G., A. Rhie, J. Balacco, B. Haase, J. Mountcastle, et al., 2021 Complete vertebrate mitogenomes reveal widespread repeats and gene duplications. Genome Biol 22: 120. https://doi.org/10.1186/s13059-021-02336-9

Forsburg S. L., and N. Rhind, 2006 Basic methods for fission yeast. Yeast 23: 173–183. https://doi.org/10.1002/yea.1347

Gao X., E. R. Havecker, P. V. Baranov, J. F. Atkins, and D. F. Voytas, 2003 Translational recoding signals between gag and pol in diverse LTR retrotransposons. RNA 9: 1422–1430. https://doi.org/10.1261/rna.5105503

Gao B., Y. Wang, M. Diaby, W. Zong, D. Shen, et al., 2020 Evolution of pogo, a separate superfamily of IS630-Tc1-mariner transposons, revealing recurrent domestication events in vertebrates. Mob DNA 11: 25. https://doi.org/10.1186/s13100-020-00220-0

Ghazvini M., V. Ribes, and B. Arcangioli, 1995 The essential DNA-binding protein sap1 of Schizosaccharomyces pombe contains two independent oligomerization interfaces that dictate the relative orientation of the DNA-binding domain. Mol Cell Biol 15: 4939–4946. https://doi.org/10.1128/MCB.15.9.4939

Gilchrist C. L. M., and Y.-H. Chooi, 2021 Clinker & clustermap.js: Automatic generation of gene cluster comparison figures. Bioinformatics btab007. https://doi.org/10.1093/bioinformatics/btab007

Gómez Luciano L. B., I. J. Tsai, I. Chuma, Y. Tosa, Y.-H. Chen, et al., 2019 Blast Fungal Genomes Show Frequent Chromosomal Changes, Gene Gains and Losses, and Effector Gene Turnover. Mol Biol Evol 36: 1148– 1161. https://doi.org/10.1093/molbev/msz045

Goodwin T. J., and R. T. Poulter, 2001 The diversity of retrotransposons in the yeast Cryptococcus neoformans. Yeast 18: 865–880. https://doi.org/10.1002/yea.733

Guindon S., J.-F. Dufayard, V. Lefort, M. Anisimova, W. Hordijk, et al., 2010 New algorithms and methods to estimate maximum-likelihood phylogenies: assessing the performance of PhyML 3.0. Syst Biol 59: 307–321. https://doi.org/10.1093/sysbio/syq010

Guo Y., P. K. Singh, and H. L. Levin, 2015 A long terminal repeat retrotransposon of Schizosaccharomyces japonicus integrates upstream of RNA pol III transcribed genes. Mob DNA 6: 19. https://doi.org/10.1186/s13100-015-0048-2

Haas B. J., S. L. Salzberg, W. Zhu, M. Pertea, J. E. Allen, et al., 2008 Automated eukaryotic gene structure annotation using EVidenceModeler and the Program to Assemble Spliced Alignments. Genome Biol 9: R7. https://doi.org/10.1186/gb-2008-9-1-r7

Hackl T., S. Duponchel, K. Barenhoff, A. Weinmann, and M. G. Fischer, 2021 Virophages and retrotransposons colonize the genomes of a heterotrophic flagellate. Elife 10: e72674. https://doi.org/10.7554/eLife.72674

Hall M. B., 2022 Rasusa: Randomly subsample sequencing reads to a specified coverage. Journal of Open Source Software 7: 3941. https://doi.org/10.21105/joss.03941

Hanson S. J., and K. H. Wolfe, 2017 An Evolutionary Perspective on Yeast Mating-Type Switching. Genetics 206: 9–32. https://doi.org/10.1534/genetics.117.202036

Harris M. A., K. M. Rutherford, J. Hayles, A. Lock, J. Bähler, et al., 2022 Fission stories: using PomBase to understand Schizosaccharomyces pombe biology. Genetics 220: iyab222. https://doi.org/10.1093/genetics/iyab222

Hayles J., V. Wood, L. Jeffery, K.-L. Hoe, D.-U. Kim, et al., 2013 A genome-wide resource of cell cycle and cell shape genes of fission yeast. Open Biol 3: 130053. https://doi.org/10.1098/rsob.130053

Hayles J., and P. Nurse, 2018 Introduction to Fission Yeast as a Model System. Cold Spring Harb Protoc 2018. https://doi.org/10.1101/pdb.top079749

Helston R. M., J. A. Box, W. Tang, and P. Baumann, 2010 Schizosaccharomyces cryophilus sp. nov., a new species of fission yeast. FEMS Yeast Res 10: 779–786. https://doi.org/10.1111/j.1567-1364.2010.00657.x

Hoffman C. S., V. Wood, and P. A. Fantes, 2015 An Ancient Yeast for Young Geneticists: A Primer on the Schizosaccharomyces pombe Model System. Genetics 201: 403–423. https://doi.org/10.1534/genetics.115.181503

Hu W., Z.-D. Jiang, F. Suo, J.-X. Zheng, W.-Z. He, et al., 2017 A large gene family in fission yeast encodes spore killers that subvert Mendel’s law. Elife 6: e26057. https://doi.org/10.7554/eLife.26057

Huang S., X. Tao, S. Yuan, Y. Zhang, P. Li, et al., 2016 Discovery of an Active RAG Transposon Illuminates the Origins of V(D)J Recombination. Cell 166: 102–114. https://doi.org/10.1016/j.cell.2016.05.032

Ilyinskii P. O., and R. C. Desrosiers, 1998 Identification of a sequence element immediately upstream of the polypurine tract that is essential for replication of simian immunodeficiency virus. EMBO J 17: 3766–3774. https://doi.org/10.1093/emboj/17.13.3766

Irelan J. T., G. I. Gutkin, and L. Clarke, 2001 Functional redundancies, distinct localizations and interactions among three fission yeast homologs of centromere protein-B. Genetics 157: 1191–1203. https://doi.org/10.1093/genetics/157.3.1191

Jin J.-J., W.-B. Yu, J.-B. Yang, Y. Song, C. W. dePamphilis, et al., 2020 GetOrganelle: a fast and versatile toolkit for accurate de novo assembly of organelle genomes. Genome Biol 21: 241. https://doi.org/10.1186/s13059-020-02154-5

Jungreis I., M. F. Lin, R. Spokony, C. S. Chan, N. Negre, et al., 2011 Evidence of abundant stop codon readthrough in Drosophila and other metazoa. Genome Res 21: 2096–2113. https://doi.org/10.1101/gr.119974.110

Kalyaanamoorthy S., B. Q. Minh, T. K. F. Wong, A. von Haeseler, and L. S. Jermiin, 2017 ModelFinder: fast model selection for accurate phylogenetic estimates. Nat Methods 14: 587–589. https://doi.org/10.1038/nmeth.4285

Kamatani T., and T. Yamamoto, 2007 Comparison of codon usage and tRNAs in mitochondrial genomes of Candida species. Biosystems 90: 362–370. https://doi.org/10.1016/j.biosystems.2006.09.039

Kannan R., R. M. Helston, R. O. Dannebaum, and P. Baumann, 2015 Diverse mechanisms for spliceosome-mediated 3’ end processing of telomerase RNA. Nat Commun 6: 6104. https://doi.org/10.1038/ncomms7104

Katoh K., and D. M. Standley, 2013 MAFFT multiple sequence alignment software version 7: improvements in performance and usability. Mol Biol Evol 30: 772–780. https://doi.org/10.1093/molbev/mst010

Kelly M., J. Burke, M. Smith, A. Klar, and D. Beach, 1988 Four mating-type genes control sexual differentiation in the fission yeast. EMBO J 7: 1537–1547. https://doi.org/10.1002/j.1460-2075.1988.tb02973.x

Kelly F. D., and H. L. Levin, 2005 The evolution of transposons in Schizosaccharomyces pombe. Cytogenet Genome Res 110: 566–574. https://doi.org/10.1159/000084990

Kikuchi J., J. Iwahara, T. Kigawa, Y. Murakami, T. Okazaki, et al., 2002 Solution structure determination of the two DNA-binding domains in the Schizosaccharomyces pombe Abp1 protein by a combination of dipolar coupling and diffusion anisotropy restraints. J Biomol NMR 22: 333–347. https://doi.org/10.1023/a:1014977808170

Kim D.-U., J. Hayles, D. Kim, V. Wood, H.-O. Park, et al., 2010 Analysis of a genome-wide set of gene deletions in the fission yeast Schizosaccharomyces pombe. Nat Biotechnol 28: 617–623. https://doi.org/10.1038/nbt.1628

Klar A. J. S., 2013 Schizosaccharomyces japonicus yeast poised to become a favorite experimental organism for eukaryotic research. G3 (Bethesda) 3: 1869–1873. https://doi.org/10.1534/g3.113.007187

Klar A. J. S., K. Ishikawa, and S. Moore, 2014 A Unique DNA Recombination Mechanism of the Mating/Cell-type Switching of Fission Yeasts: a Review. Microbiol Spectr 2. https://doi.org/10.1128/microbiolspec.MDNA3-0003-2014

Korf I., 2004 Gene finding in novel genomes. BMC Bioinformatics 5: 59. https://doi.org/10.1186/1471-2105-5-59

Krissinel E., and K. Henrick, 2007 Inference of macromolecular assemblies from crystalline state. J Mol Biol 372: 774–797. https://doi.org/10.1016/j.jmb.2007.05.022

Krzywinski M., J. Schein, I. Birol, J. Connors, R. Gascoyne, et al., 2009 Circos: an information aesthetic for comparative genomics. Genome Res 19: 1639–1645. https://doi.org/10.1101/gr.092759.109

Kumar S., G. Stecher, M. Suleski, and S. B. Hedges, 2017 TimeTree: A Resource for Timelines, Timetrees, and Divergence Times. Mol Biol Evol 34: 1812–1819. https://doi.org/10.1093/molbev/msx116

Kumar P., and D. N. Woolfson, 2021 Socket2: A Program for Locating, Visualising, and Analysing Coiled-coil Interfaces in Protein Structures. Bioinformatics btab631. https://doi.org/10.1093/bioinformatics/btab631

Lagesen K., P. Hallin, E. A. Rødland, H.-H. Staerfeldt, T. Rognes, et al., 2007 RNAmmer: consistent and rapid annotation of ribosomal RNA genes. Nucleic Acids Res 35: 3100–3108. https://doi.org/10.1093/nar/gkm160

Lang B. F., M.-J. Laforest, and G. Burger, 2007 Mitochondrial introns: a critical view. Trends Genet 23: 119–125. https://doi.org/10.1016/j.tig.2007.01.006

Lechner M., S. Findeiss, L. Steiner, M. Marz, P. F. Stadler, et al., 2011 Proteinortho: detection of (co-)orthologs in large-scale analysis. BMC Bioinformatics 12: 124. https://doi.org/10.1186/1471-2105-12-124

Lechner M., M. Hernandez-Rosales, D. Doerr, N. Wieseke, A. Thévenin, et al., 2014 Orthology detection combining clustering and synteny for very large datasets. PLoS One 9: e105015. https://doi.org/10.1371/journal.pone.0105015

Leng P., D. H. Klatte, G. Schumann, J. D. Boeke, and T. L. Steck, 1998 Skipper, an LTR retrotransposon of Dictyostelium. Nucleic Acids Res 26: 2008–2015. https://doi.org/10.1093/nar/26.8.2008

Leupold U., 1950 Die Vererbung von Homothallie und Heterothallie bei Schizosaccharomyces pombe. Compt. Rend. Lab. Carlsberg 24: 381–480.

Levin H. L., D. C. Weaver, and J. D. Boeke, 1990 Two related families of retrotransposons from Schizosaccharomyces pombe. Mol Cell Biol 10: 6791–6798. https://doi.org/10.1128/mcb.10.12.6791-6798.1990

Levin H. L., 1995 A novel mechanism of self-primed reverse transcription defines a new family of retroelements. Mol Cell Biol 15: 3310–3317. https://doi.org/10.1128/MCB.15.6.3310

Levin H. L., 1996 An unusual mechanism of self-primed reverse transcription requires the RNase H domain of reverse transcriptase to cleave an RNA duplex. Mol Cell Biol 16: 5645–5654. https://doi.org/10.1128/MCB.16.10.5645

Levin H. L., 2007 Newly Identified Retrotransposons of the Ty3/gypsy Class in Fungi, Plants, and Vertebrates, pp. 684–701 in Mobile DNA II, John Wiley & Sons, Ltd.

Li G., and C. M. Rice, 1993 The signal for translational readthrough of a UGA codon in Sindbis virus RNA involves a single cytidine residue immediately downstream of the termination codon. J Virol 67: 5062–5067. https://doi.org/10.1128/JVI.67.8.5062-5067.1993

Li H., and R. Durbin, 2009 Fast and accurate short read alignment with Burrows-Wheeler transform. Bioinformatics 25: 1754–1760. https://doi.org/10.1093/bioinformatics/btp324

Li H., 2011 A statistical framework for SNP calling, mutation discovery, association mapping and population genetical parameter estimation from sequencing data. Bioinformatics 27: 2987–2993. https://doi.org/10.1093/bioinformatics/btr509

Li H., 2018 Minimap2: pairwise alignment for nucleotide sequences. Bioinformatics 34: 3094–3100. https://doi.org/10.1093/bioinformatics/bty191

Li J., H.-T. Wang, W.-T. Wang, X.-R. Zhang, F. Suo, et al., 2019 Systematic analysis reveals the prevalence and principles of bypassable gene essentiality. Nat Commun 10: 1002. https://doi.org/10.1038/s41467-019-08928-1

Lin J. H., and H. L. Levin, 1997a A complex structure in the mRNA of Tf1 is recognized and cleaved to generate the primer of reverse transcription. Genes Dev 11: 270–285. https://doi.org/10.1101/gad.11.2.270

Lin J. H., and H. L. Levin, 1997b Self-primed reverse transcription is a mechanism shared by several LTR-containing retrotransposons. RNA 3: 952–953.

Liu Y., J. W. Leigh, H. Brinkmann, M. T. Cushion, N. Rodriguez-Ezpeleta, et al., 2009 Phylogenomic analyses support the monophyly of Taphrinomycotina, including Schizosaccharomyces fission yeasts. Mol Biol Evol 26: 27–34. https://doi.org/10.1093/molbev/msn221

Llorens C., M. A. Fares, and A. Moya, 2008 Relationships of gag-pol diversity between Ty3/Gypsy and Retroviridae LTR retroelements and the three kings hypothesis. BMC Evol Biol 8: 276. https://doi.org/10.1186/1471-2148-8-276

Llorens C., A. Muñoz-Pomer, L. Bernad, H. Botella, and A. Moya, 2009 Network dynamics of eukaryotic LTR retroelements beyond phylogenetic trees. Biol Direct 4: 41. https://doi.org/10.1186/1745-6150-4-41

Llorens C., R. Futami, L. Covelli, L. Domínguez-Escribá, J. M. Viu, et al., 2011 The Gypsy Database (GyDB) of mobile genetic elements: release 2.0. Nucleic Acids Res 39: D70–74. https://doi.org/10.1093/nar/gkq1061

Loftus B. J., E. Fung, P. Roncaglia, D. Rowley, P. Amedeo, et al., 2005 The genome of the basidiomycetous yeast and human pathogen Cryptococcus neoformans. Science 307: 1321–1324. https://doi.org/10.1126/science.1103773

Loughran G., M.-Y. Chou, I. P. Ivanov, I. Jungreis, M. Kellis, et al., 2014 Evidence of efficient stop codon readthrough in four mammalian genes. Nucleic Acids Res 42: 8928–8938. https://doi.org/10.1093/nar/gku608

Lowe T. M., and P. P. Chan, 2016 tRNAscan-SE On-line: integrating search and context for analysis of transfer RNA genes. Nucleic Acids Res 44: W54–57. https://doi.org/10.1093/nar/gkw413

Lynch M., and J. S. Conery, 2000 The evolutionary fate and consequences of duplicate genes. Science 290: 1151– 1155. https://doi.org/10.1126/science.290.5494.1151

Majors J. E., and H. E. Varmus, 1981 Nucleotide sequences at host-proviral junctions for mouse mammary tumour virus. Nature 289: 253–258. https://doi.org/10.1038/289253a0

Marçais G., A. L. Delcher, A. M. Phillippy, R. Coston, S. L. Salzberg, et al., 2018 MUMmer4: A fast and versatile genome alignment system. PLoS Comput Biol 14: e1005944. https://doi.org/10.1371/journal.pcbi.1005944

Marín I., and C. Lloréns, 2000 Ty3/Gypsy retrotransposons: description of new Arabidopsis thaliana elements and evolutionary perspectives derived from comparative genomic data. Mol Biol Evol 17: 1040–1049. https://doi.org/10.1093/oxfordjournals.molbev.a026385

Mateo L., and J. González, 2014 Pogo-like transposases have been repeatedly domesticated into CENP-B-related proteins. Genome Biol Evol 6: 2008–2016. https://doi.org/10.1093/gbe/evu153

Matthews G. D., T. J. Goodwin, M. I. Butler, T. A. Berryman, and R. T. Poulter, 1997 pCal, a highly unusual Ty1/copia retrotransposon from the pathogenic yeast Candida albicans. J Bacteriol 179: 7118–7128. https://doi.org/10.1128/jb.179.22.7118-7128.1997

Maxwell P. H., 2020 Diverse transposable element landscapes in pathogenic and nonpathogenic yeast models: the value of a comparative perspective. Mob DNA 11: 16. https://doi.org/10.1186/s13100-020-00215-x

McKenna A., M. Hanna, E. Banks, A. Sivachenko, K. Cibulskis, et al., 2010 The Genome Analysis Toolkit: a MapReduce framework for analyzing next-generation DNA sequencing data. Genome Res 20: 1297–1303. https://doi.org/10.1101/gr.107524.110

Minh B. Q., M. A. T. Nguyen, and A. von Haeseler, 2013 Ultrafast approximation for phylogenetic bootstrap. Mol Biol Evol 30: 1188–1195. https://doi.org/10.1093/molbev/mst024

Minh B. Q., H. A. Schmidt, O. Chernomor, D. Schrempf, M. D. Woodhams, et al., 2020 IQ-TREE 2: New Models and Efficient Methods for Phylogenetic Inference in the Genomic Era. Mol Biol Evol 37: 1530–1534. https://doi.org/10.1093/molbev/msaa015

Muhire B. M., A. Varsani, and D. P. Martin, 2014 SDT: a virus classification tool based on pairwise sequence alignment and identity calculation. PLoS One 9: e108277. https://doi.org/10.1371/journal.pone.0108277

Nawrocki E. P., and S. R. Eddy, 2013 Infernal 1.1: 100-fold faster RNA homology searches. Bioinformatics 29: 2933–2935. https://doi.org/10.1093/bioinformatics/btt509

Neumann P., P. Novák, N. Hoštáková, and J. Macas, 2019 Systematic survey of plant LTR-retrotransposons elucidates phylogenetic relationships of their polyprotein domains and provides a reference for element classification. Mob DNA 10: 1. https://doi.org/10.1186/s13100-018-0144-1

Nickels J. F., M. E. Della-Rosa, I. Miguelez Goyeneche, S. J. Charlton, K. Sneppen, et al., 2022 The transcription factor Atf1 lowers the transition barrier for nucleosome-mediated establishment of heterochromatin. Cell Rep 39: 110828. https://doi.org/10.1016/j.celrep.2022.110828

Noad R. J., N. S. Al-Kaff, D. S. Turner, and S. N. Covey, 1998 Analysis of polypurine tract-associated DNA plus-strand priming in vivo utilizing a plant pararetroviral vector carrying redundant ectopic priming elements. J Biol Chem 273: 32568–32575. https://doi.org/10.1074/jbc.273.49.32568

Noé L., and G. Kucherov, 2005 YASS: enhancing the sensitivity of DNA similarity search. Nucleic Acids Res 33: W540–543. https://doi.org/10.1093/nar/gki478

Noma K null, C. D. Allis, and S. I. Grewal, 2001 Transitions in distinct histone H3 methylation patterns at the heterochromatin domain boundaries. Science 293: 1150–1155. https://doi.org/10.1126/science.1064150

Novikova O., G. Smyshlyaev, and A. Blinov, 2010 Evolutionary genomics revealed interkingdom distribution of Tcn1-like chromodomain-containing Gypsy LTR retrotransposons among fungi and plants. BMC Genomics 11: 231. https://doi.org/10.1186/1471-2164-11-231

Nuckolls N. L., M. A. Bravo Núñez, M. T. Eickbush, J. M. Young, J. J. Lange, et al., 2017 wtf genes are prolific dual poison-antidote meiotic drivers. Elife 6: e26033. https://doi.org/10.7554/eLife.26033

Nurk S., B. P. Walenz, A. Rhie, M. R. Vollger, G. A. Logsdon, et al., 2020 HiCanu: accurate assembly of segmental duplications, satellites, and allelic variants from high-fidelity long reads. Genome Res 30: 1291–1305. https://doi.org/10.1101/gr.263566.120

Nurk S., S. Koren, A. Rhie, M. Rautiainen, A. V. Bzikadze, et al., 2022 The complete sequence of a human genome. Science 376: 44–53. https://doi.org/10.1126/science.abj6987

Ou S., W. Su, Y. Liao, K. Chougule, J. R. A. Agda, et al., 2019 Benchmarking transposable element annotation methods for creation of a streamlined, comprehensive pipeline. Genome Biol 20: 275. https://doi.org/10.1186/s13059-019-1905-y

Paquin B., and B. F. Lang, 1996 The mitochondrial DNA of Allomyces macrogynus: the complete genomic sequence from an ancestral fungus. J Mol Biol 255: 688–701. https://doi.org/10.1006/jmbi.1996.0056

Paquin B., M. J. Laforest, L. Forget, I. Roewer, Z. Wang, et al., 1997 The fungal mitochondrial genome project: evolution of fungal mitochondrial genomes and their gene expression. Curr Genet 31: 380–395. https://doi.org/10.1007/s002940050220

Paquin B., M. J. Laforest, and B. F. Lang, 2000 Double-hairpin elements in the mitochondrial DNA of allomyces: evidence for mobility. Mol Biol Evol 17: 1760–1768. https://doi.org/10.1093/oxfordjournals.molbev.a026274

Paradis E., J. Claude, and K. Strimmer, 2004 APE: Analyses of Phylogenetics and Evolution in R language. Bioinformatics 20: 289–290. https://doi.org/10.1093/bioinformatics/btg412

Perkins V., S. Vignola, M.-H. Lessard, P.-L. Plante, J. Corbeil, et al., 2020 Phenotypic and Genetic Characterization of the Cheese Ripening Yeast Geotrichum candidum. Front Microbiol 11: 737. https://doi.org/10.3389/fmicb.2020.00737

Persson J., B. Steglich, A. Smialowska, M. Boyd, J. Bornholdt, et al., 2016 Regulating retrotransposon activity through the use of alternative transcription start sites. EMBO Rep 17: 753–768. https://doi.org/10.15252/embr.201541866

Peska V., M. Mátl, T. Mandáková, D. Vitales, P. Fajkus, et al., 2020 Human-like telomeres in Zostera marina reveal a mode of transition from the plant to the human telomeric sequences. J Exp Bot 71: 5786–5793. https://doi.org/10.1093/jxb/eraa293

Peska V., P. Fajkus, M. Bubeník, V. Brázda, N. Bohálová, et al., 2021 Extraordinary diversity of telomeres, telomerase RNAs and their template regions in Saccharomycetaceae. Sci Rep 11: 12784. https://doi.org/10.1038/s41598-021-92126-x

Pidoux A. L., and R. C. Allshire, 2004 Kinetochore and heterochromatin domains of the fission yeast centromere. Chromosome Res 12: 521–534. https://doi.org/10.1023/B:CHRO.0000036586.81775.8b

Plasterk R. H., Z. Izsvák, and Z. Ivics, 1999 Resident aliens: the Tc1/mariner superfamily of transposable elements. Trends Genet 15: 326–332. https://doi.org/10.1016/s0168-9525(99)01777-1

Pockrandt C., M. Alzamel, C. S. Iliopoulos, and K. Reinert, 2020 GenMap: ultra-fast computation of genome mappability. Bioinformatics 36: 3687–3692. https://doi.org/10.1093/bioinformatics/btaa222

Poplin R., P.-C. Chang, D. Alexander, S. Schwartz, T. Colthurst, et al., 2018 A universal SNP and small-indel variant caller using deep neural networks. Nat Biotechnol 36: 983–987. https://doi.org/10.1038/nbt.4235

Prince S., C. Munoz, F. Filion-Bienvenue, P. Rioux, M. Sarrasin, et al., 2022 Refining Mitochondrial Intron Classification With ERPIN: Identification Based on Conservation of Sequence Plus Secondary Structure Motifs. Front Microbiol 13: 866187. https://doi.org/10.3389/fmicb.2022.866187

Procházka E., F. Franko, S. Poláková, and P. Sulo, 2012 A complete sequence of Saccharomyces paradoxus mitochondrial genome that restores the respiration in S. cerevisiae. FEMS Yeast Res 12: 819–830. https://doi.org/10.1111/j.1567-1364.2012.00833.x

Quinlan A. R., and I. M. Hall, 2010 BEDTools: a flexible suite of utilities for comparing genomic features. Bioinformatics 26: 841–842. https://doi.org/10.1093/bioinformatics/btq033

Raghavan V., 2021 seqvisr

Rajaei N., K. K. Chiruvella, F. Lin, and S. U. Aström, 2014 Domesticated transposase Kat1 and its fossil imprints induce sexual differentiation in yeast. Proc Natl Acad Sci U S A 111: 15491–15496. https://doi.org/10.1073/pnas.1406027111

Rausch J. W., and S. F. J. Le Grice, 2004 “Binding, bending and bonding”: polypurine tract-primed initiation of plus-strand DNA synthesis in human immunodeficiency virus. Int J Biochem Cell Biol 36: 1752–1766. https://doi.org/10.1016/j.biocel.2004.02.016

Rausch J. W., J. T. Miller, and S. F. J. Le Grice, 2017 Reverse Transcription in the Saccharomyces cerevisiae Long-Terminal Repeat Retrotransposon Ty3. Viruses 9: E44. https://doi.org/10.3390/v9030044

Rhind N., Z. Chen, M. Yassour, D. A. Thompson, B. J. Haas, et al., 2011 Comparative functional genomics of the fission yeasts. Science 332: 930–936. https://doi.org/10.1126/science.1203357

Riley R., S. Haridas, K. H. Wolfe, M. R. Lopes, C. T. Hittinger, et al., 2016 Comparative genomics of biotechnologically important yeasts. Proc Natl Acad Sci U S A 113: 9882–9887. https://doi.org/10.1073/pnas.1603941113

Robinson J. T., H. Thorvaldsdóttir, W. Winckler, M. Guttman, E. S. Lander, et al., 2011 Integrative genomics viewer. Nat Biotechnol 29: 24–26. https://doi.org/10.1038/nbt.1754

Robson N. D., and A. Telesnitsky, 1999 Effects of 3’ untranslated region mutations on plus-strand priming during moloney murine leukemia virus replication. J Virol 73: 948–957. https://doi.org/10.1128/JVI.73.2.948-957.1999

Rutherford K. M., M. A. Harris, S. Oliferenko, and V. Wood, 2022 JaponicusDB: rapid deployment of a model organism database for an emerging model species. Genetics 220: iyab223. https://doi.org/10.1093/genetics/iyab223

Saha A., J. A. Mitchell, Y. Nishida, J. E. Hildreth, J. A. Ariberre, et al., 2015 A trans-dominant form of Gag restricts Ty1 retrotransposition and mediates copy number control. J Virol 89: 3922–3938. https://doi.org/10.1128/JVI.03060-14

Samson S., É. Lord, and V. Makarenkov, 2022 SimPlot ++: a Python application for representing sequence similarity and detecting recombination. Bioinformatics btac287. https://doi.org/10.1093/bioinformatics/btac287

Sanz-Ramos M., and J. P. Stoye, 2013 Capsid-binding retrovirus restriction factors: discovery, restriction specificity and implications for the development of novel therapeutics. J Gen Virol 94: 2587–2598. https://doi.org/10.1099/vir.0.058180-0

Sato K., M. Akiyama, and Y. Sakakibara, 2021 RNA secondary structure prediction using deep learning with thermodynamic integration. Nat Commun 12: 941. https://doi.org/10.1038/s41467-021-21194-4

Scannell D. R., and K. H. Wolfe, 2008 A burst of protein sequence evolution and a prolonged period of asymmetric evolution follow gene duplication in yeast. Genome Res 18: 137–147. https://doi.org/10.1101/gr.6341207

Sehnal D., S. Bittrich, M. Deshpande, R. Svobodová, K. Berka, et al., 2021 Mol* Viewer: modern web app for 3D visualization and analysis of large biomolecular structures. Nucleic Acids Res 49: W431–W437. https://doi.org/10.1093/nar/gkab314

Seike T., and H. Niki, 2017 Mating response and construction of heterothallic strains of the fission yeast Schizosaccharomyces octosporus. FEMS Yeast Res 17. https://doi.org/10.1093/femsyr/fox045

Shen X.-X., J. L. Steenwyk, A. L. LaBella, D. A. Opulente, X. Zhou, et al., 2020 Genome-scale phylogeny and contrasting modes of genome evolution in the fungal phylum Ascomycota. Sci Adv 6: eabd0079. https://doi.org/10.1126/sciadv.abd0079

Simão F. A., R. M. Waterhouse, P. Ioannidis, E. V. Kriventseva, and E. M. Zdobnov, 2015 BUSCO: assessing genome assembly and annotation completeness with single-copy orthologs. Bioinformatics 31: 3210–3212. https://doi.org/10.1093/bioinformatics/btv351

Smith S. A., and M. J. Donoghue, 2008 Rates of molecular evolution are linked to life history in flowering plants. Science 322: 86–89. https://doi.org/10.1126/science.1163197

Stanke M., M. Diekhans, R. Baertsch, and D. Haussler, 2008 Using native and syntenically mapped cDNA alignments to improve de novo gene finding. Bioinformatics 24: 637–644. https://doi.org/10.1093/bioinformatics/btn013

Stiebler A. C., J. Freitag, K. O. Schink, T. Stehlik, B. A. M. Tillmann, et al., 2014 Ribosomal readthrough at a short UGA stop codon context triggers dual localization of metabolic enzymes in Fungi and animals. PLoS Genet 10: e1004685. https://doi.org/10.1371/journal.pgen.1004685

Tamura K., G. Stecher, and S. Kumar, 2021 MEGA11: Molecular Evolutionary Genetics Analysis Version 11. Mol Biol Evol 38: 3022–3027. https://doi.org/10.1093/molbev/msab120

Tang H., V. Krishnakumar, J. Li, and X. Zhang, 2015 jcvi: JCVI utility libraries. Zenodo.

Tang A. D., C. M. Soulette, M. J. van Baren, K. Hart, E. Hrabeta-Robinson, et al., 2020 Full-length transcript characterization of SF3B1 mutation in chronic lymphocytic leukemia reveals downregulation of retained introns. Nat Commun 11: 1438. https://doi.org/10.1038/s41467-020-15171-6

Tao Y.-T., F. Suo, S. Tusso, Y.-K. Wang, S. Huang, et al., 2019 Intraspecific Diversity of Fission Yeast Mitochondrial Genomes. Genome Biol Evol 11: 2312–2329. https://doi.org/10.1093/gbe/evz165

Tawaramoto M. S., S.-Y. Park, Y. Tanaka, O. Nureki, H. Kurumizaka, et al., 2003 Crystal structure of the human centromere protein B (CENP-B) dimerization domain at 1.65-A resolution. J Biol Chem 278: 51454–51461. https://doi.org/10.1074/jbc.M310388200

Teysset L., V.-D. Dang, M. K. Kim, and H. L. Levin, 2003 A long terminal repeat-containing retrotransposon of Schizosaccharomyces pombe expresses a Gag-like protein that assembles into virus-like particles which mediate reverse transcription. J Virol 77: 5451–5463. https://doi.org/10.1128/jvi.77.9.5451-5463.2003

Thomas J. A., J. J. Welch, R. Lanfear, and L. Bromham, 2010 A generation time effect on the rate of molecular evolution in invertebrates. Mol Biol Evol 27: 1173–1180. https://doi.org/10.1093/molbev/msq009

Thon G., K. P. Bjerling, and I. S. Nielsen, 1999 Localization and properties of a silencing element near the mat3-M mating-type cassette of Schizosaccharomyces pombe. Genetics 151: 945–963. https://doi.org/10.1093/genetics/151.3.945

Tong P., A. L. Pidoux, N. R. T. Toda, R. Ard, H. Berger, et al., 2019 Interspecies conservation of organisation and function between nonhomologous regional centromeres. Nat Commun 10: 2343. https://doi.org/10.1038/s41467-019-09824-4

Tork S., I. Hatin, J.-P. Rousset, and C. Fabret, 2004 The major 5’ determinant in stop codon read-through involves two adjacent adenines. Nucleic Acids Res 32: 415–421. https://doi.org/10.1093/nar/gkh201

Tudor M., M. Lobocka, M. Goodell, J. Pettitt, and K. O’Hare, 1992 The pogo transposable element family of Drosophila melanogaster. Mol Gen Genet 232: 126–134. https://doi.org/10.1007/BF00299145

Tusso S., F. Suo, Y. Liang, L.-L. Du, and J. B. W. Wolf, 2022 Reactivation of transposable elements following hybridization in fission yeast. Genome Res 32: 324–336. https://doi.org/10.1101/gr.276056.121

Upadhyay U., S. Srivastava, I. Khatri, J. S. Nanda, S. Subramanian, et al., 2017 Ablation of RNA interference and retrotransposons accompany acquisition and evolution of transposases to heterochromatin protein CENPB. Mol Biol Cell 28: 1132–1146. https://doi.org/10.1091/mbc.E16-07-0485

Varadi M., S. Anyango, M. Deshpande, S. Nair, C. Natassia, et al., 2022 AlphaFold Protein Structure Database: massively expanding the structural coverage of protein-sequence space with high-accuracy models. Nucleic Acids Res 50: D439–D444. https://doi.org/10.1093/nar/gkab1061

Vaser R., I. Sović, N. Nagarajan, and M. Šikić, 2017 Fast and accurate de novo genome assembly from long uncorrected reads. Genome Res 27: 737–746. https://doi.org/10.1101/gr.214270.116

Vaughan-Martini A., and A. Martini, 2011 Chapter 66 - Schizosaccharomyces Lindner (1893), pp. 779–784 in The Yeasts (Fifth Edition), edited by Kurtzman C. P., Fell J. W., Boekhout T. Elsevier, London.

Vještica A., L. Merlini, P. J. Nkosi, and S. G. Martin, 2018 Gamete fusion triggers bipartite transcription factor assembly to block re-fertilization. Nature 560: 397–400. https://doi.org/10.1038/s41586-018-0407-5

Vyas A., A. V. Freitas, Z. A. Ralston, and Z. Tang, 2021 Fission Yeast Schizosaccharomyces pombe: A Unicellular “Micromammal” Model Organism. Curr Protoc 1: e151. https://doi.org/10.1002/cpz1.151

Walker B. J., T. Abeel, T. Shea, M. Priest, A. Abouelliel, et al., 2014 Pilon: an integrated tool for comprehensive microbial variant detection and genome assembly improvement. PLoS One 9: e112963. https://doi.org/10.1371/journal.pone.0112963

Waterhouse A. M., J. B. Procter, D. M. A. Martin, M. Clamp, and G. J. Barton, 2009 Jalview Version 2--a multiple sequence alignment editor and analysis workbench. Bioinformatics 25: 1189–1191. https://doi.org/10.1093/bioinformatics/btp033

Weaver D. C., G. V. Shpakovski, E. Caputo, H. L. Levin, and J. D. Boeke, 1993 Sequence analysis of closely related retrotransposon families from fission yeast. Gene 131: 135–139. https://doi.org/10.1016/0378-1119(93)90682-s

Wilhelm M., M. Boutabout, T. Heyman, and F. X. Wilhelm, 1999a Reverse transcription of the yeast Ty1 retrotransposon: the mode of first strand transfer is either intermolecular or intramolecular. J Mol Biol 288: 505–510. https://doi.org/10.1006/jmbi.1999.2723

Wilhelm M., T. Heyman, M. Boutabout, and F. X. Wilhelm, 1999b A sequence immediately upstream of the plus-strand primer is essential for plus-strand DNA synthesis of the Saccharomyces cerevisiae Ty1 retrotransposon. Nucleic Acids Res 27: 4547–4552. https://doi.org/10.1093/nar/27.23.4547

Wilhelm M., and F. X. Wilhelm, 2001 Reverse transcription of retroviruses and LTR retrotransposons. Cell Mol Life Sci 58: 1246–1262. https://doi.org/10.1007/PL00000937

Wood V., R. Gwilliam, M.-A. Rajandream, M. Lyne, R. Lyne, et al., 2002 The genome sequence of Schizosaccharomyces pombe. Nature 415: 871–880. https://doi.org/10.1038/nature724

Wu C. I., and W. H. Li, 1985 Evidence for higher rates of nucleotide substitution in rodents than in man. Proc Natl Acad Sci U S A 82: 1741–1745. https://doi.org/10.1073/pnas.82.6.1741

Xavier B. B., V. P. W. Miao, Z. O. Jónsson, and Ó. S. Andrésson, 2012 Mitochondrial genomes from the lichenized fungi Peltigera membranacea and Peltigera malacea: features and phylogeny. Fungal Biol 116: 802–814. https://doi.org/10.1016/j.funbio.2012.04.013

Xie Y., H. Li, X. Luo, H. Li, Q. Gao, et al., 2022 IBS 2.0: an upgraded illustrator for the visualization of biological sequences. Nucleic Acids Res gkac373. https://doi.org/10.1093/nar/gkac373

Yoon S.-H., S.-M. Ha, J. Lim, S. Kwon, and J. Chun, 2017 A large-scale evaluation of algorithms to calculate average nucleotide identity. Antonie Van Leeuwenhoek 110: 1281–1286. https://doi.org/10.1007/s10482-017-0844-4

Yoshinaka Y., I. Katoh, T. D. Copeland, and S. Oroszlan, 1985 Murine leukemia virus protease is encoded by the gag-pol gene and is synthesized through suppression of an amber termination codon. Proc Natl Acad Sci U S A 82: 1618–1622. https://doi.org/10.1073/pnas.82.6.1618

Yu C., M. J. Bonaduce, and A. J. S. Klar, 2013 Defining the epigenetic mechanism of asymmetric cell division of Schizosaccharomyces japonicus yeast. Genetics 193: 85–94. https://doi.org/10.1534/genetics.112.146233

Yue J.-X., and G. Liti, 2018 Long-read sequencing data analysis for yeasts. Nat Protoc 13: 1213–1231. https://doi.org/10.1038/nprot.2018.025

Zaratiegui M., M. W. Vaughn, D. V. Irvine, D. Goto, S. Watt, et al., 2011 CENP-B preserves genome integrity at replication forks paused by retrotransposon LTR. Nature 469: 112–115. https://doi.org/10.1038/nature09608

Zhang H., X. Zheng, and Z. Zhang, 2016 The Magnaporthe grisea species complex and plant pathogenesis. Mol Plant Pathol 17: 796–804. https://doi.org/10.1111/mpp.12342

Zhang S., and Y.-J. Zhang, 2019 Proposal of a new nomenclature for introns in protein-coding genes in fungal mitogenomes. IMA Fungus 10: 15. https://doi.org/10.1186/s43008-019-0015-5

Zhang R.-G., G.-Y. Li, X.-L. Wang, J. Dainat, Z.-X. Wang, et al., 2022 TEsorter: an accurate and fast method to classify LTR-retrotransposons in plant genomes. Hortic Res uhac017. https://doi.org/10.1093/hr/uhac017

Zhou L., T. Feng, S. Xu, F. Gao, T. T. Lam, et al., 2022 ggmsa: a visual exploration tool for multiple sequence alignment and associated data. Brief Bioinform 23: bbac222. https://doi.org/10.1093/bib/bbac222

